# Are we ready to track climate-driven shifts in marine species across international boundaries? - A global survey of scientific bottom trawl data

**DOI:** 10.1101/2020.06.18.125930

**Authors:** Aurore Maureaud, Romain Frelat, Laurène Pécuchet, Nancy Shackell, Bastien Mérigot, Malin L. Pinsky, Kofi Amador, Sean C. Anderson, Alexander Arkhipkin, Arnaud Auber, Iça Barri, Rich Bell, Jonathan Belmaker, Esther Beukhof, Mohamed Lamine Camara, Renato Guevara-Carrasco, Junghwa Choi, Helle Torp Christensen, Jason Conner, Luis A. Cubillos, Hamet Diaw Diadhiou, Dori Edelist, Margrete Emblemsvåg, Billy Ernst, Tracey P. Fairweather, Heino O. Fock, Kevin D. Friedland, Camilo B. Garcia, Didier Gascuel, Henrik Gislason, Menachem Goren, Jérôme Guitton, Didier Jouffre, Tarek Hattab, Manuel Hidalgo, Johannes N. Kathena, Ian Knuckey, Saïkou Oumar Kidé, Mariano Koen-Alonso, Matt Koopman, Vladimir Kulik, Jacqueline Palacios León, Ya’arit Levitt-Barmats, Martin Lindegren, Marcos Llope, Félix Massiot-Granier, Hicham Masski, Matthew McLean, Beyah Meissa, Laurène Mérillet, Vesselina Mihneva, Francis K.E. Nunoo, Richard O’Driscoll, Cecilia A. O’Leary, Elitsa Petrova, Jorge E. Ramos, Wahid Refes, Esther Román-Marcote, Helle Siegstad, Ignacio Sobrino, Jón Sólmundsson, Oren Sonin, Ingrid Spies, Petur Steingrund, Fabrice Stephenson, Nir Stern, Feriha Tserkova, Georges Tserpes, Evangelos Tzanatos, Itai van Rijn, Paul A.M. van Zwieten, Paraskevas Vasilakopoulos, Daniela V. Yepsen, Philippe Ziegler, James Thorson

## Abstract

Marine biota is redistributing at a rapid pace in response to climate change and shifting seascapes. While changes in fish populations and community structure threaten the sustainability of fisheries, our capacity to adapt by tracking and projecting marine species remains a challenge due to data discontinuities in biological observations, lack of data availability, and mismatch between data and real species distributions. To assess the extent of this challenge, we review the global status and accessibility of ongoing scientific bottom trawl surveys. In total, we gathered metadata for 283,925 samples from 95 surveys conducted regularly from 2001 to 2019. 59% of the metadata collected are not publicly available, highlighting that the availability of data is the most important challenge to assess species redistributions under global climate change. We further found that single surveys do not cover the full range of the main commercial demersal fish species and that an average of 18 surveys is needed to cover at least 50% of species ranges, demonstrating the importance of combining multiple surveys to evaluate species range shifts. We assess the potential for combining surveys to track transboundary species redistributions and show that differences in sampling schemes and inconsistency in sampling can be overcome with vector autoregressive spatio-temporal modeling to follow species density redistributions. In light of our global assessment, we establish a framework for improving the management and conservation of transboundary and migrating marine demersal species. We provide directions to improve data availability and encourage countries to share survey data, to assess species vulnerabilities, and to support management adaptation in a time of climate-driven ocean changes.

## Introduction

Marine species worldwide are redistributing in response to climate-induced shifting seascapes, while constrained by physiological features over latitudinal, longitudinal and bathymetric gradients^1,2^. The movement of individuals and species from one location to another in response to climate change, either through active migration or passive dispersal of early life-stages, results in colonizations and potential invasions into previously unoccupied areas^3,4^. Such redistributions have profound consequences for community composition and biodiversity, as demonstrated by substantial changes in taxonomic^5–9^ and trait diversity^10–14^. They also affect the structure and functions of marine ecosystems^15–18^. While species on the move have important socioeconomic consequences^19–22^, our capacity to adapt to these changes by tracking and projecting species range shifts across regional boundaries remains a challenge, not only scientifically, but also economically and politically^23–25^.

The capacity to detect species range and community changes in response to climate variability and change depends foremost on the ability to monitor species through, among others, the existence, coverage and quality of surveys. There is a broad variety of monitoring efforts on species distributions and abundances on land and in the oceans^26–28^. Among them, scientific bottom trawl surveys have been started in the ∼1900s and hence collected demersal marine species (living over and on the sea bottom) on continental shelves and slopes in many areas of the world^29,30^. The primary purpose of these surveys is to provide fishery-independent data to inform stock assessment of commercially important species and populations, and more recently for multidisciplinary ecosystem monitoring. Many of the surveys offer long and often continuous time-series of data on community composition and provide a unique opportunity to both track species range shifts within and across international boundaries and improve the assessment of biodiversity under global change.

Studies examining climate change impacts on marine communities across large regions have mostly focused on the North Atlantic and Northeast Pacific ecosystems^9,31–34^. This is due to the lack of availability and knowledge on the existence of bottom trawl surveys elsewhere, but also to fewer surveys and reduced transnational cooperation in the southern hemisphere compared to the northern hemisphere. While there is a global movement towards “open science”^35^, particularly by making data publicly available^36–39^, it has also sparked considerable debate on how to proceed^40,41^. Therefore, the application of open science principles, making primary outputs of scientific research available and reproducible, remains a challenge. From a broader political and management perspective, there is a need to access surveys covering species range across international boundaries to assess species responses to environmental change because there is an increasing need for transboundary assessment of commercial species. Particularly, the lack of assessment of species range shifts across international boundaries and management areas may lead to political disputes on shifting fisheries resources^24,42–44^. If the data generated by bottom trawl surveys are available—combined and modeled properly—they may allow comparisons of species distribution and abundance in time and space. This can offer the opportunity to quantitatively monitor the dynamics of species distributions and community structure by following the different stages of species redistribution and mechanisms through which communities are altered by climate and anthropogenic stressors^45^. Developing knowledge on marine species responses to climate change is the first step towards developing transboundary and international management plans.

To uncover the difficulties preventing a global assessment of marine species redistributions, we first review the existence and availability of bottom trawl surveys worldwide by collecting survey metadata. We assess the global coverage of productive and trawled seas by bottom trawl surveys. Second, we show the importance of combining surveys to cover commercial species range. Third, we demonstrate that modelling can incorporate multiple surveys and unbalanced sampling to follow species density in time and space. We propose a framework where open science would help to support transboundary management and conservation.

## Global availability of trawl survey data

Scientific bottom trawl surveys have been conducted in many countries to sample continental shelves and slopes around the world, inhabited by essential demersal fisheries resources^46^. They have formed the backbone of information supporting research on marine fish communities in response to climate change and variability across ecosystems and over large spatial scales^10,31,47–51^, as well as meta-analysis across taxonomic groups and biomes^9,34^. However, a single survey sampling demersal communities is typically carried out nationally, regionally or within a delimited management zone. The monitoring protocol might differ among surveys, and the resulting data are not always publicly available, creating an obstacle for assessing potential species range shifts. Determining the consequences of climate change on marine species critically depends on the availability and quality of the data^52,53^.

### Global data synthesis

To assess the existence and accessibility of fishery-independent data on demersal species in the global ocean, we collected metadata (latitude, longitude, and depth if available) from scientific bottom trawl surveys (henceforth surveys) including samples (hauls) of marine communities. We only collected metadata for recent and ongoing surveys that included at least one year of sampling since 2015 and use otter trawl gear, the most widely spread type of trawl to sample demersal communities. In addition, we only retained the surveys that were performed for four years or more between 2001–2019 and reported the first year surveyed if it was prior to 2001 (complete list in Supplementary Table 1.1). Finally, we excluded surveys covering near-shore areas (conducted within 3 miles from the coast) as they are designed to primarily target juvenile or coastal fish. However, we kept track of all surveys that did not follow our selection criteria (Supplementary Table 2.1).

The survey information was gathered and built on knowledge of previously existing survey collections (https://datras.ices.dk/, https://oceanadapt.rutgers.edu/, https://james-thorson.shinyapps.io/FishViz/) and previous studies using aggregated survey data^29,31,47–51^. In order to ensure a broad geographical coverage, we sent a standardized query to established and identified contacts of surveys, particularly where geographical gaps were identified (South America, Africa, Asia, and Oceania). In case no obvious contact person was available, we contacted national fisheries institutes. We acknowledge that despite the rigorous surveying and querying, some surveys might still be missing.

For each survey, based on haul coordinates, we estimated the spatial area covered using an alpha-convex hull method^54^, where we set the parameter controlling for the volume shape at *α* = 1. This allowed the creation of polygons from the location of samples at the extremes of the surveyed area and a rough estimation of the area covered by surveys. Overall, available metadata covered 95 surveys across 78 Exclusive Economic Zones (EEZs) around the world and constitute a total of 283,925 unique geo-referenced hauls from 2001 to 2019 (Figure 1, Supplementary Table 1.1), covering approximately 2,509,000 km^2^ in total.

**Figure 1:**
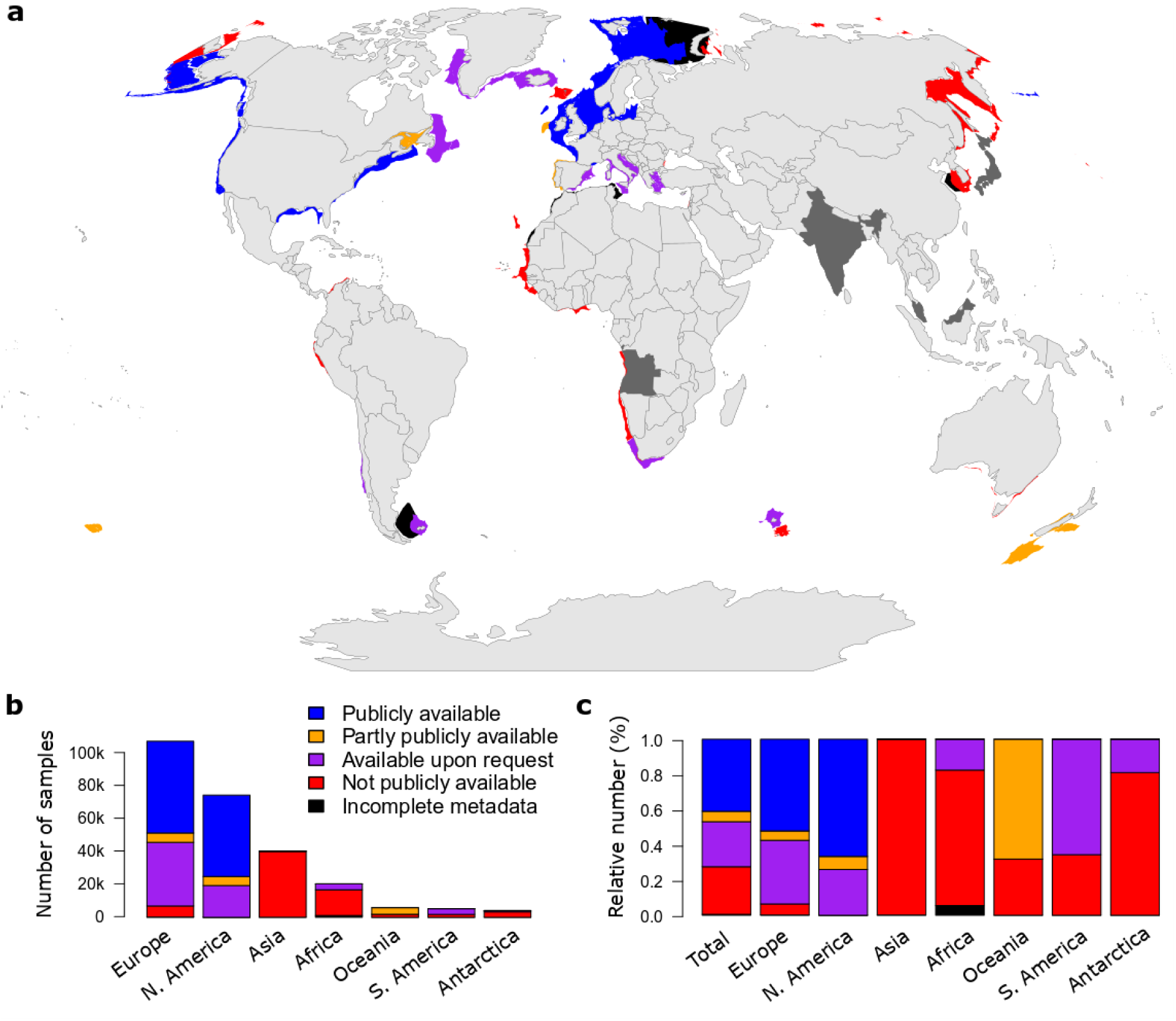
Worldwide availability of bottom trawl surveys, classified as follows: *publicly available* (blue), *partly publicly available* (orange), *available upon request* (purple), *not publicly available* (red), *incomplete metadata* (black) and *unavailable metadata* (dark grey for countries conducting the survey). **a, l**ocation of ongoing scientific bottom trawl surveys, represented by the survey convex hull. Surveys are classified according to their availability. Additional visualizations are available in Supplementary 4. **b,** number of samples for 2001-019 and availability across continents. (c) Relative availability of samples across continents.

### Surveys cover productive and fished continental shelves

We compared the spatial extent of the surveys with the area covered by productive continental shelves and the area fished by bottom trawlers. To estimate the area of productive continental shelves, we used global depth data (GEBCO https://www.gebco.net/) and chlorophyll-a concentration data (GlobColour GSM, 2005–2015)^55^. The fished area was estimated from bottom trawl fishing data from Global Fishing Watch for the years 2013 to 2016^56^. The environmental and fisheries data were aggregated on a grid (0.04°×0.04°). A grid cell was considered as a productive shelf site if its depth was between 30-500 m and its chlorophyll-a concentration was higher than 0.5 mg/m^3^ (several thresholds tested are shown in Supplementary 3). A grid cell was counted as fished if more than one trawling activity was detected in the grid cell in the period 2013-2016. Thus, a grid cell could be designated as productive, fished, neither, or both. We then compared the productive and fished grid cells with the convex hull of surveys to compute the global proportion of productive shelves and fished areas covered by the surveys. We estimated that the surveys cover 62% of continental productive seas and 54% of coastal bottom-trawl fished areas (Supplementary Figure 3.3 and Figure 3.5).

### Criteria for data accessibility

The survey metadata were classified based on their relative degree of accessibility, using the following classification criteria:

- *Publicly available*: data for all years and species sampled were available in a public repository
- *Partly publicly available*: data only for some years, or only for some species were available in a public repository and access to full data is possible upon request
- *Available upon request*: data are not publicly available but access to data is possible upon request. This category was assigned if at least one person not affiliated to the institution that owns the data obtained the full raw data via request
- *Not publicly available*: when the data, to the best of our knowledge, were not publicly available, access to data is not possible upon request, but access to metadata is possible upon request
- *Incomplete metadata*: when the data were not publicly available and we received access to partial survey metadata via request, or were reconstructed from the literature
- *Unavailable metadata*: when we are aware of ongoing surveys but did not receive access to the metadata, and/or were unable to reconstruct the metadata from the literature

### Global status of availability

Among all collected surveys, species abundance/biomass data from 41% of the survey hauls are *publicly available*, while an additional 31% of the surveys *partly publicly available* or *available upon request* (Figure 1). The remaining 28% of the surveys are classified as *not publicly available* or have *incomplete metadata* and are therefore not available. While species range shifts in response to climate change have occurred across a broad range of aquatic organisms worldwide^2,21,34^, most marine studies are concentrated in the Northern hemisphere with a majority of surveys located in the North Atlantic and Northeast Pacific. This can be explained by the geographical coverage of surveys in the Southern hemisphere, which is considerably more restricted and includes almost exclusively not publicly available data, except for South Africa, Chile, New Zealand, Falkland Islands and Kerguelen Islands (classified as *partly publicly available* and *available upon request*, Figure 1**a,b**). Lower transnational collaboration within Regional Fisheries Management Organizations (RMFOs) in the southern hemisphere may explain this difference in availability. While our international review of the coverage and accessibility of scientific surveys shows that surveys are regularly conducted across continental shelf seas worldwide (78 EEZs), a vast majority of the publicly available data are located in Europe and North America (Figure 1**c**).

### Need for improving data availability

The dominance of Northern hemisphere climate change studies has been specifically criticized^32,34,57,58^. The under-representation of tropical seas, polar areas and southern hemisphere studies may mislead our knowledge of demersal communities’ response to global change. The non-availability of data can be, among many other factors, driven by lack of human and/or logistical resources and capacity to maintain data management systems, or by institutional incentives controlling data access. Furthermore, the difficulty and inability to obtain even metadata from established contacts of current, known surveys, might illustrate that the location of sampling is considered as sensitive, likely from a political and economic perspective. We provide here the most exhaustive assessment of ongoing bottom trawl surveys metadata around the world, and provide information on who owns the data and where it can be requested, aiming at enhancing data sharing (Box. 1).

#### Box 1. Applying open science principles to bottom trawl survey data.

Open Science is broadly defined as “Open data and content that can be freely used, modified, and shared by anyone for any [ethical] purpose” (http://opendefinition.org/), and is more specifically described by six main principles (see ref ^39^ for a general description). Following is a summary of advances towards Open Science and challenges regarding the use of bottom trawl surveys.

##### 1. Ensure ethical use of shared information

It is crucial that the push towards open science recognizes the value and human side of information. Nations and communities, particularly those that have been historically exploited must be able to benefit from their own data and be able to control their own information to minimize potential abuse. Open science must ensure that open data do not enable an opportunistic fishing company to exploit a nation’s or community’s resources. While open science can promote transparent science and understanding, it is essential that any use of open data give priority, proper credit, acknowledgement and potentially compensation to those who collected the information, paid for collection and recognize the nation where the data were collected. Access to data from economically stressed nations may require some type of compensation to ensure the data continue to be collected and made available.

##### 2. Improve knowledge on existing trawl surveys (“Open resources”)

Knowledge about the existence of a survey and about the essential course of actions to request and access the survey data can be a challenge. This could be facilitated through a network or a platform where scientists can share such relevant information. Regional platforms currently exist in some areas: western Africa (http://www.projet-istam.org/; http://pescao-demerstem.org/), southeast Asia^30^, Europe (https://datras.ices.dk/) and North America (https://oceanadapt.rutgers.edu). Such platforms would ideally improve the visibility of their resources by making their metadata available and easily visible. Here, we established a global network for open resources regarding bottom trawl surveys, where metadata of surveys and contacts or links to access full data are provided (Supplementary Table 1.1 and https://rfrelat.shinyapps.io/metabts/). The difficulty in obtaining survey metadata and accessing it suggests that challenges remain to create an exhaustive global resource for bottom trawl surveys that can be maintained on the medium and long term.

##### 3. Improve the accessibility and availability of surveys (“Open data”)

An evaluation of the accessibility and availability of surveys is necessary to enhance further open data science. We assessed that 59% of the samples collected are not publicly available to varying degrees (Figure 1). The network created here greatly improves the visibility of surveys, by presenting their metadata. Further, the availability of data can only be improved by changing the way we share scientific information, for instance by publishing data^52^ and ensuring quality-controlled use of data. Several bottom trawl surveys are published online, but the most recent years are not always included, or links to access the data are not always maintained, e.g. the Norwegian surveys^107^, the southern Gulf of St Lawrence survey^108^, Mauritania^109^, southeast Asian surveys^30^. Ensuring online publication of data are updated and maintained is key, as is done for other repositories (e.g. DATRAS from the International Council for the Exploration of the Sea https://datras.ices.dk/). Existing platforms that enable online data publication, however, may not always allow updating or involve peer-review of the data (e.g. PANGAEA https://www.pangaea.de/ and DRYAD https://datadryad.org/stash). To ensure data are available beyond a single report or publication, a dedicated, sustainable, long-term management strategy is required with dedicated personal. The data repositories mentioned above that update and maintain their information all have dedicated programs and resources to ensure the data are available.

##### 4. Improve the visibility of the expertise on surveys (“Open source” and “Open methods”)

To ensure appropriate use of the data, it is highly important that survey protocols, reports, and common practices are shared together with the raw survey data. Furthermore, providing example code to clean the data or to combine data from multiple surveys can ensure the appropriate use of data. Such types of open source and open access methods have been developed in recent years, mostly for Europe and North America (for instance https://oceanadapt.rutgers.edu/; https://james-thorson.github.io// for codes; https://datras.ices.dk/ and ref ^72^ for codes and published reports). However, such documentation and tools need to be available and easy to find beyond Europe and North America, as well as in multiple languages. Together with the survey metadata information, we started gathering such information (see Supplementary Table 1.1).

##### 5. Open access of the produced research (“Open access”, “Open peer review”)

The research published based on bottom trawl surveys can be open access, even when the raw underlying data are not (see ref ^47^ for an example). The accessibility of published papers in science is improving, but is undermined by journals charging high costs for open access, which is particularly prejudicing open publications in low and middle-income countries. Open peer review is developing in several journals and will also increase transparency towards publishing. Similarly, providing programming code and the underlying data are increasingly required for publishing, thereby also enhancing open science.

There are many documented cases where disagreements regarding fishing rights have led to serious international conflicts^44,59^. Improved science regarding range shifts across regional boundaries, their impacts on fisheries and fishing communities^22,43,60^, as well as political and regulation landscapes^19,61,62^, could lead to better planning for contingencies regarding climate-driven distribution shifts^24^. This would provide scientific information to design adaptation and management measures that anticipate potential international conflicts. We therefore argue that financial or political incentives should be identified to better share the existing data, and develop good data management systems^63^. However, benefits from sharing data are diffuse, while their costs (in terms of lost publication opportunities for local teams) are concentrated^63^, and this leads to the well-known “concentrated-diffuse’’ mechanism for policy failure^64^. This type of policy failure can be partly overcome by concentrating scattered incentives, either by providing multilateral forums where many scientists can jointly benefit from data sharing (e.g. North Pacific Marine Science Organization, https://meetings.pices.int/, International Council for the Exploration of the Sea, https://www.ices.dk/, RFMOs) or by bilateral data-sharing agreements^65^. The movement towards publicly available and accessible data in science can lead to a lack of recognition of the source, devaluation of essential investments such as data collection, preparation and curation^63^. As a result, it remains hard to enhance public availability of data. Publishing data, following FAIR principles (Findable, Accessible, Interoperable and Reusable, https://www.go-fair.org/fair-principles/) as well as open science principles, could ultimately increase the visibility of surveys but will face the challenges of thoughtfully use the data by prioritizing the need of and give credit to the data providers^52^ (Box. 1).

## Species range covered by surveys

Studies quantifying species range shifts often assume that surveys are representative of species’ native range^3,4,66,67^ when in fact species’ ranges may extend well beyond the monitored area. Demersal fish habitats are often only partially covered by surveys, particularly since surveys are designed to sample soft bottoms on primarily shallow continental shelves, hence excluding hard bottoms and reefs. In addition, most of the surveys are limited by depth, sampling the continental shelves but rarely the slopes at greater depths. Moreover, ecosystems beyond national jurisdiction are often excluded. To assess the percentage of species range covered by current surveys, and to evaluate the probability of species range shifts to occur beyond surveyed areas, we compared the habitat from species distribution models to the areas covered by the surveys.

To estimate the extent to which existing surveys cover species distribution range, we selected the top three demersal species with the highest commercial catch in each of twelve main FAO fishing areas from FishStats^68^ (a platform reporting annual fisheries catch per fishing zone, http://www.fao.org/fishery/statistics/software/fishstat/), defined as the average catch over 2001–2019. We ended up studying 37 demersal species (with some species covering several FAO areas). For the commercial species identified, we downloaded the native range from AquaMaps^69^, which shows the probability of occurrence of each species on a 0.5°×0.5° grid. We used the modeled native range and considered it as the “true” habitat and species range. The habitat in AquaMaps is based on publicly available global occurrence data and expert judgment on species environmental niches. Even though AquaMaps may sometimes misrepresent species ranges because of poor occurrence data and lack of knowledge for some species^70^, the ranges of the selected commercial species are generally well documented. The preferred habitat data layer of each commercial species for the analysis was defined as all locations from the AquaMaps habitat maps where the probability of occurrence was higher than 0.5 (more details in Supplementary 5).

Next, we calculated the percentage of grid cells from the AquaMaps commercial species habitat area covered by the survey footprints, showing the overlap between the two across all FAO fishing areas. We also included the availability status of surveys. We demonstrate that no combination of available surveys covers the entire range of any single species (Figure 2). Nevertheless, for about a quarter of the species considered, existing surveys cover more than 50% of the species habitats, up to a maximum of 79% for Atlantic cod (*Gadus morhua*). However, even for these well-surveyed species, the surveys are sometimes not available (MVO, *Lophius vomerinus* and HKK, *Merluccius capensis* in Southeast Atlantic and South Africa) or only a part are *publicly available* (PCO, *Gadus macrocephalus* and ALK, *Gadus chalcogrammus* in the North Pacific). We computed the number of surveys that overlapped with the species native range and show that the number needed to cover at least 50% of the main commercial species habitat is highly variable (from 4 to 31, average of 18, Figure 2**b**) and depends on the areas covered by each survey. The restricted spatial extent of some surveys conducted in Europe explains why more than 15 surveys may be needed to cover 50% of a specific species range, emphasizing the need for standardizing surveys and developing tools to combine data from different surveys.

**Figure 2:**
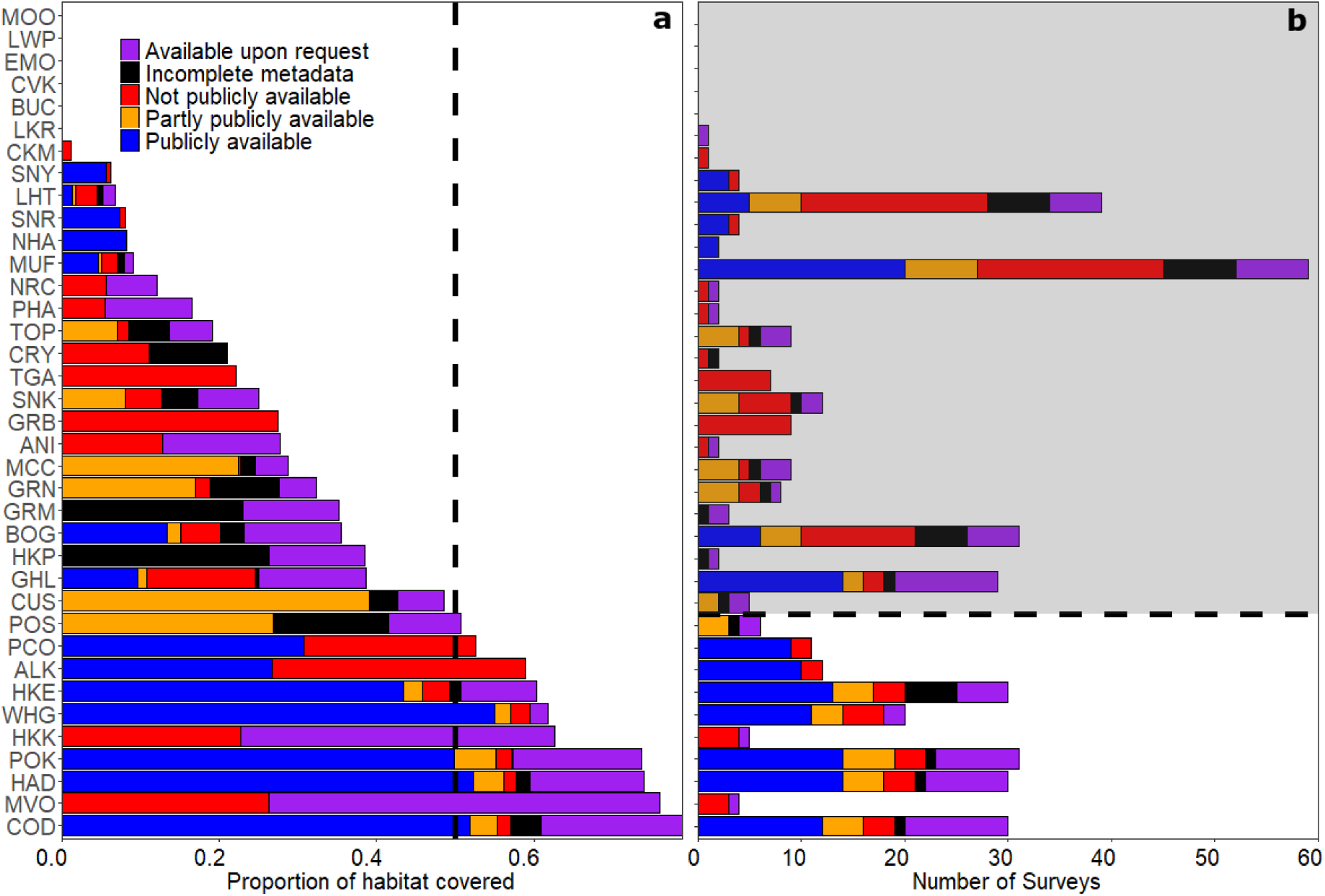
Main commercial demersal species identified by the ASFIS 3-letter codes and the corresponding coverage by the surveys: **a,** proportion of AquaMaps habitat covered by the surveys (the vertical dotted line indicates 50% of range covered) and **b,** number of surveys behind the proportion covered (species for which less than 50% of range covered are shaded). Corresponding Latin names to species are available in Supplementary Table 5.1. Colors indicate the availability status attributed to each survey.

## Tracking species densities across management areas

We have shown that demersal fish ranges and habitats are not fully covered by bottom trawl surveys, which may be particularly problematic when fisheries stocks and populations are transboundary. The capacity to track such transboundary species throughout their range critically depends on the ability to combine surveys from multiple sources and regions.

In cases where data are available and gathered from different sources, formatting, language differences and lack of user expertise on the survey itself may limit the ability to use the data appropriately. For instance, information on units, haul duration or swept area estimates are sometimes lacking, limiting the combined use of multiple independent surveys. In addition, differences in gear, sampling designs, species identification procedures and catchability across and within surveys may bias perceptions of species distribution and regional changes in abundance^71^. In order to standardize processing of such data, we recommend improving the availability of survey documentation, including explanations of survey methodology and associated coding that can be freely applied to clean, standardize, and combine surveys (Box. 1). Making expert knowledge easily accessible will facilitate studies combining multiple surveys^72^.

### A case study to combine surveys across regions

We used arrowtooth flounder (*Atheresthes stomias*) to illustrate how to combine survey data across multiple regions when tracking and investigating population-scale range shifts in species distribution. Arrowtooth flounder is a widespread and ecologically important predator in the Northeast Pacific^73^, monitored and assessed by 10 distinct but contiguous surveys across the region from the California Current to the Bering Sea between 2001 and 2018. To predict densities within the entire survey domain, we fit a spatio-temporal Poisson-link delta-gamma model^74^ to biomass data from each survey using the R-package VAST^71,75^ (vector autoregressive spatio-temporal). This model has the advantage of interpolating density across time and space within the survey domain when survey data are lacking in a given area or time step (see Supplementary 6). We assumed that each survey has identical gear performance (i.e. catches the same proportion of individuals within the area-swept by bottom trawl gear). The validated model shows that the highest densities of arrowtooth flounder are observed in the center of distribution within the Gulf of Alaska (Figure 3). However, densities have recently increased in the eastern Bering Sea and the distribution has shifted inshore and northward. Simultaneously, its distribution has slightly moved southward in the California Current. Despite this expansion at both ends of its range, the centroid of the population shows a net change northward by 40 km in less than 20 years (Figure 3**c**).

**Figure 3:**
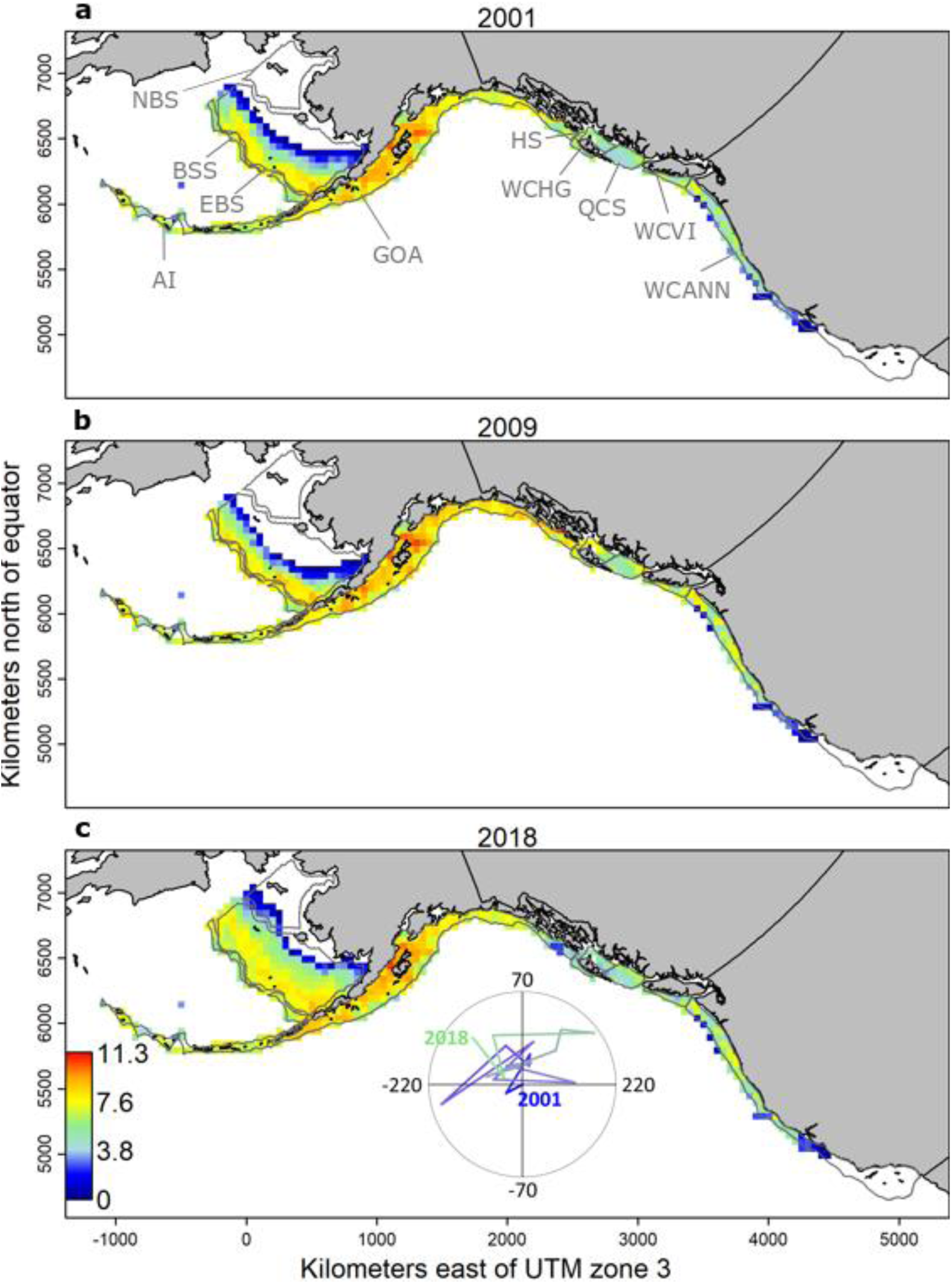
Density estimates for arrowtooth flounder (*Atheresthes stomias*) along the northeastern Pacific coast containing contiguous sampling data from multiple surveys in log(kg/km^2^) using regional bottom trawl surveys: West Coast US (WCANN), West Coast Vancouver Island (WCVI), Hecate Strait (HS), West Coast Haida Gwaii (WCHG), Queen Charlotte (WCS), Gulf of Alaska (GOA), eastern Bering Sea (EBS), northern Bering Sea (NBS), Aleutian Islands (AI), and Bering Sea slope (BSS). Polygon contours represent the different surveys, as indicated by the corresponding codes. Densities are presented for three years: **a** 2001, **b** 2009 and **c** 2018. Only densities higher than 0.1% of the maximum were selected to clearly differentiate areas occupied by the species (colored-coded) from those mostly unoccupied (white). The inset in the bottom panel shows the change in the center of gravity through time, where longitude and latitude for 2001 are (0;0).

### Improving the management of transboundary species

The use and coordination of multiple surveys is needed to monitor commercially important species distributions that extend beyond a singular survey. Our case study and other recent studies^76–79^ show that statistical tools to reconstruct species densities across surveys is possible, and can appropriately quantify the changes of densities through time and across regions by correcting for unbalanced sampling^71,75^. The multiple-survey approach—applied here to the arrowtooth flounder—is applicable for many other wide-ranging species, including commercially important transboundary species such as Greenland halibut^80^, Pacific cod^81^ and Atlantic cod^67,82^. For each of these species, combining surveys will require initial research to determine the most appropriate methods to account for differences in catchability between surveys, species and sites^75,83–85^. For example, this could be done by using regression-discontinuity-designs to estimate catchability ratios for surveys that are contiguous, but not overlapping. Furthermore, ongoing efforts to standardize national surveys will also help to combine surveys. For instance, Russia and Norway started a joint ecosystem-wide survey in the Barents Sea instead of conducting them separately^86^. This is also the case of the MEDITS program, ensuring consistent sampling protocol across EU regions of the Mediterranean Sea^87^. Such intercalibration of different survey schemes and sampling designs will improve long-term intercomparison of surveys and will help assess marine species range shifts under global warming. Our knowledge of species redistribution across surveys will clearly benefit from long-term consistent surveys, when combined and modeled appropriately.

Long-term global monitoring datasets are essential to develop transboundary science, and offer opportunities to improve the management and conservation of migrating transboundary species (Box. 2). Under global change, migrating species create a potential for economic and political conflicts^88^, and may lead to species overexploitation or collapse in the case of lack of adaptation and cooperation^24,89–92^. The 1982 United Nations Convention on the Law of the Sea (UNCLOS) provides the legal framework for international obligations towards safeguarding marine resources. Migratory and transboundary stocks are principally managed by RFMOs^92–94^, or conservation-related initiatives such as the Global Transboundary Conservation Network (http://www.tbpa.net/). Still, these organizations need to explicitly consider governance in the context of climate change^90,94,95^. While building a common understanding of status and trends is a key first step towards transboundary cooperation^24^, international governance will require global geopolitical flexibility and the establishment of transnational agreements^96,97^, which need to be supported by cross-boundary open science (Box. 1 and Box. 2). The adaptation of management and policy is essential for the sustainability and conservation of shared resources, but are often motivated by economic and cultural values rather than ecological considerations^96^. Scientifically-supported transboundary governance will avoid overexploitation, conflicts about newly or historically shared resources and conservation of vulnerable species. The identification of priority areas, vulnerable transboundary species and capacity for adaptation is essential to initiate fast and efficient governance of ‘biodiversity beyond national jurisdictions’^98–100^.

#### Box 2. Towards a framework for transboundary management and conservation

##### 1. Agreement of Survey Data and Analyses

All parties must agree on data, information, current abundance and spatial footprint of the fisheries stock and species of interest.

###### a. Find the surveys covering the species range

The first key task is to establish which surveys cover the native range of the species of interest. The next task is to find and access surveys corresponding to that species native range. A list of existing surveys, their time coverage and available samples are available here for demersal species (see Supplementary Table 1.1). If surveys are not publicly available, one can use the network of contacts published in this paper and/or establish bilateral/multilateral agreements to gain access to the survey data. Metadata should also ideally include a list of species recorded in the surveys.

###### b. Estimate the change in species density and distribution

Surveys vary in terms of design, gear, catchability and sampling methods. Multiple surveys can be combined to estimate species density and reconstruct past temporal changes in spatial distribution. Modeling the change in species distributions can be done with multiple models and needs to take into account survey discrepancies^76–79^. Sharing information across international boundaries could enable a more complete picture of the distribution of a fish population and reconsideration of the definition of their stock structure.

###### c. Forecast changes in densities and distributions

Building on the knowledge of past species distribution change and ecology, forecasting species distribution will enable adaptation to changes in advance^110^. The spatio-temporal vmodel described here can also be used to forecast species distributions in the near future^111^, and thus the spatial scale at which adaptation measures should be applied.

###### d. Measure ecosystem impact in local management areas

Climate change enhances the dynamic nature of changes in species abundance and requires fisheries management to adapt, not only directly to the resource, but also to assess the impacts on port infrastructure, fishing fleets/gears and other human activities^20,22,112^. The immigration/emigration of species into local areas can lead to substantial changes in community structure and diversity, and may lead other species to outcompete or be outcompeted. By monitoring not only commercial species, but the entire community—as is generally possible with bottom trawl surveys—we can understand ecological changes and inform the conservation of vulnerable species^17,96^.

##### 2. Management and cooperation

###### a. International agreements

All parties must create a management agreement for the regulation of the resource that is legally binding, regardless of how the distribution or abundance of the resource might change or not change in the future. Policies could be developed within the agreement to adjust regulations depending on a range of future scenarios such as when stocks move poleward, or decrease/increase in abundance. Pre-agreements covering a range of options can help reduce future conflicts and reduce the need to renegotiate or abandon the agreement^113^. An important goal of the management agreement is to acknowledge that changes are likely to occur, while recognizing that the specific change is likely unpredictable.

###### b. Transboundary cooperation

In the case of transboundary species and distribution over multiple management areas, changes in spatial distribution under climate change and variability may favor/exclude countries or regions^90,114^. Therefore, some areas will ‘win’ or ‘lose’ and create conflicts and/or lead to species overexploitation^24,88^. Building agreements among countries to share resources equitably—or compensate when not possible—is necessary to ensure the sustainability of resources and the dependent human communities^92^. It is essential that all parties perceive benefit from cooperating and remaining within the agreement. In the case of non-exploited/non-targeted species, cooperative conservation actions should be established. Such cooperation can only be built with open and transparent science (Box. 1) to conserve the species and avoid conflicts.

###### c. Regulation and enforcement

To truly implement transboundary management and conservation, the involved parties must develop a method to enforce their agreement. Effective monitoring to gather information on the shared resource and to measure compliance is important. Compensation and/or penalties may also be involved to ensure all parties adhere to the regulations. Side payments are one means of compensation that can take different forms^92^. Direct cash payments are possible, however countries can also share monitoring and research capacity across international boundaries as is done for a number of stocks that straddle Canadian and United States waters^92^. Nations can allow other nations to fish for a specific shared resource in their EEZ as is done between Norway and Russia^115,116^ or swap quota in a multispecies fishery as has been done in the Baltic^117^. Once again, the goal is to develop an agreement in which all parties perceive benefits to properly manage shared stocks.

## Maintaining surveys to face future challenges in the oceans

Regular bottom trawl surveys do not cover the entire global ocean nor the whole continental shelves and the lack of monitoring makes it problematic to track ecosystem change and adapt management and policy to shifting resources^23^. A range of alternative approaches could be considered to better cover demersal species habitats, understand changes in spatial distribution under global change, and promote an ecosystem approach to fisheries and other human activities management. Such alternatives could include both citizen science initiatives reporting species observed well outside their known ranges (e.g. the Range Extension Database and Mapping project; Redmap^101^ https://www.redmap.org.au/ and the European Alien Species Information; EASIN^102^ https://easin.jrc.ec.europa.eu/easin) as well as formal fishery-dependent data (such as observer, landings, vessel trip report data) able to report species occurrences and abundances. Other promising sources of data could be derived from environmental DNA^103^. Datasets derived from such alternative sources can improve evidence of range shifts, particularly in poorly sampled areas.

Bottom trawl surveys are highly valuable for following marine species redistribution and biodiversity change. However, maintaining surveys in a consistent way through time is a challenge as they are costly and require political agreements. However, ecological time series become more informative the longer their timespan, highlighting the need to maintain long-term monitoring programs^53,104^. The existence of international programs such as the Nansen program (http://www.fao.org/in-action/eaf-nansen/en/) is valuable to inform ecosystem-based management^105,106^ and could be expanded, assuming that the data collected become available. Bottom trawl surveys impact seafloor habitat, benthic communities, and sampled fish^29^ and we should ensure that this kind of monitoring benefits science as much as possible. Surveys provide relevant and essential information for fisheries management and marine ecology, but they must be designed to be as efficient as possible by sharing (meta)data, providing opportunities for innovative uses of the data and improve the economic and ecological efficiency of monitoring. In any case, challenges of sampling marine communities and sharing data need to be overcome to allow scientific assessment and adequate management of shifting marine resources.

## Conclusions

The concentration of marine studies in the Northern hemisphere profoundly limits not only our ability to track and understand climate change effects and species range shifts, but also our capacity to adapt, mitigate or avoid potential conflicts and socio-economic consequences that follow. This is particularly important in parts of the developing world where fisheries constitute a primary source of food and livelihood for coastal communities, but information supporting management is often scarce or non-existing. To alleviate these issues, a coherent framework to monitor, understand and inform sound and scientifically underpinned management actions to adapt to species range shifts is needed. Our study provides a first step towards creating such a framework by conducting a joint and internationally coordinated synthesis of the global coverage and availability of survey data, and it will be of great assistance to various users aiming to assess and predict the response of marine biodiversity to climate change. Our study has identified important gaps in data availability and accessibility, and suggested ways to make the best use of surveys at hand by combining data from multiple sources to assess species redistributions over multiple management areas. In order to support comparative studies on species range shifts, we stress the need for nations to strengthen monitoring efforts, particularly in under sampled areas of the world. Moreover, we make a general plea for open science as well as fair and transparent sharing of data. This is needed to support scientific advice on coordinated spatial management actions, allowing us to adapt and prepare for the inevitable ecological and socio-economic consequences of climate change yet to come.

## Acknowledgments

We thank all persons who have collected the survey data around the world. We are also grateful to everyone who answered our emails and provided contacts, expertise and guidance on survey existence and status: Abdul Wahab Abdullah, Daniela Alemany, Valérie Allain, Denis Bernier, Arnaud Bertrand, Philippe Bouchet, José Miguel Casas, George Daskalov, Andrey V. Dolgov, Bernadine Everett, Louise Flensborg, Bertrand Richer de Forges, Elizabeth Fulton, Carolina Giraldo, Donald R. Kobayashi, Teoh Shwu Jiau, Michel Kulbicki, Marc Léopold, Rich Little, Franck Magron, David Mills, Sten Munch-Petersen, J. Rasmus Nielsen, Daniel Pauly, Gretta Pecl, Cohen Philippa, Raul Primicerio, Romeo Saldívar-Lucio, Nicolas Rolland, Sarah Samadi, Jörn Schmidt, Kwang-Tsao Shao, Andrew Smith, Alex Tilley, Stephanie Turner, Vincent Vallée, Francisco Velasco, Laurent Vigliola and Manuel J. Zetina-Rejón. We thank Inês Dias Bernardes for providing the bottom trawl survey metadata from the Nansen Program. We thank the International Council for the Exploration of the Sea (ICES) for providing the space to organize side-meetings during the Annual Science Conference (Göteborg, Sweden, 2019), at the origin of this project. Aurore Maureaud received funding from Villum research grant awarded to Martin Lindegren (No. 13159) and conducted the work within the Centre for Ocean Life, a Villum Kann Rasmussen Center of Excellence supported by the Villum Foundation. We further thank National Institute of Fisheries Science (R2020021) for providing the Korean trawl survey data. The scientific results and conclusions, as well as any views or opinions expressed herein, are those of the author(s) and do not necessarily reflect those of NOAA or the Department of Commerce. The EU NAFO data used in this paper have been funded by the EU through the European Maritime and Fisheries Fund (EMFF) within the National Program of collection, management and use of data in the fisheries sector and support for scientific advice regarding the Common Fisheries Policy.

## Author contributions

A.M., R.F., L.P., N.S. and J.T. designed the project, directed and performed the data collection and analyses. A.M. was leading the project and R.F. was leading the data curation. J.T. produced the code and ran the model to estimate density of species with multiple surveys. R.F., J.T. and A.M. produced figures and conducted analyses. All authors have either provided metadata, provided contact lists and/or lists of surveys to help collecting the metadata, conducted metadata requests and/or helped with the interpretation of results. A.M. wrote the first draft of the manuscript and all co-authors were given the opportunity to revise it.

## References

1. Pinsky, M. L., Eikeset, A. M., McCauley, D. J., Payne, J. L. & Sunday, J. M. Greater vulnerability to warming of marine versus terrestrial ectotherms. Nature 569, 108–111 (2019).

2. Poloczanska, E. S. et al. Global imprint of climate change on marine life. Nat. Clim. Change 3, 919–925 (2013).

3. Dulvy, N. K. et al. Climate change and deepening of the North Sea fish assemblage: a biotic indicator of warming seas. J. Appl. Ecol. 45, 1029–1039 (2008).

4. Perry, A. L. Climate Change and Distribution Shifts in Marine Fishes. Science 308, 1912–1915 (2005).

5. Batt, R. D., Morley, J. W., Selden, R. L., Tingley, M. W. & Pinsky, M. L. Gradual changes in range size accompany long-term trends in species richness. Ecol. Lett. 20, 1148–1157 (2017).

6. Fossheim, M. et al. Recent warming leads to a rapid borealization of fish communities in the Arctic. Nat. Clim. Change 5, 673–677 (2015).

7. Magurran, A. E., Dornelas, M., Moyes, F., Gotelli, N. J. & McGill, B. Rapid biotic homogenization of marine fish assemblages. Nat. Commun. 6, 8405 (2015).

8. Hiddink, J. G. & ter Hofstede, R. Climate induced increases in species richness of marine fishes. Glob. Change Biol. 14, 453–460 (2008).

9. Antão, L. H. et al. Temperature-related biodiversity change across temperate marine and terrestrial systems. Nat. Ecol. Evol. 1–7 (2020) doi: 10.1038/s41559-020-1185-7.

10. Burrows, M. T. et al. Ocean community warming responses explained by thermal affinities and temperature gradients. Nat. Clim. Change 9, 959–963 (2019).

11. Dencker, T. S. et al. Temporal and spatial differences between taxonomic and trait biodiversity in a large marine ecosystem: Causes and consequences. PLOS ONE 12, e0189731 (2017).

12. McLean, M. et al. A Climate-Driven Functional Inversion of Connected Marine Ecosystems. Curr. Biol. 28, 3654-3660.e3 (2018).

13. McLean, M. J., Mouillot, D., Goascoz, N., Schlaich, I. & Auber, A. Functional reorganization of marine fish nurseries under climate warming. Glob. Change Biol. 25, 660–674 (2019).

14. Frainer, A. et al. Climate-driven changes in functional biogeography of Arctic marine fish communities. Proc. Natl. Acad. Sci. 114, 12202–12207 (2017).

15. Friedland, K. D. et al. Changes in higher trophic level productivity, diversity and niche space in a rapidly warming continental shelf ecosystem. Sci. Total Environ. 704, 135270 (2020).

16. Kortsch, S., Primicerio, R., Fossheim, M., Dolgov, A. V. & Aschan, M. Climate change alters the structure of arctic marine food webs due to poleward shifts of boreal generalists. Proc. R. Soc. B Biol. Sci. 282, 20151546 (2015).

17. Pinsky, M. L., Selden, R. L. & Kitchel, Z. J. Climate-Driven Shifts in Marine Species Ranges: Scaling from Organisms to Communities. Annu. Rev. Mar. Sci. 12, 153–179 (2020).

18. Mérillet, L., Kopp, D., Robert, M., Mouchet, M. & Pavoine, S. Environment outweighs the effects of fishing in regulating demersal community structure in an exploited marine ecosystem. Glob. Change Biol. 26, 2106–2119 (2020).

19. Dubik, B. A. et al. Governing fisheries in the face of change: Social responses to long-term geographic shifts in a U.S. fishery. Mar. Policy 99, 243–251 (2019).

20. Greenan, B. J. W. et al. Climate Change Vulnerability of American Lobster Fishing Communities in Atlantic Canada. Front. Mar. Sci. 6, (2019).

21. Pecl, G. T. et al. Biodiversity redistribution under climate change: Impacts on ecosystems and human well-being. Science 355, (2017).

22. Pinsky, M. L. & Fogarty, M. Lagged social-ecological responses to climate and range shifts in fisheries. Clim. Change 115, 883–891 (2012).

23. Lindegren, M. & Brander, K. Adapting Fisheries and Their Management To Climate Change: A Review of Concepts, Tools, Frameworks, and Current Progress Toward Implementation. Rev. Fish. Sci. Aquac. 26, 400–415 (2018).

24. Pinsky, M. L. et al. Preparing ocean governance for species on the move. Science 360, 1189–1191 (2018).

25. Ramesh, N., Rising, J. A. & Oremus, K. L. The small world of global marine fisheries: The cross-boundary consequences of larval dispersal. Science 364, 1192–1196 (2019).

26. Dornelas, M. et al. BioTIME: A database of biodiversity time series for the Anthropocene. Glob. Ecol. Biogeogr. 27, 760–786 (2018).

27. Blowes, S. A. et al. The geography of biodiversity change in marine and terrestrial assemblages. Science 366, 339–345 (2019).

28. Edgar, G. J. et al. Abundance and local-scale processes contribute to multi-phyla gradients in global marine diversity. Sci. Adv. 3, e1700419 (2017).

29. Trenkel, V. M. et al. We can reduce the impact of scientific trawling on marine ecosystems. Mar. Ecol. Prog. Ser. 609, 277–282 (2019).

30. Garces, L. R. et al. A regional database management system—the fisheries resource information system and tools (FiRST): Its design, utility and future directions. Fish. Res. 78, 119–129 (2006).

31. Pinsky, M. L., Worm, B., Fogarty, M. J., Sarmiento, J. L. & Levin, S. A. Marine Taxa Track Local Climate Velocities. Science 341, 1239–1242 (2013).

32. Richardson, A. J. et al. Climate change and marine life. Biol. Lett. 8, 907–909 (2012).

33. Shackell, N. L., Ricard, D. & Stortini, C. Thermal Habitat Index of Many Northwest Atlantic Temperate Species Stays Neutral under Warming Projected for 2030 but Changes Radically by 2060. PLOS ONE 9, e90662 (2014).

34. Lenoir, J. et al. Species better track the shifting isotherms in the oceans than on lands. http://biorxiv.org/lookup/doi/10.1101/765776 (2019) doi: 10.1101/765776.

35. OCDE. Making Open Science a Reality. (2015) doi: https://doi.org/10.1787/5jrs2f963zs1-en.

36. Nosek, B. A. et al. Promoting an open research culture. Science 348, 1422–1425 (2015).

37. Cheruvelil, K. S. & Soranno, P. A. Data-Intensive Ecological Research Is Catalyzed by Open Science and Team Science. BioScience 68, 813–822 (2018).

38. Poisot, T., Bruneau, A., Gonzalez, A., Gravel, D. & Peres-Neto, P. Ecological Data Should Not Be So Hard to Find and Reuse. Trends Ecol. Evol. 34, 494–496 (2019).

39. Gallagher, R. V. et al. Open Science principles for accelerating trait-based science across the Tree of Life. Nat. Ecol. Evol. 1–10 (2020) doi: 10.1038/s41559-020-1109-6.

40. Moles, A., Dickie, J. & Flores-Moreno, H. A response to Poisot et al.: Publishing your dataset is not always virtuous. Ideas Ecol. Evol. 6, (2013).

41. Poisot, T., Mounce, R. & Gravel, D. Moving toward a sustainable ecological science: don’t let data go to waste! Ideas Ecol. Evol. 6, (2013).

42. Baudron, A. R. et al. Changing fish distributions challenge the effective management of European fisheries. Ecography 42, 1–12 (2020).

43. Shackell, N. L., Frank, K. T., Nye, J. A. & den Heyer, C. E. A transboundary dilemma: dichotomous designations of Atlantic halibut status in the Northwest Atlantic. ICES J. Mar. Sci. 73, 1798–1805 (2016).

44. Spijkers, J. & Boonstra, W. J. Environmental change and social conflict: the northeast Atlantic mackerel dispute. Reg. Environ. Change 17, 1835–1851 (2017).

45. Bates, A. E. et al. Defining and observing stages of climate-mediated range shifts in marine systems. Glob. Environ. Change 26, 27–38 (2014).

46. Meeting the sustainable development goals. (2018).

47. Beukhof, E. et al. Marine fish traits follow fast-slow continuum across oceans. Sci. Rep. 9, 1–9 (2019).

48. Branch, T. A. et al. The trophic fingerprint of marine fisheries. Nature 468, 431–435 (2010).

49. Pecuchet, L. et al. From traits to life-history strategies: Deconstructing fish community composition across European seas. Glob. Ecol. Biogeogr. 26, 812–822 (2017).

50. Shin, Y.-J., Rochet, M.-J., Jennings, S., Field, J. G. & Gislason, H. Using size-based indicators to evaluate the ecosystem effects of fishing. ICES J. Mar. Sci. 62, 384–396 (2005).

51. Gislason, H. et al. Species richness in North Atlantic fish: Process concealed by pattern. Glob. Ecol. Biogeogr. geb.13068 (2020) doi: 10.1111/geb.13068.

52. Costello, M. J., Michener, W. K., Gahegan, M., Zhang, Z.-Q. & Bourne, P. E. Biodiversity data should be published, cited, and peer reviewed. Trends Ecol. Evol. 28, 454–461 (2013).

53. Schindler, D. E. & Hilborn, R. Prediction, precaution, and policy under global change. Science 347, 953–954 (2015).

54. Pateiro López, B. & Rodriguez-Casal, A. alphahull: Generalization of the Convex Hull of a Sample of Points in the Plane. R package version 2.2. (2019).

55. Maritorena, S., d’Andon, O. H. F., Mangin, A. & Siegel, D. A. Merged satellite ocean color data products using a bio-optical model: Characteristics, benefits and issues. Remote Sens. Environ. 114, 1791–1804 (2010).

56. Kroodsma, D. A. et al. Tracking the global footprint of fisheries. Science 359, 904–908 (2018).

57. Lenoir, J. & Svenning, J.-C. Climate-related range shifts – a global multidimensional synthesis and new research directions. Ecography 38, 15–28 (2015).

58. Feeley, K. J., Stroud, J. T. & Perez, T. M. Most ‘global’ reviews of species’ responses to climate change are not truly global. Divers. Distrib. 23, 231–234 (2017).

59. GuÐmundsson, G. J. The Cod and the Cold War. Scand. J. Hist. 31, 97–118 (2006).

60. Young, T. et al. Adaptation strategies of coastal fishing communities as species shift poleward. ICES J. Mar. Sci. 76, 93–103 (2019).

61. Cheung, W. W. L., Pinnegar, J., Merino, G., Jones, M. C. & Barange, M. Review of climate change impacts on marine fisheries in the UK and Ireland: CLIMATE CHANGE AND MARINE FISHERIES IN IRELAND AND THE UK. Aquat. Conserv. Mar. Freshw. Ecosyst. 22, 368–388 (2012).

62. McIlgorm, A. et al. How will climate change alter fishery governance? Insights from seven international case studies. Mar. Policy 34, 170–177 (2010).

63. Fenichel, E. P. & Skelly, D. K. Why Should Data Be Free; Don’t You Get What You Pay For? BioScience 65, 541–542 (2015).

64. Shanahan, E. A., Jones, M. D., McBeth, M. K. & Lane, R. R. An Angel on the Wind: How Heroic Policy Narratives Shape Policy Realities: Narrative Policy Framework. Policy Stud. J. 41, 453–483 (2013).

65. Hammer, M. & Hoel, A. H. The Development of Scientific Cooperation under the Norway–Russia Fisheries Regime in the Barents Sea. Arct. Rev. 3, (2012).

66. Albouy, C., Guilhaumon, F., Araújo, M. B., Mouillot, D. & Leprieur, F. Combining projected changes in species richness and composition reveals climate change impacts on coastal Mediterranean fish assemblages. Glob. Change Biol. 18, 2995–3003 (2012).

67. Morley, J. W. et al. Projecting shifts in thermal habitat for 686 species on the North American continental shelf. PLOS ONE 13, e0196127 (2018).

68. FAO. Fisheries and aquaculture software. FishStat Plus - Universal software for fishery statistical time series. http://www.fao.org/fishery/meta:FI:citation/topic/16073/en?url=http://www.fao.org/fishery/statistics/software/fishstat/en (2017).

69. Kaschner, K. et al. AquaMaps: Predicted range maps for aquatic species. www.aquamaps.org (2016).

70. O’Hara, C. C., Afflerbach, J. C., Scarborough, C., Kaschner, K. & Halpern, B. S. Aligning marine species range data to better serve science and conservation. PLOS ONE 12, e0175739 (2017).

71. Thorson, J. T. & Barnett, L. A. K. Comparing estimates of abundance trends and distribution shifts using single- and multispecies models of fishes and biogenic habitat. ICES J. Mar. Sci. 74, 1311–1321 (2017).

72. Moriarty, M., Greenstreet, Simon. P. R. & Rasmussen, J. Derivation of Groundfish Survey Monitoring and Assessment Data Products for the Northeast Atlantic Area. Scott. Mar. Freshw. Sci. 8, 240 pp. (2017).

73. Aydin, K. & Mueter, F. The Bering Sea—A dynamic food web perspective. Deep Sea Res. Part II Top. Stud. Oceanogr. 54, 2501–2525 (2007).

74. Thorson, J. T. Three problems with the conventional delta-model for biomass sampling data, and a computationally efficient alternative. Can. J. Fish. Aquat. Sci. 75, 1369–1382 (2017).

75. Thorson, J. T. Guidance for decisions using the Vector Autoregressive Spatio-Temporal (VAST) package in stock, ecosystem, habitat and climate assessments. Fish. Res. 210, 143–161 (2019).

76. Thorson, J. T., Pinsky, M. L. & Ward, E. J. Model-based inference for estimating shifts in species distribution, area occupied and centre of gravity. Methods Ecol. Evol. 7, 990–1002 (2016).

77. Selden, R. L. et al. Coupled changes in biomass and distribution drive trends in availability of fish stocks to US West Coast ports. ICES J. Mar. Sci. 77, 188–199 (2020).

78. Dolder, P. J., Thorson, J. T. & Minto, C. Spatial separation of catches in highly mixed fisheries. Sci. Rep. 8, 1–11 (2018).

79. Ono, K., Ianelli, J. N., McGilliard, C. R. & Punt, A. E. Integrating data from multiple surveys and accounting for spatio-temporal correlation to index the abundance of juvenile Pacific halibut in Alaska. ICES J. Mar. Sci. 75, 572–584 (2018).

80. Wheeland, L. J. & Morgan, M. J. Age-specific shifts in Greenland halibut (Reinhardtius hippoglossoides) distribution in response to changing ocean climate. ICES J. Mar. Sci. fsz152 (2019) doi: 10.1093/icesjms/fsz152.

81. Drinkwater, K. F. The response of Atlantic cod (Gadus morhua) to future climate change. ICES J. Mar. Sci. 62, 1327–1337 (2005).

82. Spies, I. et al. Genetic evidence of a northward range expansion in the eastern Bering Sea stock of Pacific cod. Evol. Appl. 13, 362–375 (2020).

83. Moriarty, M. et al. Combining fisheries surveys to inform marine species distribution modelling. ICES J. Mar. Sci. (2020) doi: 10.1093/icesjms/fsz254.

84. Fraser, H. M., Greenstreet, S. P. R. & Piet, G. J. Taking account of catchability in groundfish survey trawls: implications for estimating demersal fish biomass. ICES J. Mar. Sci. 64, 1800–1819 (2007).

85. Walker, N. D., Maxwell, D. L., Le Quesne, W. J. F. & Jennings, S. Estimating efficiency of survey and commercial trawl gears from comparisons of catch-ratios. ICES J. Mar. Sci. J. Cons. fsw250 (2017) doi: 10.1093/icesjms/fsw250.

86. Eriksen, E. et al. From single species surveys towards monitoring of the Barents Sea ecosystem. Prog. Oceanogr. 166, 4–14 (2018).

87. Spedicato, M. T. et al. The MEDITS trawl survey specifications in an ecosystem approach to fishery management. Sci. Mar. 83, 9–20 (2020).

88. Mendenhall, E. et al. Climate change increases the risk of fisheries conflict. Mar. Policy 117, 103954 (2020).

89. Pershing, A. J. et al. Slow adaptation in the face of rapid warming leads to collapse of the Gulf of Maine cod fishery. Science 350, 809–812 (2015).

90. Oremus, K. L. et al. Governance challenges for tropical nations losing fish species due to climate change. Nat. Sustain. 1–4 (2020) doi: 10.1038/s41893-020-0476-y.

91. Vosooghi, S. Panic-Based Overfishing in Transboundary Fisheries. Environ. Resour. Econ. 73, 1287–1313 (2019).

92. Miller, K. A. & Munro, G. R. Climate and Cooperation: A New Perspective on the Management of Shared Fish Stocks. Mar. Resour. Econ. 19, 367–393 (2004).

93. Aqorau, T., Bell, J. & Kittinger, J. N. Good governance for migratory species. Science 361, 1208–1209 (2018).

94. VanderZwaag, D. L., Bailey, M. & Shackell, N. L. Canada–U.S. Fisheries Management in the Gulf of Maine: Taking Stock and Charting Future Coordinates in the Face of Climate Change. Ocean Yearb. Online 31, 1–26 (2017).

95. Pentz, B., Klenk, N., Ogle, S. & Fisher, J. A. D. Can regional fisheries management organizations (RFMOs) manage resources effectively during climate change? Mar. Policy 92, 13–20 (2018).

96. Scheffers, B. R. & Pecl, G. Persecuting, protecting or ignoring biodiversity under climate change. Nat. Clim. Change 9, 581–586 (2019).

97. Miller, K. A., Munro, G. R., Sumaila, U. R. & Cheung, W. W. L. Governing Marine Fisheries in a Changing Climate: A Game-Theoretic Perspective: GOVERNING MARINE FISHERIES IN A CHANGING CLIMATE. Can. J. Agric. Econ. Can. Agroeconomie 61, 309–334 (2013).

98. Mason, N., Ward, M., Watson, J. E. M., Venter, O. & Runting, R. K. Global opportunities and challenges for transboundary conservation. Nat. Ecol. Evol. 1–8 (2020) doi: 10.1038/s41559-020-1160-3.

99. United Nations General Assembly. Development of an international legally binding instrument under the United Nations Convention on the Law of the Sea on the conservation and sustainable use of marine biological diversity of areas beyond national jurisdiction. A/RES/69/292. https://www.un.org/ga/search/view_doc.asp?symbol=A/RES/69/292#x0026;Lang=E (2015).

100. Fletcher, R. et al. Biodiversity Beyond National Jurisdiction: Legal options for a new international agreement. (2017).

101. Pecl, G. T. et al. Redmap Australia: Challenges and Successes With a Large-Scale Citizen Science-Based Approach to Ecological Monitoring and Community Engagement on Climate Change. Front. Mar. Sci. 6, 349 (2019).

102. Schade, S. et al. Aliens in Europe. An open approach to involve more people in invasive species detection. Comput. Environ. Urban Syst. 78, 101384 (2019).

103. Salter, I., Joensen, M., Kristiansen, R., Steingrund, P. & Vestergaard, P. Environmental DNA concentrations are correlated with regional biomass of Atlantic cod in oceanic waters. Commun. Biol. 2, 1–9 (2019).

104. Hughes, B. B. et al. Long-Term Studies Contribute Disproportionately to Ecology and Policy. BioScience 67, 271–281 (2017).

105. Bianchi, G. Impact of fishing on size composition and diversity of demersal fish communities. ICES J. Mar. Sci. 57, 558–571 (2000).

106. Bianchi, G. et al. Collaboration between the Nansen Programme and the Large Marine Ecosystem Programmes. Environ. Dev. 17, 340–348 (2016).

107. Djupevåg, O. IMR bottom trawl data 1980-2017. (2018) doi: 10.21335/NMDC-1657305299.

108. Swain, D. P., Benoît, H. P. & Hammill, M. O. Data from: Spatial distribution of fishes in a Northwest Atlantic ecosystem in relation to risk of predation by a marine mammal. 2620101 bytes (2016) doi: 10.5061/DRYAD.N43QF.

109. Kidé, S. O., Manté, C., Dubroca, L., Demarcq, H. & Mérigot, B. Spatio-Temporal Dynamics of Exploited Groundfish Species Assemblages Faced to Environmental and Fishing Forcings: Insights from the Mauritanian Exclusive Economic Zone. PLOS ONE 10, e0141566 (2015).

110. Payne, M. R. et al. Lessons from the First Generation of Marine Ecological Forecast Products. Front. Mar. Sci. 4, (2017).

111. Thorson, J. T. Forecast skill for predicting distribution shifts: A retrospective experiment for marine fishes in the Eastern Bering Sea. Fish Fish. 20, 159–173 (2019).

112. Young, T. et al. Adaptation strategies of coastal fishing communities as species shift poleward. ICES J. Mar. Sci. 76, 93–103 (2019).

113. Hilborn, R., Maguire, J.-J., Parma, A. M. & Rosenberg, A. A. The Precautionary Approach and risk management: can they increase the probability of successes in fishery management? Can. J. Fish. Aquat. Sci. 58, 99–107 (2001).

114. Golden, C. D. et al. Nutrition: Fall in fish catch threatens human health. Nature 534, 317–320 (2016).

115. FAO. Report of the Norway-FAO Expert Consultation on the Management of Shared Fish Stocks. (2002).

116. Hønneland, G. Norway and Russia: Bargaining Precautionary Fisheries Management in the Barents Sea. Arct. Rev. 5, (2014).

117. Ranke, W. Co-operative Fisheries Management Issues in the Baltic Sea. 123–32 http://www.fao.org/3/y4652e/y4652e0a.htm (2003).

