## Supplementary for "Are we ready to track climate-driven shifts in marine species across international boundaries? - A global survey of scientific bottom trawl data"

This document contains the supplementary material (additional methods and analyses) associated to the following article.

### Supplementary 1: Metadata collection

**Table 1.1**: Summary of the collection of scientific bottom trawl surveys organized per continent. Surveys are classified according to their availability: publicly available (PA, blue), partly publicly available (PPA, orange), available upon request (AUR, purple), not publicly available (NPA, red), incomplete metadata (IM, black) and unavailable metadata (UM, grey). The table specifies the total number of samples, temporal coverage and depth range in meters, with N/A indicating lacking information, information on the data providers. The column 'Length' indicates if length measurements of individuals are available.

| No. | Survey | Continent | Area | Number of hauls | Years | Number of years | First year of sampling | Depth range | Area covered (km²) | Provider/link to access | Availability | Length | Technical description |
|---|---|---|---|---|---|---|---|---|---|---|---|---|---|
| 1 | AI | North America | Aleutian Islands | 3282 | 2002-2018 | 8 | 1983 | 22-488 | 72,300 | NOAA access | PA | | NOAA link |
| 2 | BSS | North America | Bering Sea Slope | 1136 | 2002-2016 | 6 | 2002 | 202-1200 | 36,000 | NOAA access, Jerry Hoff | PA | AUR | NOAA link |
| 3 | DFO-HS | North America | Hecate Strait | 1134 | 2005-2017 | 7 | 2005 | 18-340 | 18,000 | DFO, access | PA | PA | DFO link |
| 4 | DFO-NF | North America | Newfoundland | 14446 | 2001-2019 | 18 | 1995 | 36-1494 | 462,000 | DFO, Mariano Koen-Alonso | AUR | AUR | |
| 5 | DFO-QCS | North America | Queen Charlotte Sound | 2145 | 2003-2017 | 9 | 2003 | 44-574 | 29,000 | DFO, access | PA | PA | DFO link |
| 6 | DFO-WCHG | North America | West Coast Haida Gwaii | 764 | 2006-2016 | 7 | 2006 | 161-1328 | 5,900 | DFO, access | PA | PA | DFO link |
| 7 | DFO-WCVI | North America | West Coast Vancouver Island | 986 | 2004-2016 | 7 | 2004 | 43-803 | 12,800 | DFO, access | PA | PA | DFO link |
| 8 | EBS | North America | Eastern Bering Sea | 6761 | 2001-2018 | 18 | 1982 | 19-207 | 432,800 | NOAA access, Lyle Britt | PA | AUR | NOAA link |
| 9 | GOA | North America | Gulf of Alaska | 6299 | 2001-2017 | 9 | 1984 | 11-514 | 313,900 | NOAA access, Wayne Palsson | PA | AUR | NOAA link |
| 10 | GMEX | North America | Gulf of Mexico | 2653 | 2008-2017 | 10 | 1982 | 8-195 | 167,100 | Gulf States Marine Fisheries and Ocean Adapt | PA | | |

| | | | | | | | | | | | | | |
|---|---|---|---|---|---|---|---|---|---|---|---|---|---|
| 11 | GSL-N | North America | Gulf of St. Lawrence North | 3477 | 2001-2019 | 5 | 1990 | 38-524 | 138,100 | DFO, access 2004-2019, access 1990-2005 | PA | PA | DFO link 2004-2019, DFO link 1990-2005 |
| 12 | GSL-S | North America | Gulf of St. Lawrence South | 2598 | 2001-2017 | 14 | 1971 | 14-378 | 80,000 | DFO access and Nicolas Rolland | PPA | AUR | |
| 13 | LEJANAS -FLEM | North America | Flemish Cap | 3250 | 2001-2019 | 18 | | 80-1462 | 52,500 | Esther Róman-Marcote | AUR | AUR | |
| 14 | LEJANAS -3L | North America | Newfoundland NAFO area 3L | 1454 | 2003-2019 | 15 | | 104-1478 | 21,800 | Esther Róman-Marcote | AUR | AUR | |
| 15 | LEJANAS -3NO | North America | Newfoundland NAFO area 3NO | 2182 | 2001-2019 | 18 | | 36-1666 | 33,200 | Esther Róman-Marcote | AUR | AUR | |
| 16 | NBS | North America | Northern Bering Sea | 286 | 2010-2017 | 2 | 1982 | 11-79 | 162,600 | NOAA access, Lyle Britt | PA | AUR | NOAA link |
| 17 | NEUS | North America | Northeast US | 9216 | 2001-2018 | 18 | 1963 | 14-422 | 239,800 | OceanAdapt | PA | PA | |
| 18 | SCS | North America | Scotian Shelf | 3298 | 2001-2018 | 18 | 1970 | 0-282 | 195,100 | OceanAdapt, Donald Clark | PA | AUR | |
| 19 | SEUS | North America | Southeast US | 5422 | 2001-2018 | 18 | 1989 | 2-13 | 11,000 | OceanAdapt | PA | | |
| 20 | WCANN | North America | West Coast Annual | 11302 | 2003-2018 | 16 | 2003 | 24-1428 | 156,300 | OceanAdapt, Aimee Keller | PA | AUR | NOAA link |
| 21 | BAS-NR | Europe | Barents Sea, Norwegian data | 17895 | 2001-2019 | 18 | 1980 | 29-3395 | 1,054,000 | IMR access | PPA | PPA | AUR |
| 22 | BAS-RU | Europe | Barents Sea, Russian data | N/A | N/A | N/A | N/A | N/A | N/A | Andrey V. Dolgov | IM | N/A | N/A |
| 23 | BITS | Europe | Baltic Sea | 9713 | 2001-2019 | 18 | 1991 | 3-190 | 183,200 | ICES access | PA | PA | ICES link |
| 24 | CGFS | Europe | English Channel | 1444 | 2001-2016 | 16 | 1988 | 8-72 | 27,300 | ICES access | PA | PA | ICES link SIH link |
| 25 | EVHOE | Europe | Bay of Biscay | 2488 | 2001-2018 | 18 | 1997 | 13-587 | 204,400 | ICES access | PA | PA | ICES link SIH link |

| | | | | | | | | | | | | | |
|---|---|---|---|---|---|---|---|---|---|---|---|---|---|
| 26 | FAR-BANK | Europe | Faroe Islands | 946 | 2001-2018 | 18 | 1983 | 96-636 | 5,000 | Petur Steingrund | NPA | NPA | |
| 27 | FAR-DEEP | Europe | Faroe Islands | 424 | 2014-2019 | 6 | 2014 | 144-1058 | 7,700 | Petur Steingrund | NPA | NPA | |
| 28 | FAR-SHELF | Europe | Faroe Islands | 5686 | 2001-2019 | 19 | 1983 | 59-530 | 37,000 | Petur Steingrund | NPA | NPA | |
| 29 | GRL-DE | Europe | Greenland | 1528 | 2001-2017 | 17 | 1981 | 0-880 | 51,400 | Heino O. Fock | AUR | AUR | |
| 30 | GRL-GHLE | Europe | Greenland | 1077 | 2001-2017 | 7 | | 153-1481 | 39,500 | Helle Siegstad | AUR | AUR | |
| 31 | GRL-GHLW | Europe | Greenland | 1282 | 2001-2019 | 10 | | 202-1493 | 75,700 | Helle Siegstad | AUR | AUR | |
| 32 | GRL-SFE | Europe | Greenland | 966 | 2007-2017 | 5 | | 139-655 | 75,400 | Helle Siegstad | AUR | AUR | |
| 33 | GRL-SFW | Europe | Greenland | 4646 | 2001-2019 | 19 | | 43-800 | 194,000 | Helle Siegstad | AUR | AUR | |
| 34 | ICE-GFS | Europe | Iceland | 16470 | 2001-2019 | 19 | 1985 | 19-1420 | 283,300 | Jón Sólmundsson | AUR | AUR | https://www.hafogvatn.is/static/research/files/hv2020-08.pdf |
| 35 | IE-IGFS | Europe | Irish Sea | 2452 | 2003-2018 | 16 | 2003 | 10-741 | 197,000 | ICES access | PA | PA | ICES link |
| 36 | MEDITS-COR | Europe | Corsica | 339 | 2001-2016 | 15 | 1994 | 64-572 | 1,700 | IFREMER access and Georges Tserpes | PA | PA | SIH link |
| 37 | MEDITS-CRO | Europe | Croatia | 892 | 2002-2016 | 15 | 1996 | 3-703 | 36,800 | Georges Tserpes | AUR | AUR | MEDITS website |
| 38 | MEDITS-CYP | Europe | Cyprus | 285 | 2005-2016 | 11 | 2006 | 11-848 | 4,600 | Georges Tserpes | AUR | AUR | MEDITS website |
| 39 | MEDITS-GOL | Europe | Gulf of Lions | 1027 | 2001-2016 | 16 | 1994 | 22-866 | 11,300 | IFREMER access and Georges Tserpes | PA | PA | SIH link |
| 40 | MEDITS-GRC | Europe | Greece | 1501 | 2001-2016 | 16 | 1994 | 12-877 | 195,300 | Georges Tserpes | AUR | AUR | MEDITS website |

| | | | | | | | | | | | | | |
|---|---|---|---|---|---|---|---|---|---|---|---|---|---|
| 41 | MEDITS-ITA | Europe | Italia | 10912 | 2001-2016 | 16 | 1994 | 1-964 | 351,900 | Georges Tserpes | AUR | AUR | MEDITS website |
| 42 | MEDITS-MLT | Europe | Malta | 538 | 2005-2016 | 13 | 2000 | 45-1513 | 29,200 | Georges Tserpes | AUR | AUR | MEDITS website |
| 43 | MEDITS-SVN | Europe | Slovenia | 32 | 2001-2016 | 16 | 1996 | 22-25 | 12 | Georges Tserpes | AUR | AUR | MEDITS website |
| 44 | MEDITS-ESP | Europe | Spain | 2630 | 2001-2016 | 16 | 1994 | 22-2689 | 116,700 | Georges Tserpes | AUR | AUR | MEDITS website |
| 45 | NIGFS | Europe | Northern Ireland | 1249 | 2005-2018 | 14 | 2005 | 10-99 | 33,000 | ICES access | PA | PA | ICES link |
| 46 | NOR-BTS | Europe | Norwegian Sea | 8342 | 2001-2019 | 19 | 1980 | 18-1385 | 606,300 | IMR access | PPA | PPA | AUR |
| 47 | NS-IBTS | Europe | North Sea | 13698 | 2001-2019 | 19 | 1965 | 10-415 | 512,600 | ICES access | PA | PA | ICES link |
| 48 | PT-IBTS | Europe | Portugal | 1289 | 2002-2017 | 15 | 2002 | 19-705 | 31,100 | ICES access | PA | PA | |
| 49 | ROCKALL | Europe | Rockall Plateau | 323 | 2001-2009 | 8 | 1999 | 140-236 | 6,800 | ICES access | PA | PA | ICES link |
| 50 | SCOROC | Europe | Rockall Plateau | 331 | 2011-2018 | 8 | 2011 | 122-460 | 21,000 | ICES access | PA | PA | ICES link |
| 51 | SCOWCGFS | Europe | Scotland Shelf Sea | 993 | 2011-2019 | 8 | 2011 | 40-500 | 113,200 | ICES access | PA | PA | ICES link |
| 52 | SP-ARSA | Europe | Gulf of Cadiz | 1180 | 2001-2018 | 18 | 1996 | 19-770 | 7,400 | ICES access, Marcos Llope | PPA | PPA | |
| 53 | SP-NORTH | Europe | North of Spain | 2373 | 2001-2018 | 18 | 1990 | 71-468 | 21,400 | ICES access, Marcos Llope | PPA | PPA | |
| 54 | SP-PORC | Europe | Porcupine Survey (Scotland) | 1460 | 2001-2018 | 18 | 2001 | 187-787 | 47,800 | ICES access, Marcos Llope | PPA | PPA | |
| 55 | SWC-IBTS | Europe | Scotland Shelf Sea | 1322 | 2001-2010 | 10 | 1985 | 10-500 | 113,500 | ICES access | PA | PA | ICES link |
| 56 | WBLS | Europe | Western Black Sea | 345 | 2014-2019 | 5 | 1955 | 20-88 | 4,000 | Elitsa Petrova, Feriha Tserkova & Vesselina Mihneva | NPA | NPA | |

| | | | | | | | | | | | |
|---|---|---|---|---|---|---|---|---|---|---|---|
| 57 | ALG | Africa | Algeria | 229 | 2001-2006 | 6 | 2000 | 22-928 | 9,900 | Wahid Refes | IM |
| 58 | GIN | Africa | Guinea | 954 | 2002-2017 | 10 | | 4-560 | 47,300 | Mohammed Lamine Camara | NPA |
| 59 | GNB | Africa | Guinee Bissau | 504 | 2002-2019 | 5 | | 5-713 | 18,500 | Içа Barri | NPA |
| 60 | MEDITS-MAR | Africa | Morocco – Mediterranean ecosystem | 53 | 2006 | 1 | | 23-787 | 8,500 | Hicham Masski | IM |
| 61 | MAR-NORTH | Africa | Morocco – Northern Mud Shelf | 82 | 2010 | 1 | | 20-3618 | 26,500 | Hicham Masski | IM |
| 62 | MAR-SOUTH | Africa | Morocco – Saharan bank | 446 | 2000-2004 | 4 | | 6-105 | 35,600 | Hicham Masski | IM |
| 63 | MRT | Africa | Mauritania | 4319 | 2001-2016 | 17 | 1982 | 4-816 | 72,600 | Meissa Beyah | NPA |
| 64 | NAM | Africa | Namibia | 3250 | 2001-2018 | 16 | | 82-785 | 124,900 | John Kathena | NPA |
| 65 | NAM-NANSEN | Africa | Nansen Namibia | 574 | 2002-2019 | 18 | | 132-659 | 114,600 | | NPA |
| 66 | AGO-NANSEN | Africa | Nansen Angola | 3145 | 2001-2019 | 18 | | 23-733 | 56,400 | | NPA |
| 67 | COD-NANSEN | Africa | Nansen Democratic Rep. Congo | 78 | 2002-2016 | 8 | | 31-744 | 600 | | NPA |
| 68 | GHA-NANSEN | Africa | Nansen Ghana | 574 | 2002-2019 | 10 | | 21-579 | 52,300 | Kofi Amador and | NPA |
| 69 | GIN-NANSEN | Africa | Nansen Guinea | 147 | 2006-2019 | 10 | | 26-507 | 31,800 | | NPA |

| | | | | | | | | | | | | | |
|---|---|---|---|---|---|---|---|---|---|---|---|---|---|
| 70 | GNB-NANSEN | Africa | Nansen Guinea-Bissau | 117 | 2006-2019 | 10 | | 25-535 | 17,000 | | NPA | | |
| 71 | CIV-NANSEN | Africa | Nansen Ivory Coast | 224 | 2002-2019 | 6 | | 22-95 | 7,000 | | NPA | | |
| 72 | ZAF | Africa | South Africa | 5297 | 2001-2019 | 17 | | 18-997 | 217,900 | Tracey P. Fairweather | AUR | AUR | |
| 73 | ZAF-NANSEN | Africa | Nansen South Africa | 1834 | 2001-2019 | 11 | | 86-661 | 181,500 | | NPA | | |
| 74 | SEN | Africa | Senegal | 1577 | 2001-2016 | 10 | | 9-3001 | 184,600 | Hamet Diaw Diadhiou | NPA | | |
| 75 | SEN-NANSEN | Africa | Nansen Senegal | 193 | 2001-2017 | 5 | | 14-833 | 24,600 | | NPA | | |
| 76 | TUN | Africa | Tunisia | 424 | 2001-2006 | 6 | | N/A | 68,700 | Tarek Hattab | IM | | |
| 77 | AUS-GABTS | Oceania | Great Australian Bight Trawl Sector | 679 | 2005-2018 | 8 | 2005 | 85-220 | 7,300 | Ian Knuckey & Matt Koopman | NPA | NPA | |
| 78 | AUS-SESSF | Oceania | Southern and Eastern Australia | 1373 | 2008-2016 | 5 | 2008 | 36-989 | 37,000 | Ian Knuckey & Matt Koopman | NPA | NPA | |
| 79 | NZ-CHAT | Oceania | New-Zealand Chatham Rise | 1825 | 2001-2018 | 16 | 1964 | 40-1291 | 189,500 | GBIF access 2008-present, access 1964-2008, Richard O'Driscoll | PPA | AUR | https://fs.fish.govt.nz/Doc/24639/FAR-2018-41-Trawl-Survey-TAN1801.pdf.ashx |
| 80 | NZ-SA | Oceania | New-Zealand Sub-Antarctic | 1263 | 2001-2018 | 14 | 1964 | 57-994 | 77,900 | GBIF access 2008-present, access 1964-2008, Richard O'Driscoll | PPA | AUR | https://fs.fish.govt.nz/Doc/24637/FAR-2018-39-Trawl-Survey-TAN1614.pdf.ashx |

| | | | | | | | | | | | | | |
|---|---|---|---|---|---|---|---|---|---|---|---|---|---|
| 81 | NZ-WCSI | Oceania | New-Zealand, West coast South Island | 544 | 2003-2017 | 7 | 1<br><br>992 | 13-391 | 28,200 | GBIF access 2008-present, access 1964-2008,<br>Fabrice Stephenson | PPA | AUR | https://fs.fish.govt.nz/Doc/24603/FAR-2018-18-WCSI-Trawl-Survey-2017.pdf.ashx |
| 82 | NZ.ECSI | Oceania | New-Zealand, East coast South Island | 749 | 2007-2018 | 7 | 1991 | 0-392 | 26,500 | GBIF access 2008-present, access 1964-2008,<br>Fabrice Stephenson | PPA | AUR | https://fs.fish.govt.nz/Doc/24664/FAR-2019-03-Inshore-trawl-survey-KAH1803.pdf.ashx |
| 83 | CHI-YS | Asia | China, Yellow Sea | 60 | N/A | N/A | | 13-85 | 143,900 | Xiujuan Shan | IM | | |
| 84 | IND | Asia | India | N/A | N/A | N/A | N/A | N/A | N/A | http://fsi.gov.in/LATEST-WB-SITE/fsi-db-frm.htm<br>http://fsi.gov.in/LATEST-WB-SITE/fsi-ev-act-frm.htm | UM | N/A | N/A |
| 85 | IS-TAU | Asia | Israel, Tel Aviv University | 68 | 2009-2015 | 7 | 2009 | 15-120 | 200 | Jonathan Belmaker | NPA | | https://link.springer.com/article/10.1007/s00227-016-2950-7 |
| 86 | IS-MOAG | Asia | Israel, Israeli Ministry of Agriculture | 894 | 2008-2019 | 10 | 1990 | 22-240 | 2,500 | Oren Sonin & Dori Edelist | NPA | | https://onlinelibrary.wiley.com/doi/full/10.1111/ddi.12002 |
| 87 | IS-NM | Asia | Israel, national monitoring | 34 | 2014-2018 | 5 | | 20-80 | 40 | Nir Stern | NPA | | https://www.sviva.gov.il/infoservices/reservoirinfo/doclib2/publications/p0801-85p0900/p088859b.pdf |

| | | | | | | | | | | | | | |
|---|---|---|---|---|---|---|---|---|---|---|---|---|---|
| 88 | JAP | Asia | Japan | N/A | N/A | N/A | N/A | N/A | N/A | N/A | UM | N/A | N/A |
| 89 | KOR | Asia | Korea | 3223 | 2001-2019 | 18 | 1995 | 2-4595 | 237,500 | Junghwa Choi | NPA | | |
| 90 | MAL | Asia | Malaysia | N/A | N/A | N/A | N/A | N/A | N/A | Abdul Wahab Abdullah | UM | N/A | N/A |
| 91 | NWP | Asia | Russia, North Pacific Ocean | 21988 | 2001-2019 | 18 | 1950 | 2-6203 | 1,705,000 | Vladimir Kulik | NPA | | |
| 92 | WBS | Asia | Russia, Western Bering Sea | 18101 | 2001-2017 | 11 | 1950 | 2-1753 | 222,000 | Vladimir Kulik | NPA | | https://www.nature.com/articles/s41598-018-34819-4 https://www.scirp.org/journal/paperinformation.aspx?paperid=62909 https://archive.org/details/IJOERDEC201519/mode/2up http://www.sciencepublishinggroup.com/journal/paperinfo.aspx?journalid=162&doi=10.11648/j.ijema.20140206.12 https://link.springer.com/article/10.1134/S1063074011070078 |
| 93 | CHL | South America | Chile | 2252 | 2004-2016 | 13 | 2004 | 38-482 | 45,200 | Daniela Yepsen | AUR | AUR | |
| 94 | COL | South America | Colombian Caribbean | 276 | 2001-2016 | 8 | | 8 | 26,400 | Camilo B. Garcia | NPA | | |
| 95 | FLK | South America | Falkland Islands | 1544 | 2001-2019 | 18 | | 2-956 | 172,300 | Alexander Arkhipkin ( | AUR | AUR | https://www.fig.gov.fk/fis |

| | | | | | | | | | | | | | |
|---|---|---|---|---|---|---|---|---|---|---|---|---|---|
| | | | | | | | | | | v.fk) Jorge E. Ramos & Alexander | | | heries/ |
| 96 | PER | South America | Peru | 1714 | 2001-2015 | 15 | 1965 | 20-576 | 51,100 | Jacqueline Palacios León Renato Guevara-Carrasco | NPA | | http://www.imarpe.gob.pe/imarpe/ |
| 97 | ARG | South America | Argentina | N/A | N/A | N/A | N/A | N/A | N/A | | IM | | |
| 98 | POKER | Antarctica | Kerguelen Islands | 853 | 2006-2017 | 4 | 2006 | 70-962 | 170,500 | Félix Massiot-Granier | AUR | AUR | |
| 99 | HIMI | Antarctica | Heard & McDonald Islands | 3644 | 2001-2019 | 19 | 1997 | 150-1100 | 83,100 | Philippe Ziegler https://data.aad.gov.au/metadata/records/HIMI_RSTS_Strata | AUR | AUR | https://www.ccamlr.org/en/wg-fsa-2019/03 (available upon request) |

**Table 1.2:** Summary of metadata collection per availability status.

| Availability status | Number of hauls | Proportion |
|---|---|---|
| *Publicly available* | 116,298 | 41% |
| *Partly publicly available* | 16,758 | 6% |
| *Available upon request* | 72,685 | 25% |
| *Not publicly available* | 76,692 | 27% |
| *Incomplete metadata* | 1,492 | 1% |
| *Unavailable metadata* | N/A | N/A |
| Total | 283,925 | 100% |

### Supplementary 2: Additional surveys not included

In this section, we list the surveys we found but did not match our selection criteria. This is a partial list of many other bottom trawl surveys that might exist. The table lists coastal, sporadic and past surveys we found but did not include. Among them, some were already publicly available, or available upon request. The rest of the surveys are colored in black because they are, to our knowledge, not available for public use. We did not assign the red color as we did not request the metadata from the data providers and only identified that these surveys exist.

**Table 2.1:** Summary of surveys we found and which did not meet the selection criteria. The color code is similar to the rest of the chapter.

| Survey | Continent | Area | Number of years | Provider/Contact | Availability | Reasons for exclusion |
|---|---|---|---|---|---|---|
| DFO-BC-SOG | North America | British Columbia, Strait of Georgia | <5 | NOAA | PA | Too coastal, not enough years |
| NEAMAP | North America | Northeast US inner shelf | | NOAA | AUR | Too coastal |
| SEAMAP | North America | Southeast US inner shelf | | NOAA | PA | Too coastal |
| CHUK | North America | Chukchi Sea | | NOAA | NA | Sporadic survey, last year 2012 |
| FR-CGFS | Europe | Western English Channel | 2 | IFREMER | AUR | Only 2 years old |
| | Europe | Scottish surveys | ? | ? | NA | Old survey |
| AUS-SA | Oceania | Southeast Australia | | CSIRO | PA | Old survey |
| AUS-NA | Oceania | Northwest Australia | | CSIRO | NA | Old survey |
| TAI | Asia | Taiwan | 4 | OBIS | PA | Old survey, and not ongoing |
| VIET | Asia | Vietnam | 2 | Sten Munch-Petersen | NA | Not ongoing, only 2 years |
| FREN-GUI | South America | French Guinea | 2 | Vincent Vallée | NA | Not regular, not ongoing |
| BRAZ | South America | Brasil | | Arnaud Bertrand | NA | Not regular |
| MEX-PC | South America | Mexico, Pacific Coast | | Manuel J. Zetina-Rejón & Romeo Saldívar-Lucio | NA | Not regular |
| BOH-CH | Asia | Bohai Sea, China | | https://www-sciencedirect- | NA | Only 3 years |

| | | | | | | |
|---|---|---|---|---|---|---|
| | | | | com.proxy.findit.dtu.dk/science/article/pii/S0165783619303327<br>Dr Xiujuan Shan<br>older years:<br>https://www.sciencedirect.com/science/article/abs/pii/S0272771403002452?via%3Dihub | | |
| NC | Oceania | New-Caledonia | | | NA | Old & sporadic surveys |
| SP-AFR | Africa | Spanish surveys: Unesco doc. | | https://unesdoc.unesco.org/ark:/48223/pf0000231430 | NA | Old and sporadic surveys |
| FIRST | Asia | Southeast Asian surveys | | https://www.sciencedirect.com/science/article/abs/pii/S0165783606000579?via%3Dihub | PA | Old surveys |
| NAN | World | Nansen | <4 | | NA | |
| | Africa | Benin, Togo, Ivory Coast | | EUMOA surveys, Jérôme Guitton | NA | Sporadic |
| NZ-EEZ | Oceania | New Zealand EEZ | | GBIF access 2008-present, access 1964-2008,<br>Fabrice Stephenson<br> | AUR | Sporadic, not enough years |
| PIGE | Antarctica | Kerguelen Islands | <4 | Félix Massiot-Granier | AUR | Only one year (2015) |
| | Africa | Angola | <4 | Ignacio Sobrino | NA | only one year (2003) |
| | Africa | Mozambique | <4 | Ignacio Sobrino | | only 3 years (2007,2008,2009) |

### Supplementary 3: Defining productive and fished areas

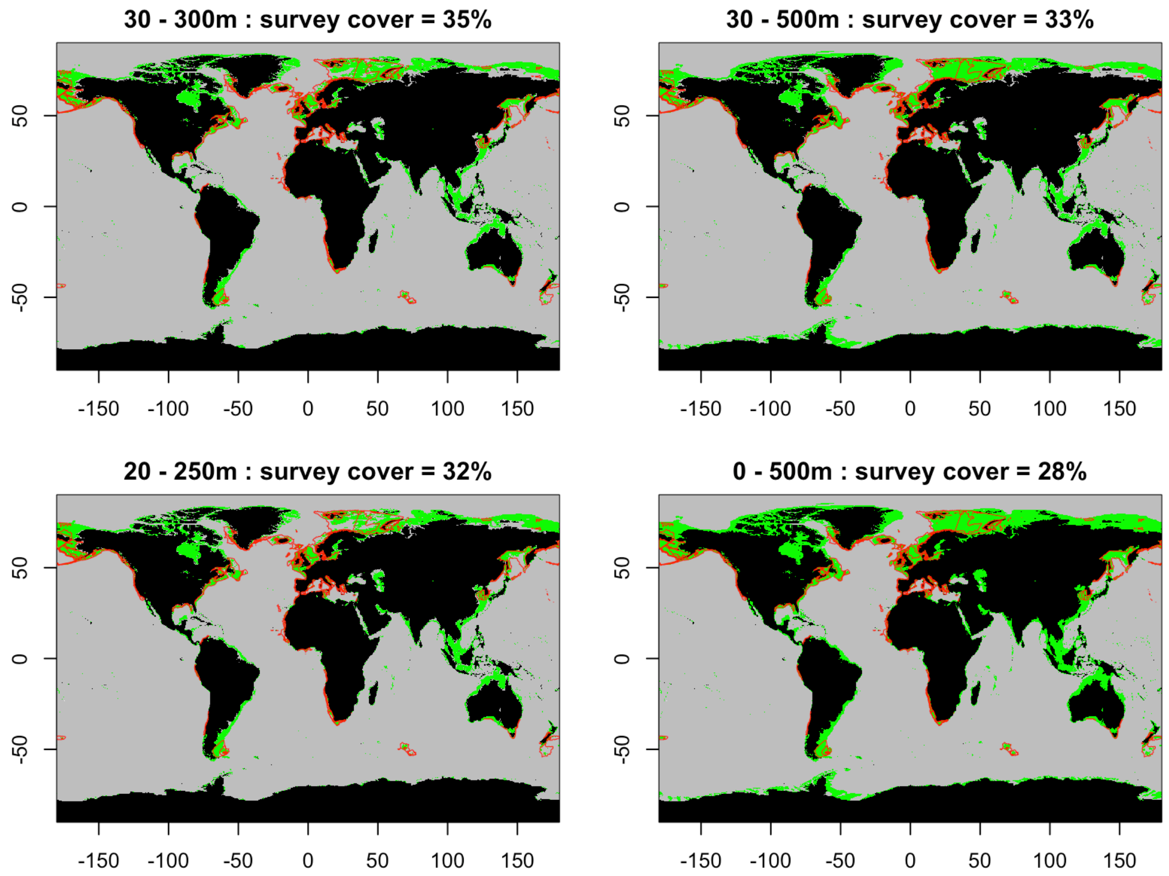

**Figure 3.1:** Map combining continental shelves and slopes (in green) and the outline of the convex hull (red) derived from the surveys. Each sub-panel combines different depth thresholds defining the continental shelf and slope. The proportion indicates the overlap between the convex hull and the green areas. The depth data was extracted from GEBCO (https://www.gebco.net/).

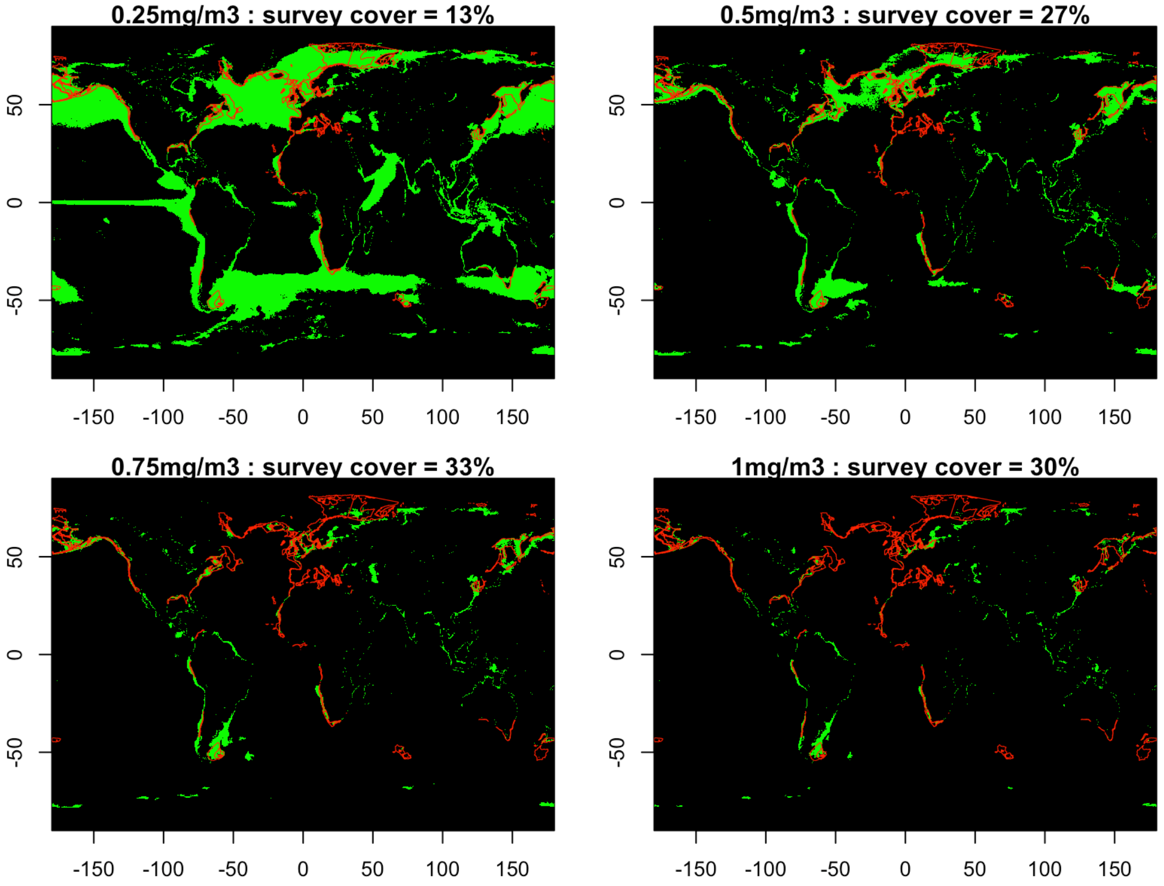

**Figure 3.2:** Map combining the chlorophyll a concentration and the outline of the convex hull derived from the surveys. Each sub-panel combines different chlorophyll a concentration thresholds defining productive sites. The proportion indicates the overlap between the convex hull and the grid cells identified as 'productive'. Red lines show the outline of the survey convex hulls; green shading shows grid cells matching a given threshold for chlorophyll a concentration. The chlorophyll a data were extracted from GlobColour[1].

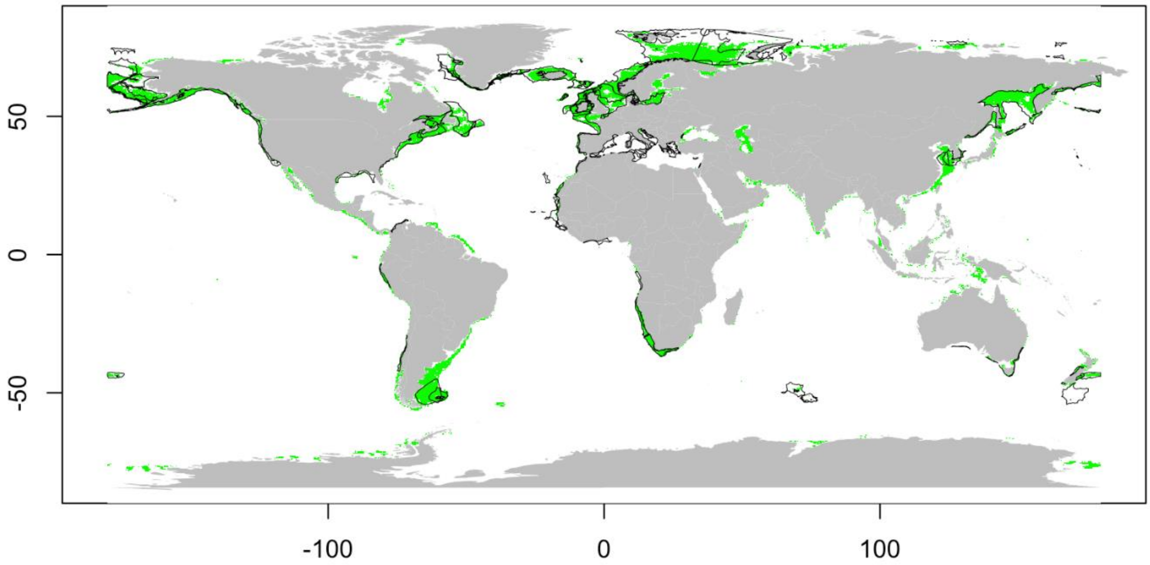

**Figure 3.3:** Map showing areas on the continental shelves and slopes (defined as depth 30-500 m) and high chlorophyll a concentration (using threshold >0.5 mg/m$^3$) in green, and the outline of the convex hull derived from the surveys (black). The overlap indicates a coverage of 62% of productive shelves by the surveys.

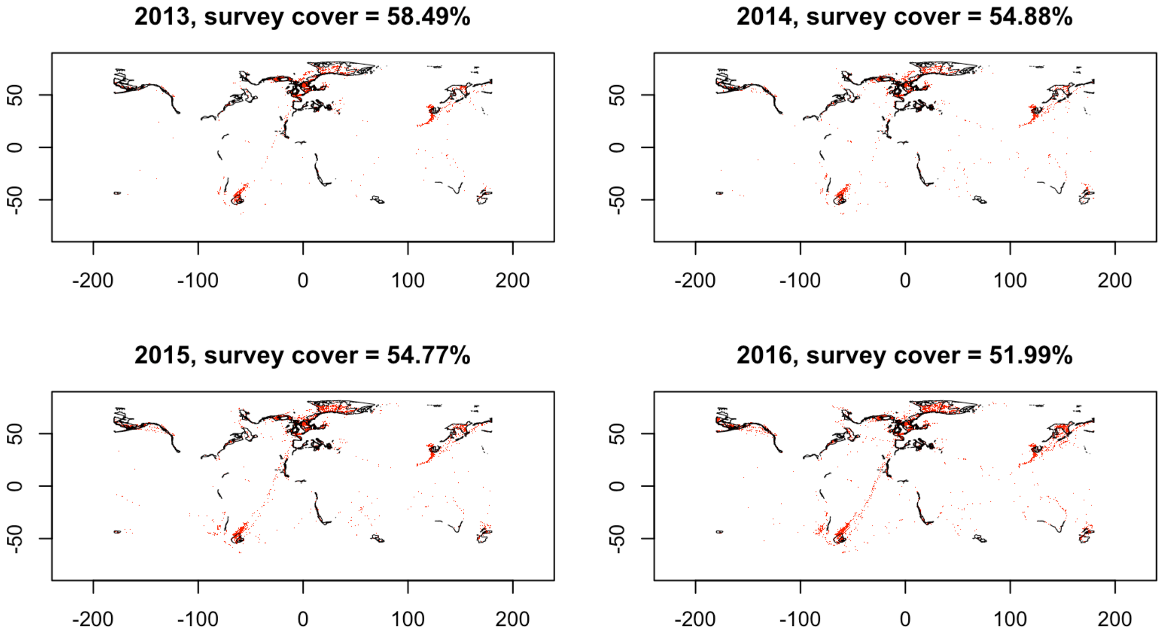

**Figure 3.4:** Map combining the fished sites (red) and the outline of the convex hull derived from the surveys (black). Each sub-panel represents a different year (from 2013 to 2016), from which the Global Fishing Watch data[2] was extracted.

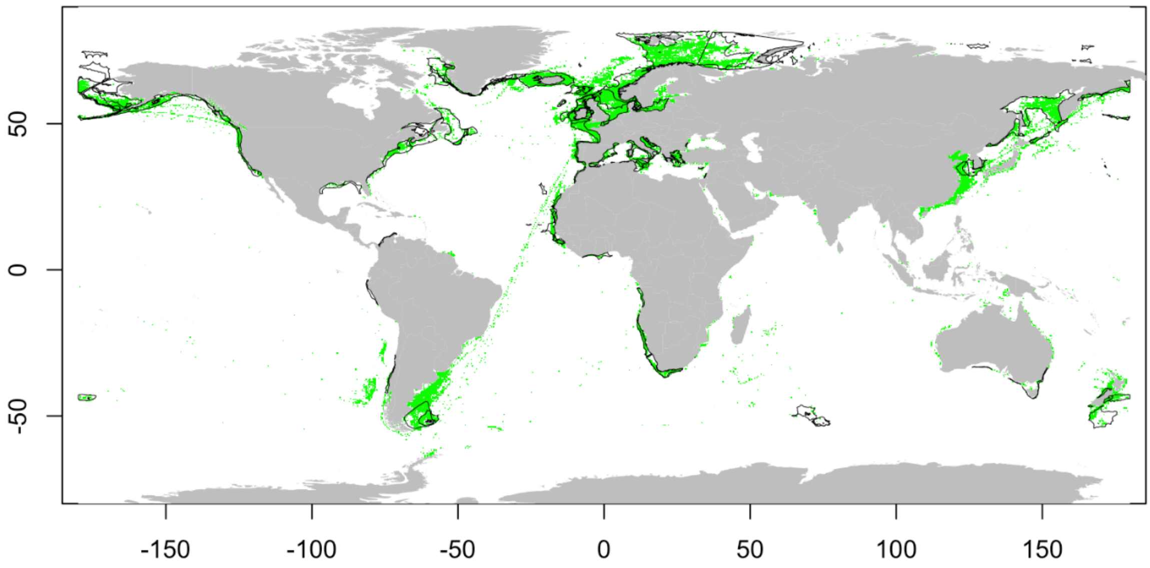

**Figure 3.5:** Map combining the fished areas (green), and the outline of the convex hull from the surveys (black). The overlap indicates a coverage of 54% of trawled fished areas by the surveys (where all years are combined).

### Supplementary 4: Additional maps

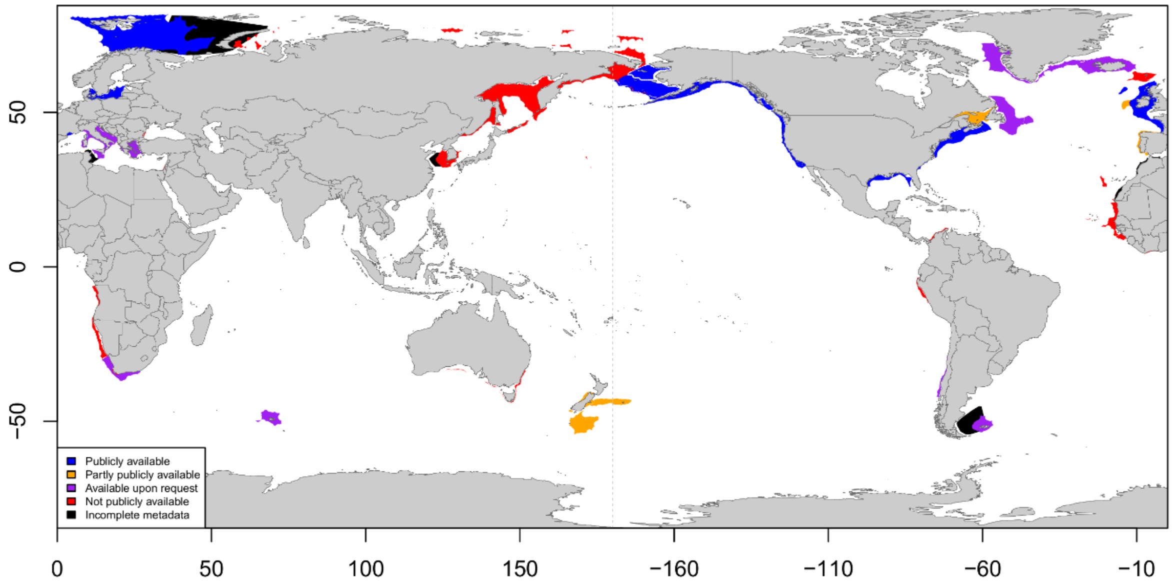

**Figure 4.1:** Worldwide location of ongoing bottom trawl scientific surveys, centered around Asia. Surveys are classified according to their availability: publicly available (blue), partly publicly available (orange), available upon request (purple) and private (red).

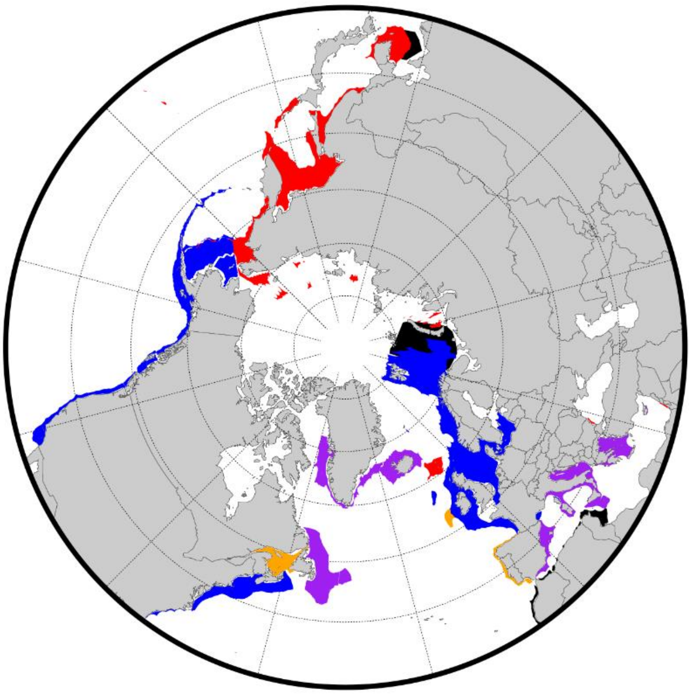

**Figure 4.2:** Location of ongoing bottom trawl scientific surveys in northern seas from a polar view. Surveys are classified according to their availability: publicly available (blue), available upon request (purple), partly publicly available (orange), not publicly available (red) and incomplete metadata (black).

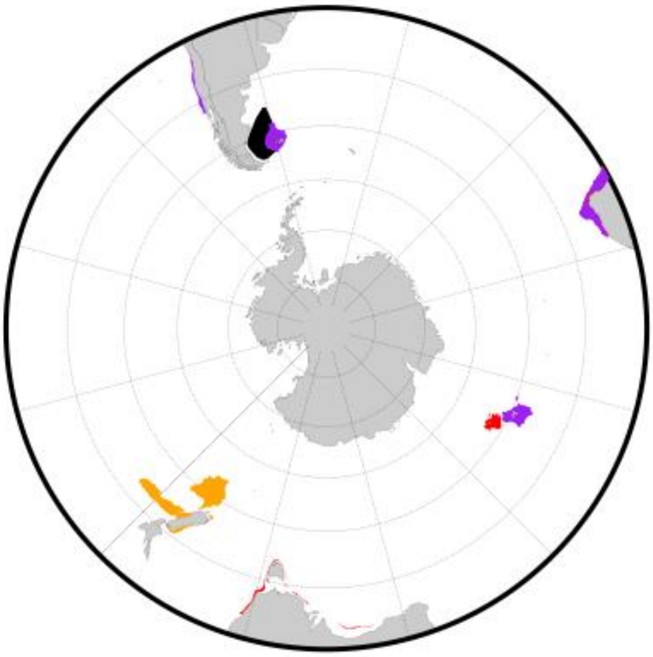

**Figure 4.3:** Location of ongoing bottom trawl scientific surveys in southern seas from a polar view. Surveys are classified according to their availability: publicly available (blue), available upon request (purple), partly publicly available (orange), not publicly available (red) and incomplete metadata (black).

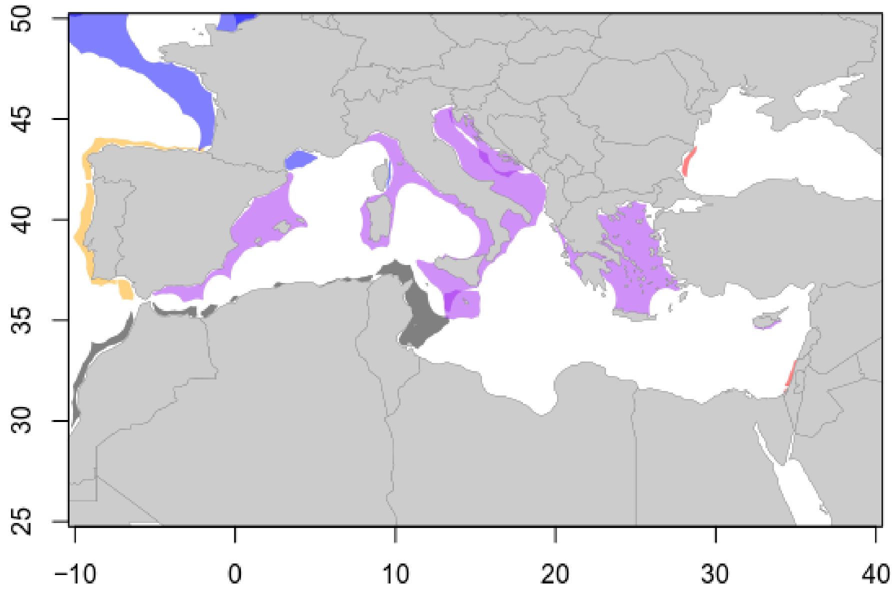

**Figure 4.4:** Location of ongoing bottom trawl scientific surveys in the Mediterranean Sea. Surveys are classified according to their availability: publicly available (blue), partly publicly available (orange), available upon request (purple), not available (red) and incomplete metadata (black).

### Supplementary 5: Species range

File ‘SI.Appendix5.pdf’: Species range maps (from Aquamaps) where the probability of presence is higher than 0.5 (orange) with the survey convex hull overlayed (grey).

**Table 5.1:** List of main FAO fishing areas (http://www.fao.org/fishery/area/search/) included to identify the main demersal commercial species in FishStat, indicated by their scientific name and ASFIS 3-letter code (http://www.fao.org/fishery/collection/asfis/).

| FAO code | Area | Included/ Excluded | Scientific name | ASFIS code | Range in several FAO? |
|---|---|---|---|---|---|
| 18 | Arctic Sea | Included | *Gadus morhua* | COD | yes |
| | | | *Melanogrammus aeglefinus* | HAD | yes |
| | | | *Reinhardtius hippoglossoides* | GHL | yes |
| 21 | Northwest Atlantic | Included | *Gadus morhua* | COD | yes |
| | | | *Melanogrammus aeglefinus* | HAD | yes |
| | | | *Reinhardtius hippoglossoides* | GHL | yes |
| 27 | Northeast Atlantic | Included | *Gadus morhua* | COD | yes |
| | | | *Pollachius virens* | POK | yes |
| | | | *Melanogrammus aeglefinus* | HAD | yes |
| 31 | Western Central Atlantic | Included | *Mugil cephalus* | MUF | yes |
| | | | *Lutjanus campechanus* | SNR | yes |
| | | | *Ocyurus chrysurus* | SNY | yes |
| 34 | Eastern Central Atlantic | Included | *Polydactylus quadrifilis* | TGA | no |
| | | | *Brachydeuterus auritus* | GRB | no |
| | | | *Trichiurus lepturus* | LHT | yes |
| 37 | Mediterranean and Black Sea | Included | *Merluccius merluccius* | HKE | yes |
| | | | *Boops boops* | BOG | yes |
| | | | *Merlangius merlangus* | WHG | yes |
| 41 | Southwest Atlantic | Included | *Merluccius hubbsi* | HKP | no |
| | | | *Micropogonias furnieri* | CKM | yes |
| | | | *Macruronus magellanicus* | GRM | yes |
| 47 | Southeast Atlantic | Included | *Merluccius capensis* | HKK | no |
| | | | *Thyrsites atun* | SNK | yes |
| | | | *Lophius vomerinus* | MVO | no |
| 48 | Antarctic Atlantic | Excluded | N/A | N/A | N/A |
| 51 | Western Indian Ocean | Included | *Harpadon nehereus* | BUC | yes |
| | | | *Trichiurus lepturus* | LHT | yes |
| | | | *Otolithes ruber* | LKR | yes |
| 57 | Eastern Indian Ocean | Included | *Harpadon nehereus* | BUC | yes |
| | | | *Cephalopholis boenak* | CVK | yes |
| | | | *Trichiurus lepturus* | LHT | yes |
| 58 | Antarctic Indian Ocean | Included | *Dissostichus eleginoides* | TOP | yes |
| | | | *Macrourus carinatus* | MCC | yes |
| | | | *Champsocephalus gunnari* | ANI | yes |
| 61 | Northwest Pacific | Included | *Trichiurus lepturus* | LHT | yes |
| | | | *Gadus chalcogrammus* | ALK | yes |
| | | | *Larimichthys polyactis* | CRY | no |
| 67 | Northeast Pacific | Included | *Gadus chalcogrammus* | ALK | yes |
| | | | *Merluccius productus* | NHA | yes |
| | | | *Gadus macrocephalus* | PCO | yes |
| 71 | Western Central Pacific | Included | *Cephalopholis boenak* | CVK | yes |
| | | | *Plectropomus leopardus* | EMO | yes |
| | | | *Mene maculata* | MOO | yes |
| 77 | Eastern Central Pacific | Included | *Lutjanus peru* | LWP | no |
| | | | *Merluccius productus* | NHA | yes |
| | | | *Mugil cephalus* | MUF | yes |
| 81 | Southwest Pacific | Included | *Macruronus novaezelandiae* | GRN | yes |
| | | | *Micromesistius australis* | POS | yes |
| | | | *Thyrsites atun* | SNK | yes |
| 87 | Southeast Pacific | Included | *Merluccius gayi gayi* | PHA | no |
| | | | *Normanichthys crockeri* | NRC | no |
| | | | *Macruronus magellanicus* | GRM | yes |
| 88 | Antarctic Pacific | Excluded | N/A | N/A | N/A |

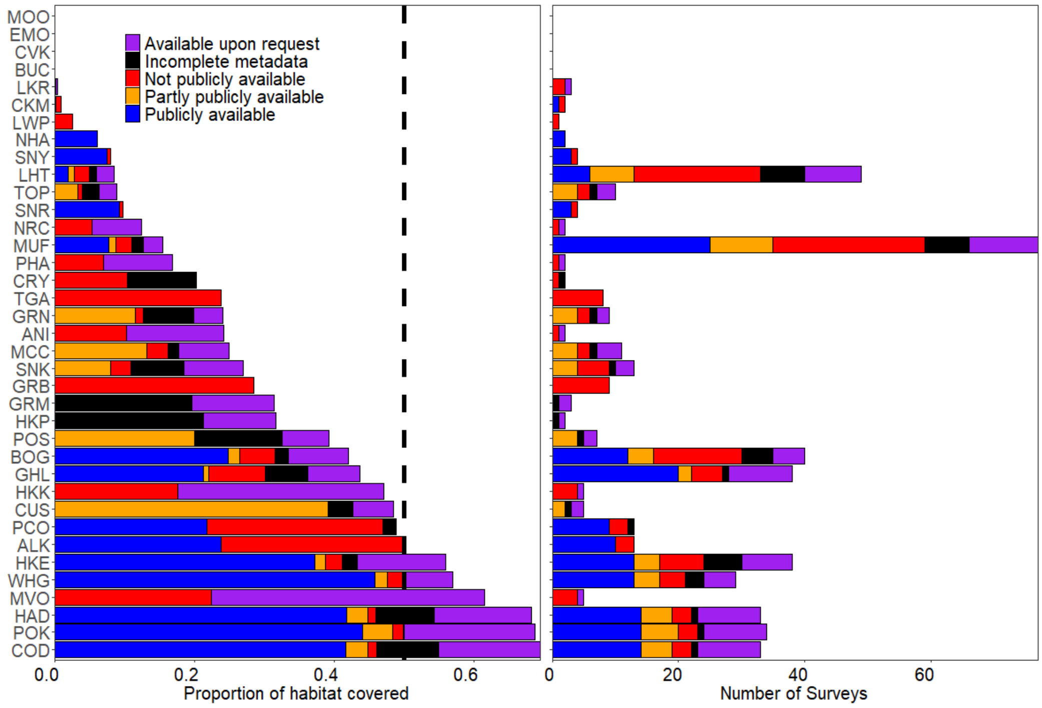

**Figure 5.1:** Main commercial demersal species and the corresponding coverage by the surveys: proportion of AquaMaps habitat covered by the surveys (left panel) and number of surveys behind the proportion covered (right panel). '*' next to species Latin names indicate if the species appeared as a main commercial species in multiple FAO areas. Colors indicate the availability status attributed to each survey. We use a threshold of 0.1 to define presence of species.

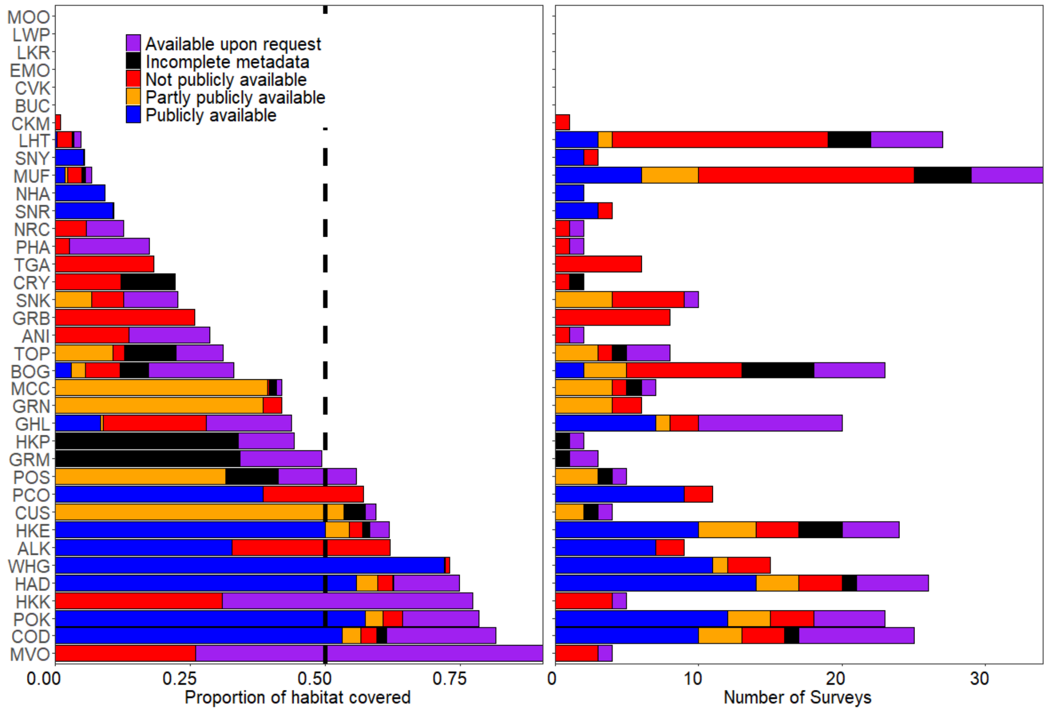

**Figure 5.2:** Main commercial demersal species and the corresponding coverage by the surveys: proportion of AquaMaps habitat covered by the surveys (left panel) and number of surveys behind the proportion covered (right panel). '*' next to species Latin names indicate if the species appeared as a main commercial species in multiple FAO areas. Colors indicate the availability status attributed to each survey. We use a threshold of 0.9 to define presence of species.

**Table 5.2:** Comparison of results when using various thresholds to determine the probability of presence from Aquamaps models.

| **Threshold** | **0.5** | **0.1** | **0.9** |
|---|---|---|---|
| **Max coverage (%)** | 79 | 70 | 90 |
| **Min. number of surveys** | 4 | 5 | 4 |
| **Max. number of surveys** | 31 | 38 | 26 |
| **Mean number of surveys** | 18 | 29 | 14 |

### Supplementary 6: Spatio-temporal modeling

The model incorporates 10 surveys including the eastern Bering Sea, northern Bering Sea, Bering Sea slope, Gulf of Alaska, Aleutian Islands, British Columbia (West Coast of Vancouver Island, Queen Charlotte, Hecate Strait and West Coast Haida Gwaii) and California Current. Between 2001 and 2018, the analysis involves 33,667 samples.

We fitted a Poisson-link delta spatio-temporal model[3] with the vector-autoregressive spatio-temporal model package (VAST)[4,5], with a gamma distribution for positive biomass catch rates. For each of the two components of the delta-model (presence-absence and biomasses), we estimated an annual intercept, a Gaussian Markov random field (GMRF) representing spatial variation in expected densities across all years (“spatial variation”), as well as a separate GMRF for each year representing annual variation in spatial densities (“spatio-temporal variation”).

We estimate density at 500 “knots” and use bilinear interpolation to predict spatial and spatio-temporal variables at locations between those knots. We also specify that spatial correlations decline as a function of distance and directional orientation (“geometric anisotropy”[6]). We then use a generalization of the delta-method to calculate standard errors[7], and the epsilon bias-correction estimator to account for retransformation bias[8] when calculating total abundance and center-of-gravity within each survey region. VAST uses Template Model Builder[9] to implement the Laplace approximation to the marginal likelihood, and we used the gradient of this approximation to identify the maximum likelihood estimate of all parameters within the R statistical environment[10].

In the model, the following parameters were estimated: the variance of spatial and spatio-temporal variation for each delta-model component (four variance parameters), the decorrelation rate for each component (two scale-range parameters), the variance of residual (sampling) variation (one parameter), the intercept for each delta-model component in each year (36 parameters), geometric anisotropy (2 parameters), and the degree of autocorrelation in spatio-temporal variation for each delta-model component (2 parameters).

We improve computational efficiency for calculating the probability of spatial and spatio-temporal variables by using the stochastic partial differential equation (SPDE)[11], and use the mesh generated by R-INLA[12].

To further improve computational speed when implementing the epsilon-estimator given a predictive-process framework, we reduce the number of locations used to measure distribution by identifying 2000 “extrapolation-cells” that approximate the total area sampled by these seven surveys.

We confirm that the model has converged by checking that the absolute gradient of the marginal likelihood with respect to fixed effects is low (<0.0001) and the Hessian is positive definite. We also checked standard model residuals (spatial Pearson residuals, quantile-quantile plots, and the ratio of predicted and observed encounter probabilities), to show that the model has satisfactory fit (i.e., that residuals show little evidence of residual spatial processes).

To check the accuracy of the model, we specifically compared estimated abundance with stock-assessment estimates for the Aleutian Islands and Bering Sea[13] (Spies et al. 2018, Fig. 6.26), the Gulf of Alaska[14] (Spies and Palsson 2018, Table 7.1), and the California Current[15] (Sampson et al. 2017, Fig. 26) and confirm that results have similar scale and trends to those single-region estimates. By contrast, the assessment in British Columbia[16] estimates separate abundance indices for each sub-area within the British Columbia bottom trawl domain, and also includes indices only for 2003-2014. It is therefore difficult to compare indices from the British Columbia assessment document directly with our results for that region.

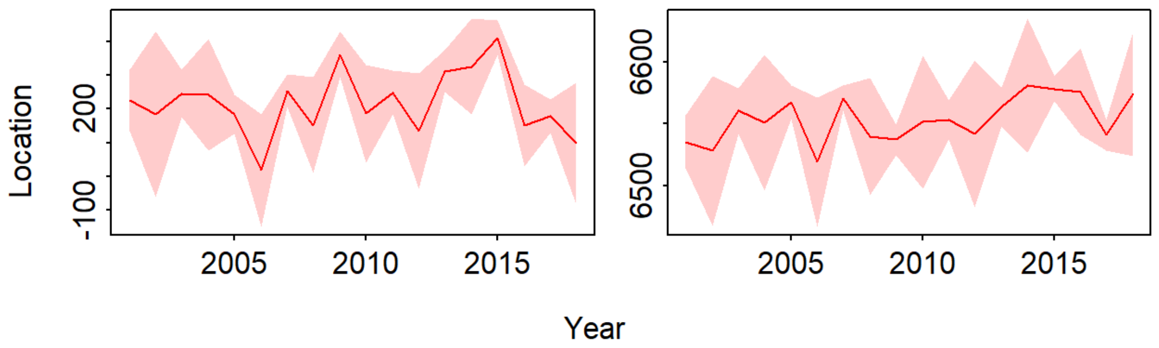

**Figure 6.1:** Center of gravity (y-axis) in each year 2001-2018 (x-axis) showing location (red lines) in kilometers east of UTM zone 3 (left panel) and kilometers north of the equator (right panel), as well as a predictive interval (red shading: +/- one standard error).

### Supplementary References

1. Maritorena, S., d'Andon, O. H. F., Mangin, A. & Siegel, D. A. Merged satellite ocean color data products using a bio-optical model: Characteristics, benefits and issues. *Remote Sens. Environ.* **114**, 1791–1804 (2010).
2. Kroodsma, D. A. *et al.* Tracking the global footprint of fisheries. *Science* **359**, 904–908 (2018).
3. Thorson, J. T. Three problems with the conventional delta-model for biomass sampling data, and a computationally efficient alternative. *Can. J. Fish. Aquat. Sci.* **75**, 1369–1382 (2017).
4. Thorson, J. T. Guidance for decisions using the Vector Autoregressive Spatio-Temporal (VAST) package in stock, ecosystem, habitat and climate assessments. *Fish. Res.* **210**, 143–161 (2019).
5. Thorson, J. T. & Barnett, L. A. K. Comparing estimates of abundance trends and distribution shifts using single- and multispecies models of fishes and biogenic habitat. *ICES J. Mar. Sci.* **74**, 1311–1321 (2017).
6. Thorson, J. T., Shelton, A. O., Ward, E. J. & Skaug, H. J. Geostatistical delta-generalized linear mixed models improve precision for estimated abundance indices for West Coast groundfishes. *ICES J. Mar. Sci.* **72**, 1297–1310 (2015).
7. Kass, R. E. & Steffey, D. Approximate Bayesian Inference in Conditionally Independent Hierarchical Models (Parametric Empirical Bayes Models). *J. Am. Stat. Assoc.* **84**, 717–726 (1989).
8. Thorson, J. T. & Kristensen, K. Implementing a generic method for bias correction in statistical models using random effects, with spatial and population dynamics examples. *Fish. Res.* **175**, 66–74 (2016).
9. Kristensen, K., Nielsen, A., Berg, C. W., Skaug, H. & Bell, B. M. TMB: Automatic Differentiation and Laplace Approximation. *J. Stat. Softw.* **70**, 1–21 (2016).
10. R Core Team. *R: A Language and Environment for Statistical Computing*. (R Foundation for Statistical Computing, 2018).
11. Lindgren, F., Rue, H. & Lindström, J. An explicit link between Gaussian fields and Gaussian Markov random fields: the stochastic partial differential equation approach. *J. R. Stat. Soc. Ser. B Stat. Methodol.* **73**, 423–498 (2011).
12. Rue, H., Martino, S. & Chopin, N. Approximate Bayesian inference for latent Gaussian models by using integrated nested Laplace approximations. *J. R. Stat. Soc. Ser. B Stat. Methodol.* **71**, 319–392 (2009).
13. Spies, I., Wilderbuer, T. K., Nichol, D. G., Hoff, J. & Palsson, W. *Assessment of the arrowtooth flounder stock in the Eastern Bering Sea and Aleutian Islands*. (2018).
14. Spies, I. & Palsson, W. *Assessment of the arrowtooth flounder stock in the Gulf of Alaska*. (2018).
15. Sampson, D. B. *et al. 2017 Assessment Update for the US West Coast Stock of Arrowtooth Flounder*. (2017).
16. Grandin, C. J. & Forrest, R. E. *Arrowtooth Flounder (Atheresthes stomias) Stock Assessment for the West Coast of British Columbia*. v + 87p. (2017).
