## Supplementary 5 for "Are we ready to track climate-driven shifts in marine species across international boundaries? - A global survey of scientific bottom trawl data"

Boops boops (BOG)

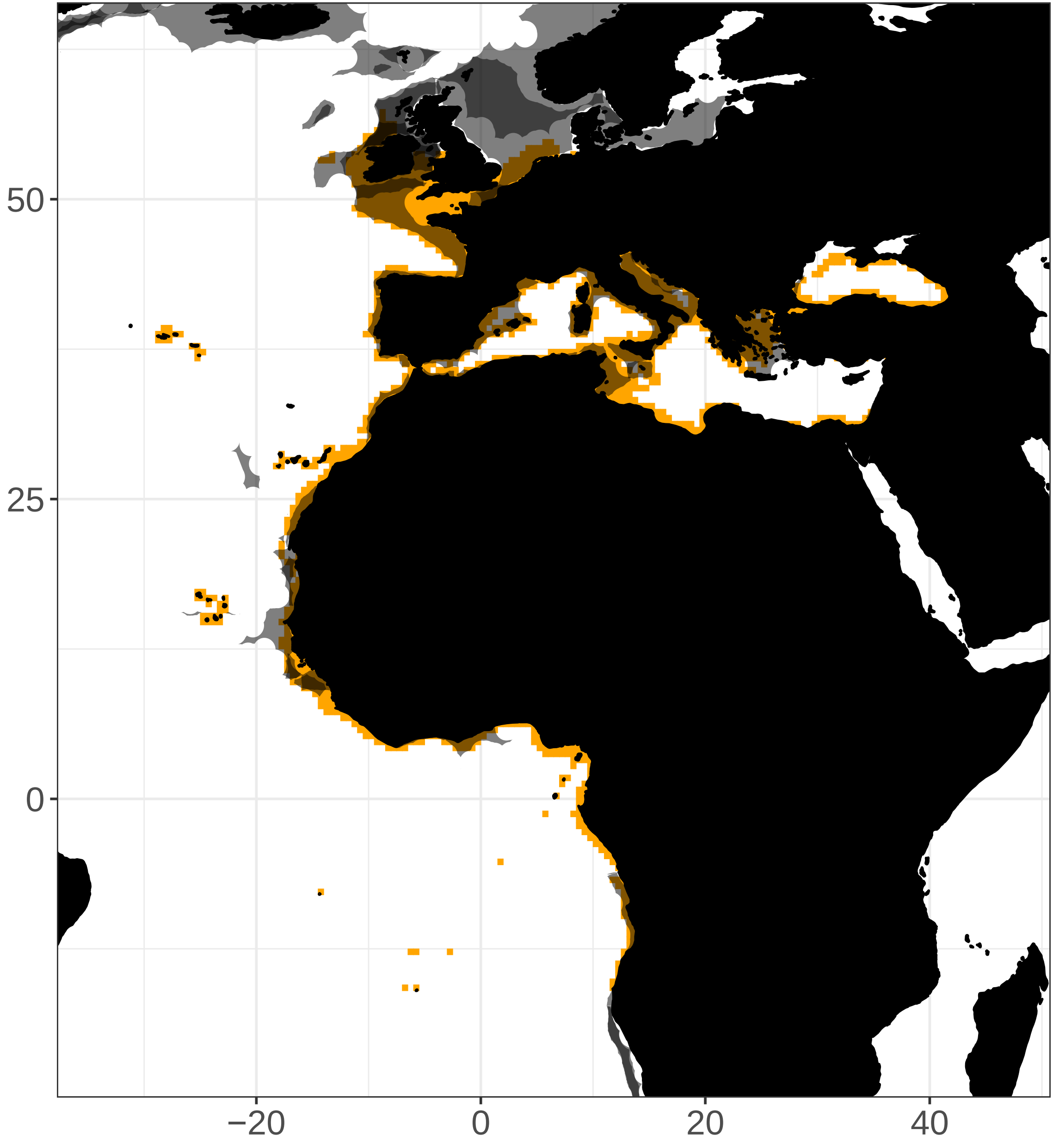

Brachydeuterus auritus (GRB)

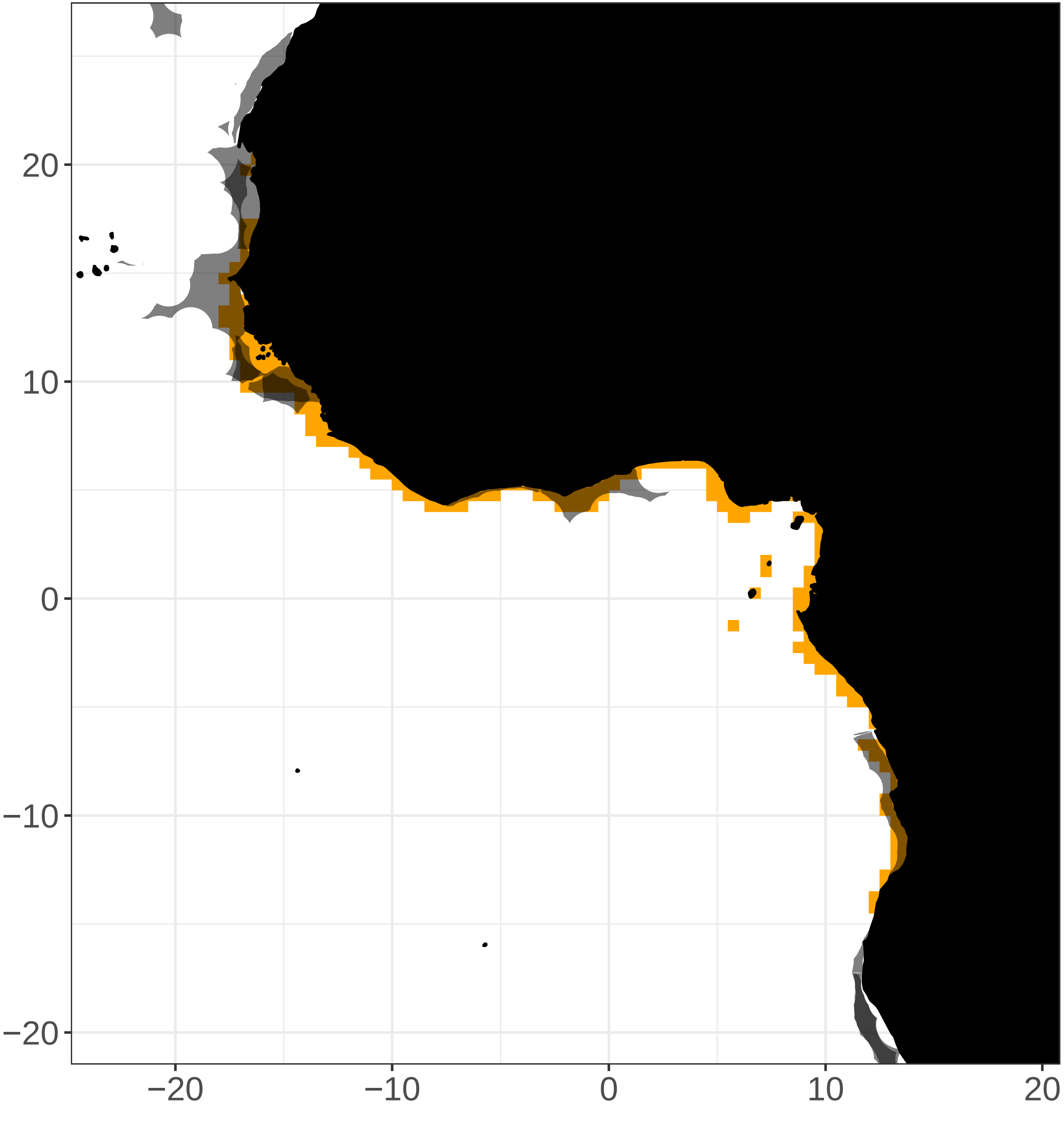

Cephalopholis boenak (CVK)

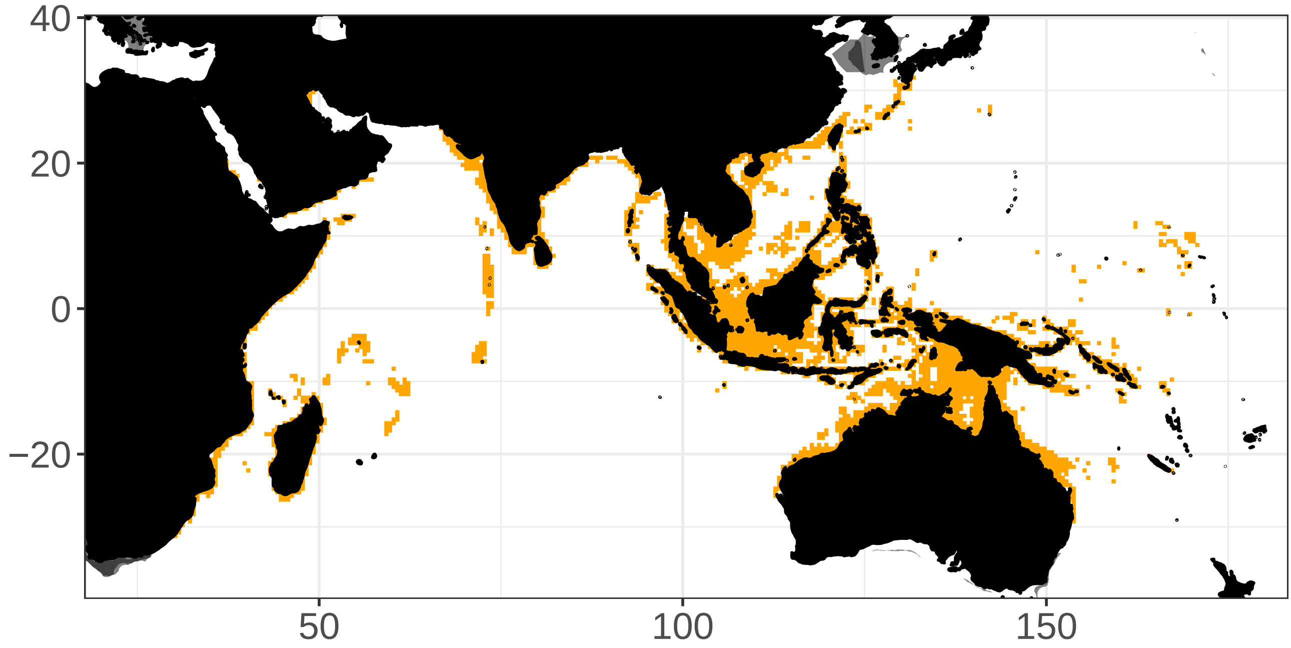

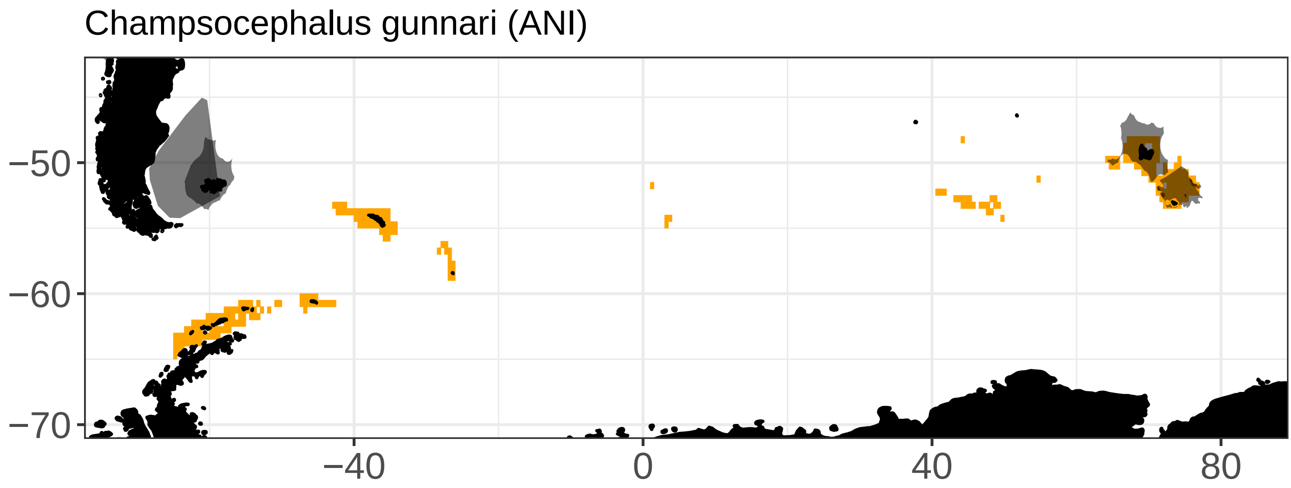
Champsocephalus gunnari (ANI)
−50
−60
−70
−40
0
40
80

Dissostichus eleginoides (TOP)

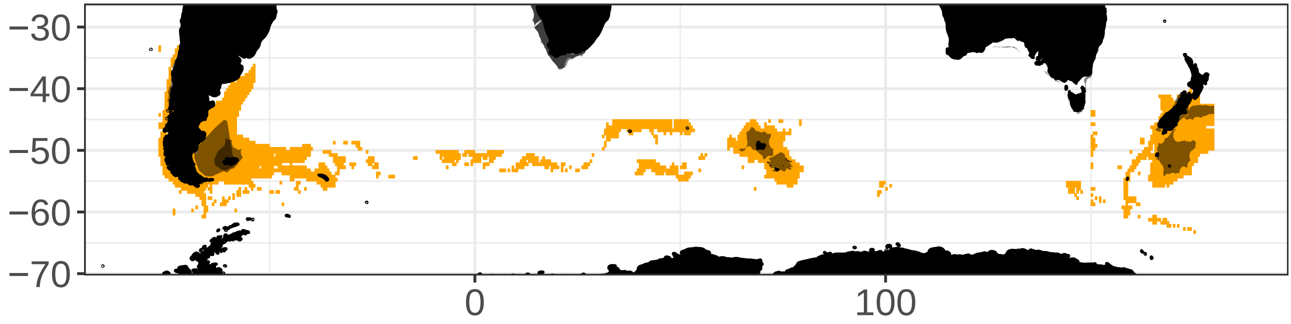

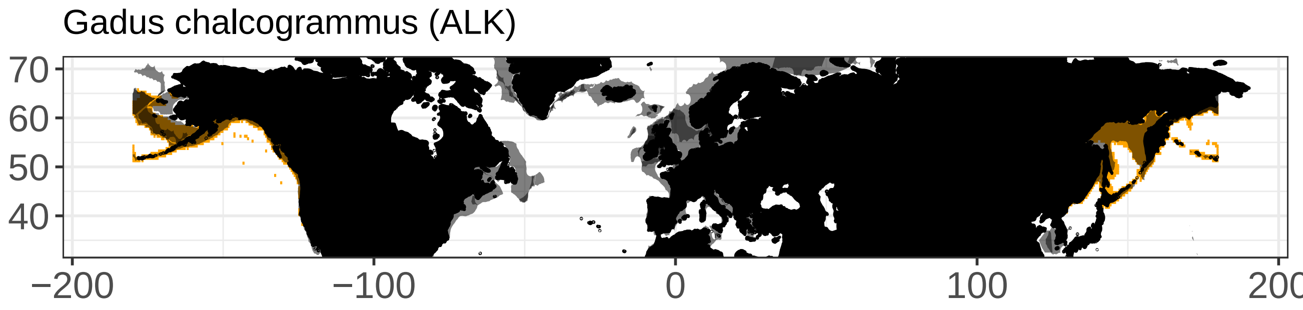

Gadus chalcogrammus (ALK)
70
60
50
40
−200
−100
0
100
200

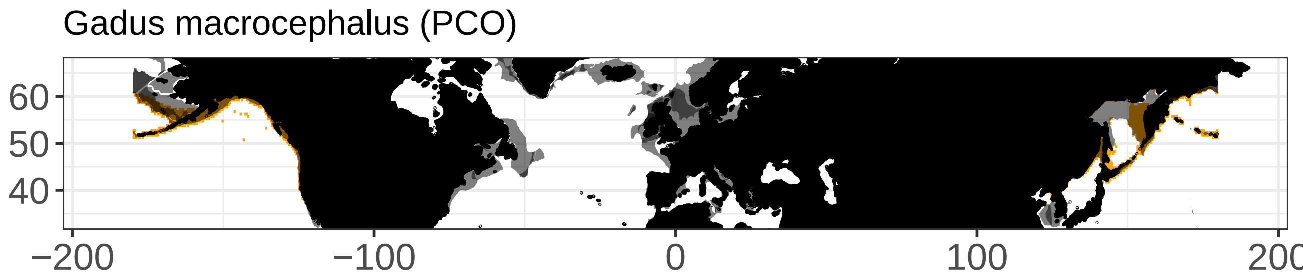

Gadus macrocephalus (PCO)
60
50
40
−200
−100
0
100
200

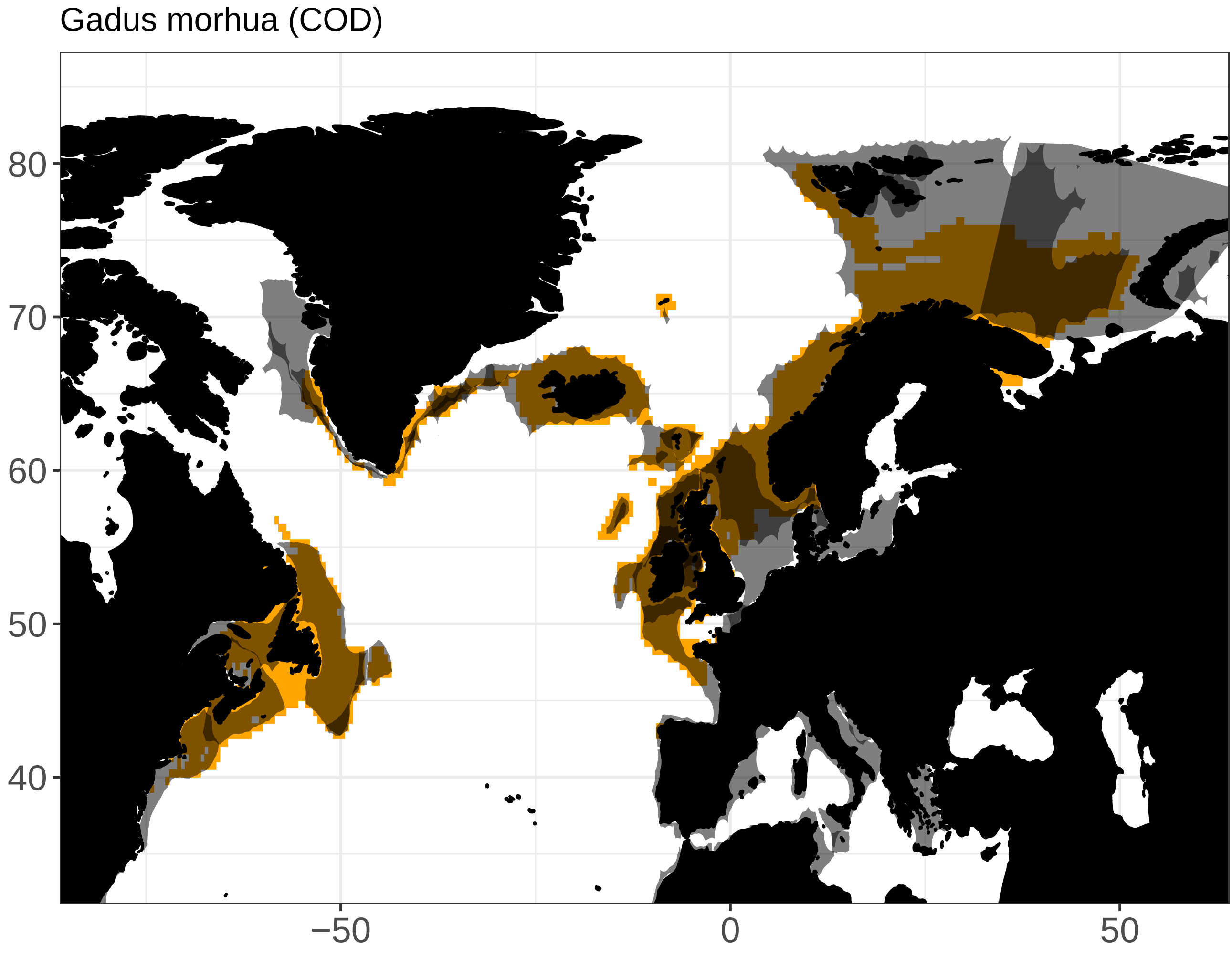

Gadus morhua (COD)
80
70
60
50
40
−50
0
50

Genypterus blacodes (CUS)

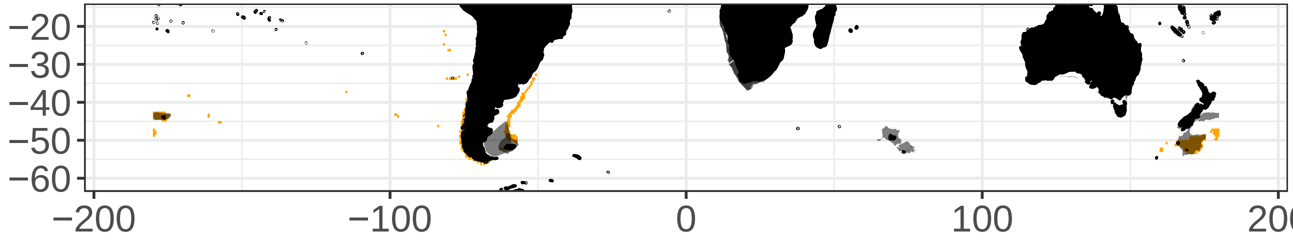

Harpadon nehereus (BUC)

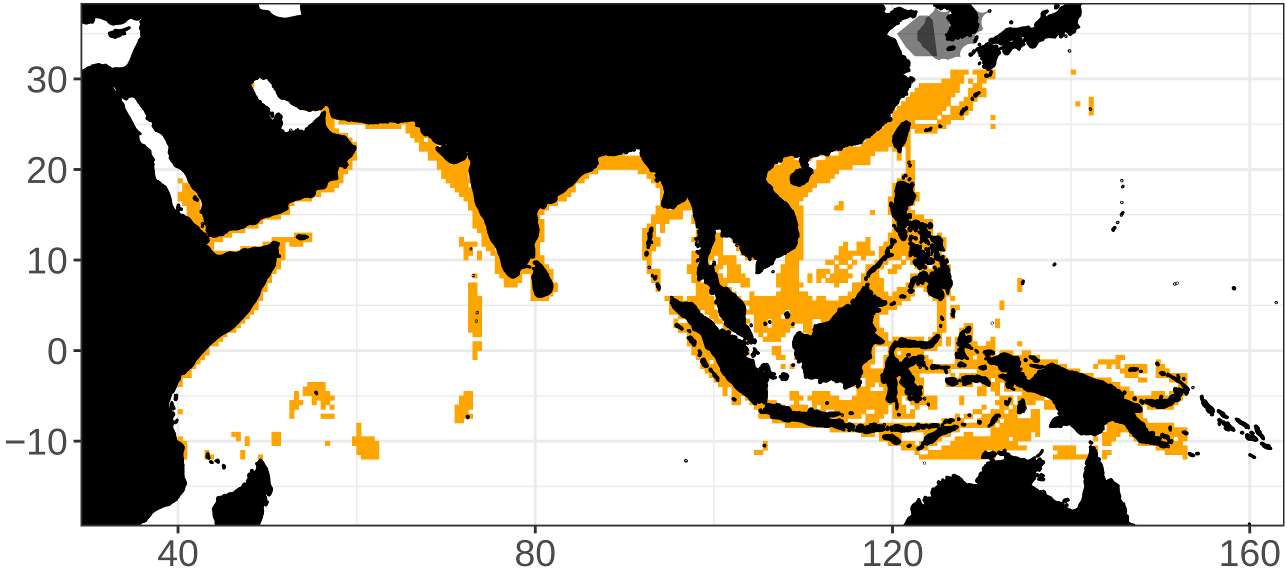

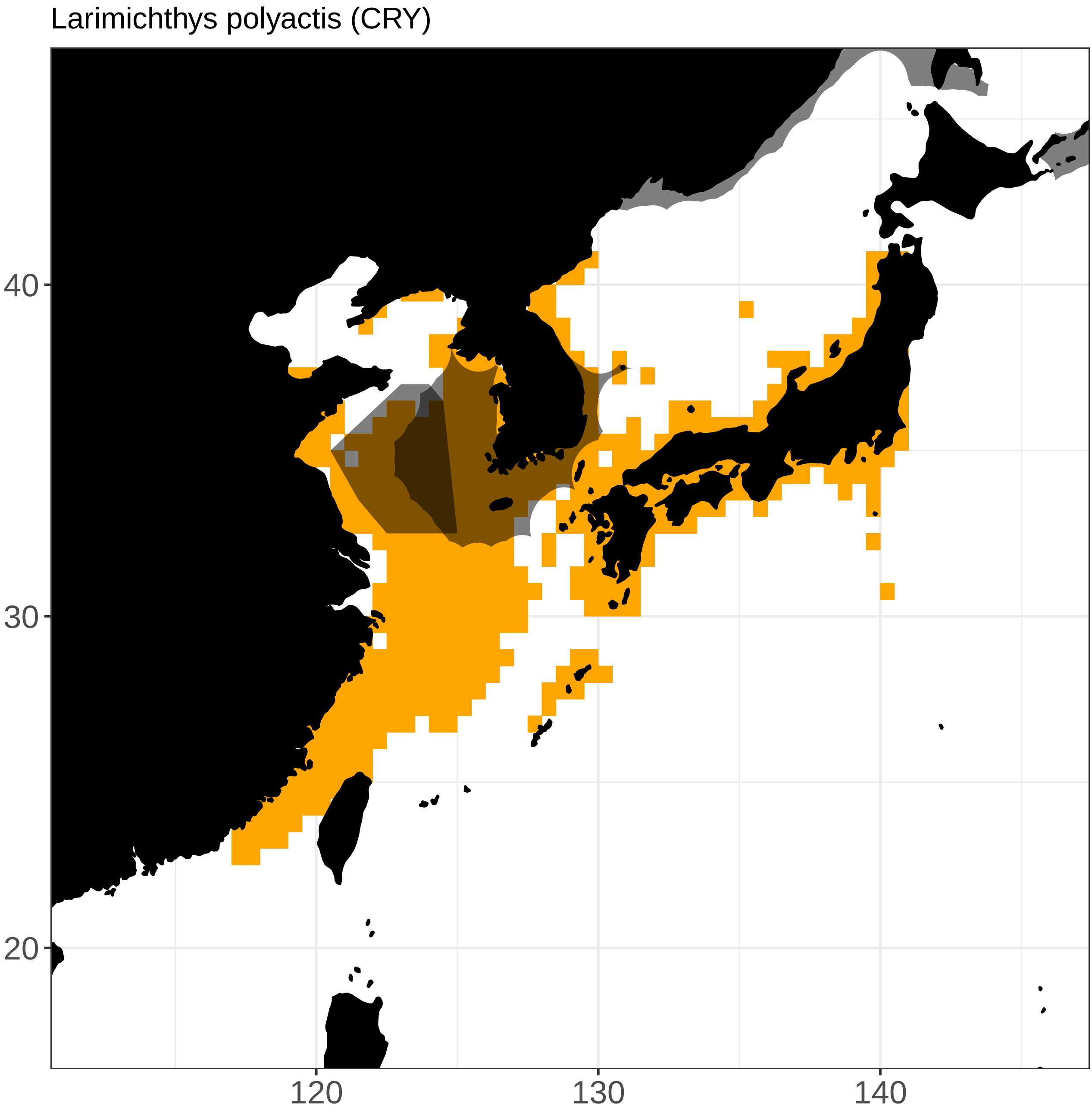

Larimichthys polyactis (CRY)
40
30
20
120
130
140

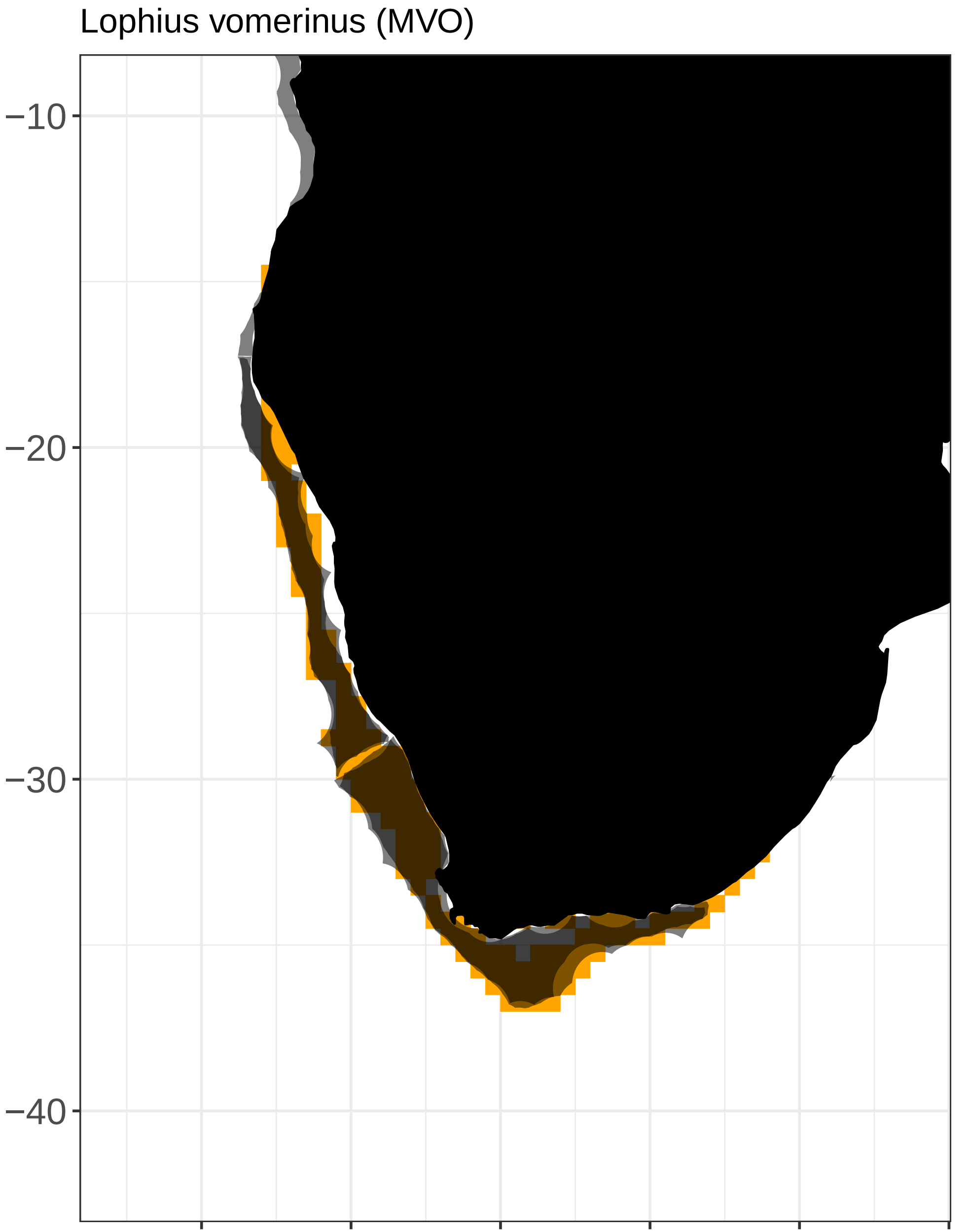

Lophius vomerinus (MVO)
−10
−20
−30
−40
10
15
20
25
30
35

Lutjanus campechanus (SNR)

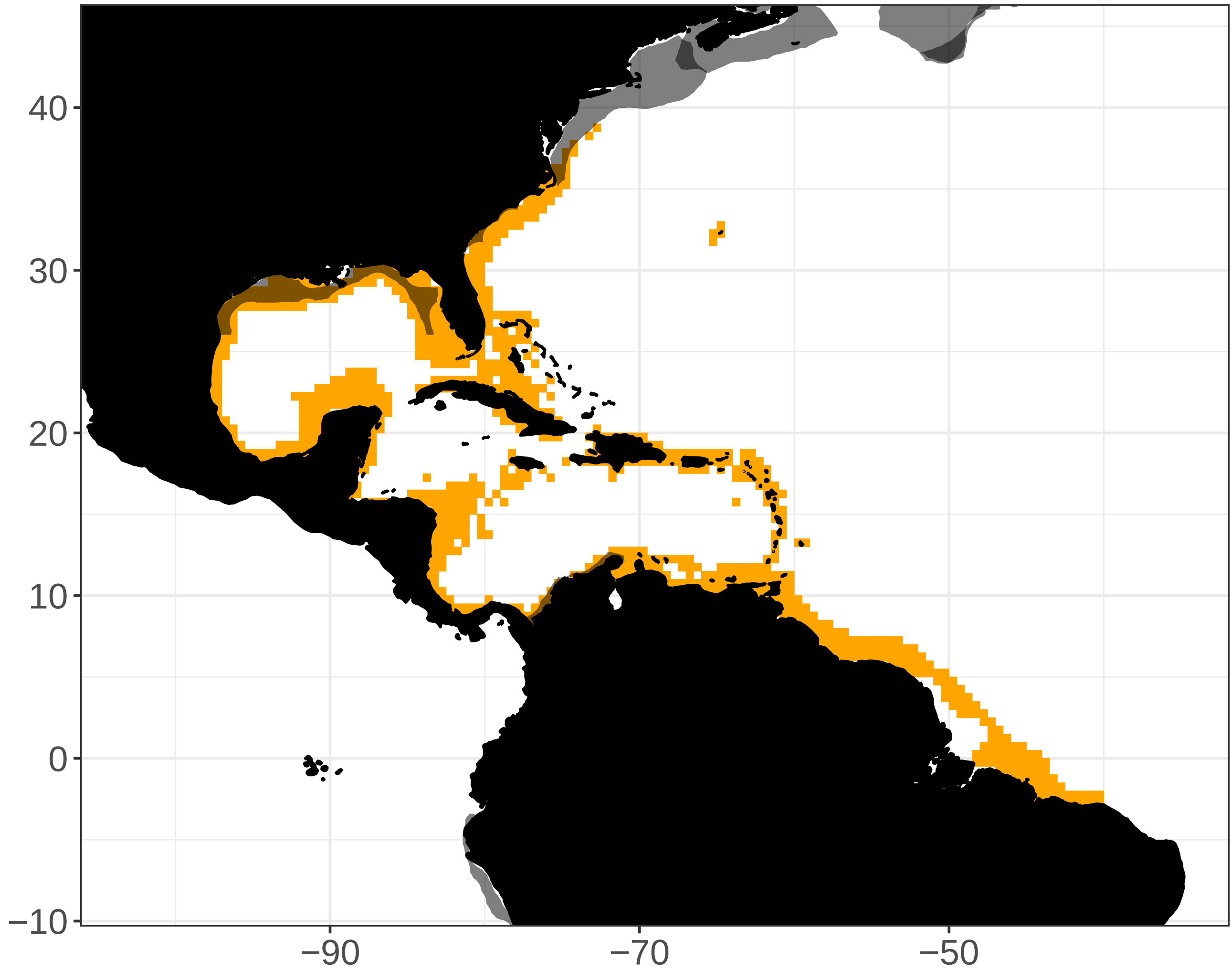

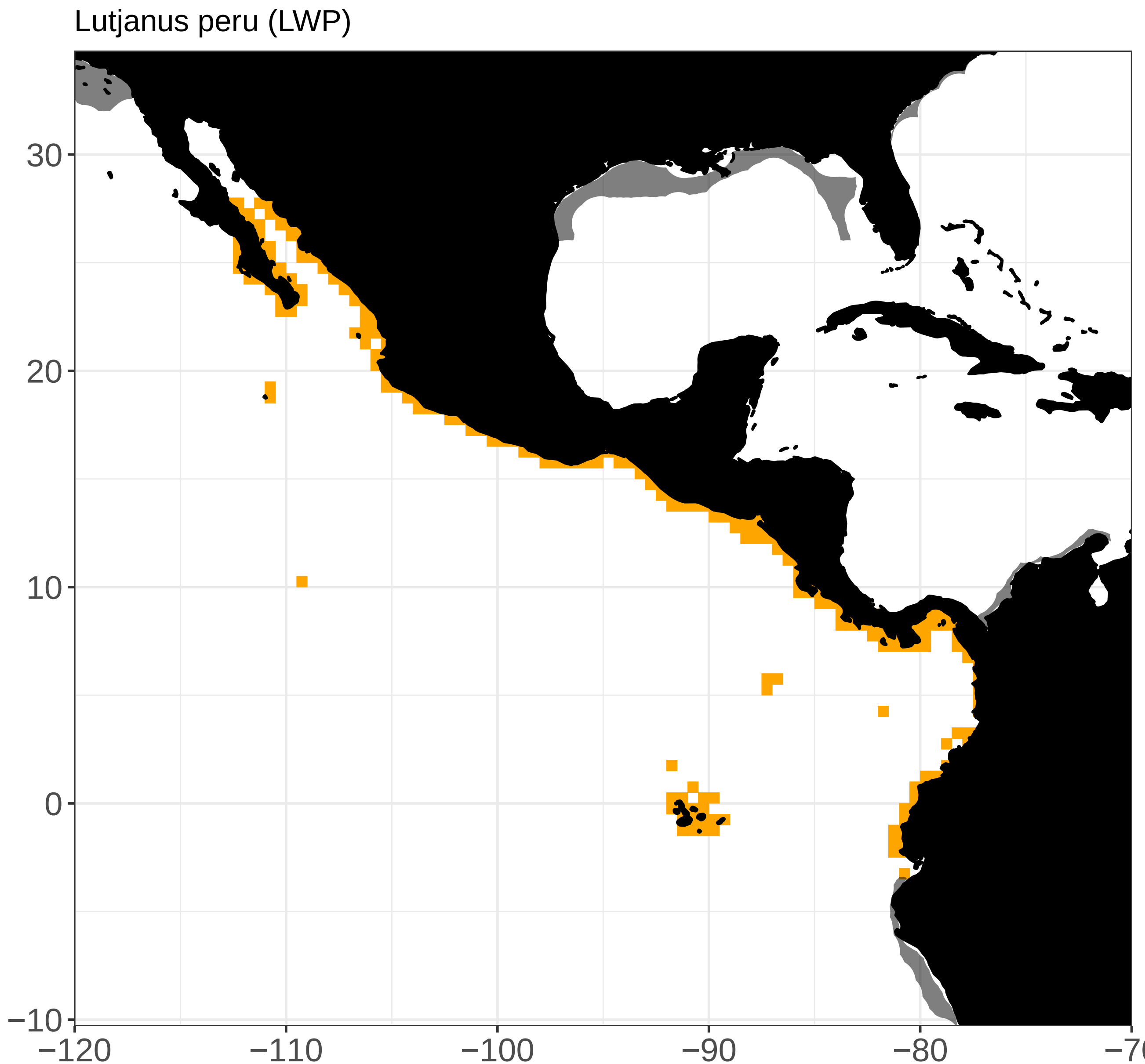

Lutjanus peru (LWP)
30
20
10
0
−10
−120
−110
−100
−90
−80
−70

Macrourus carinatus (MCC)

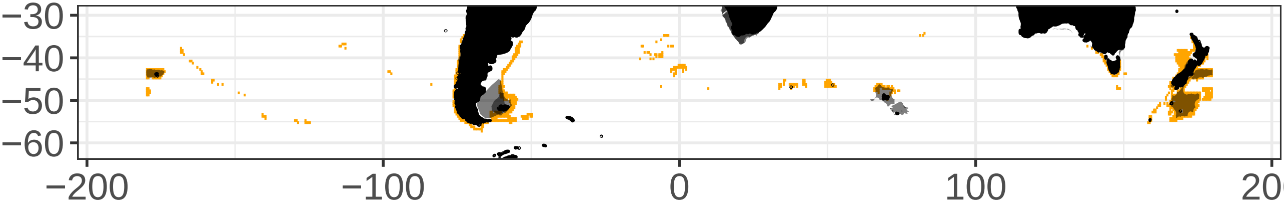

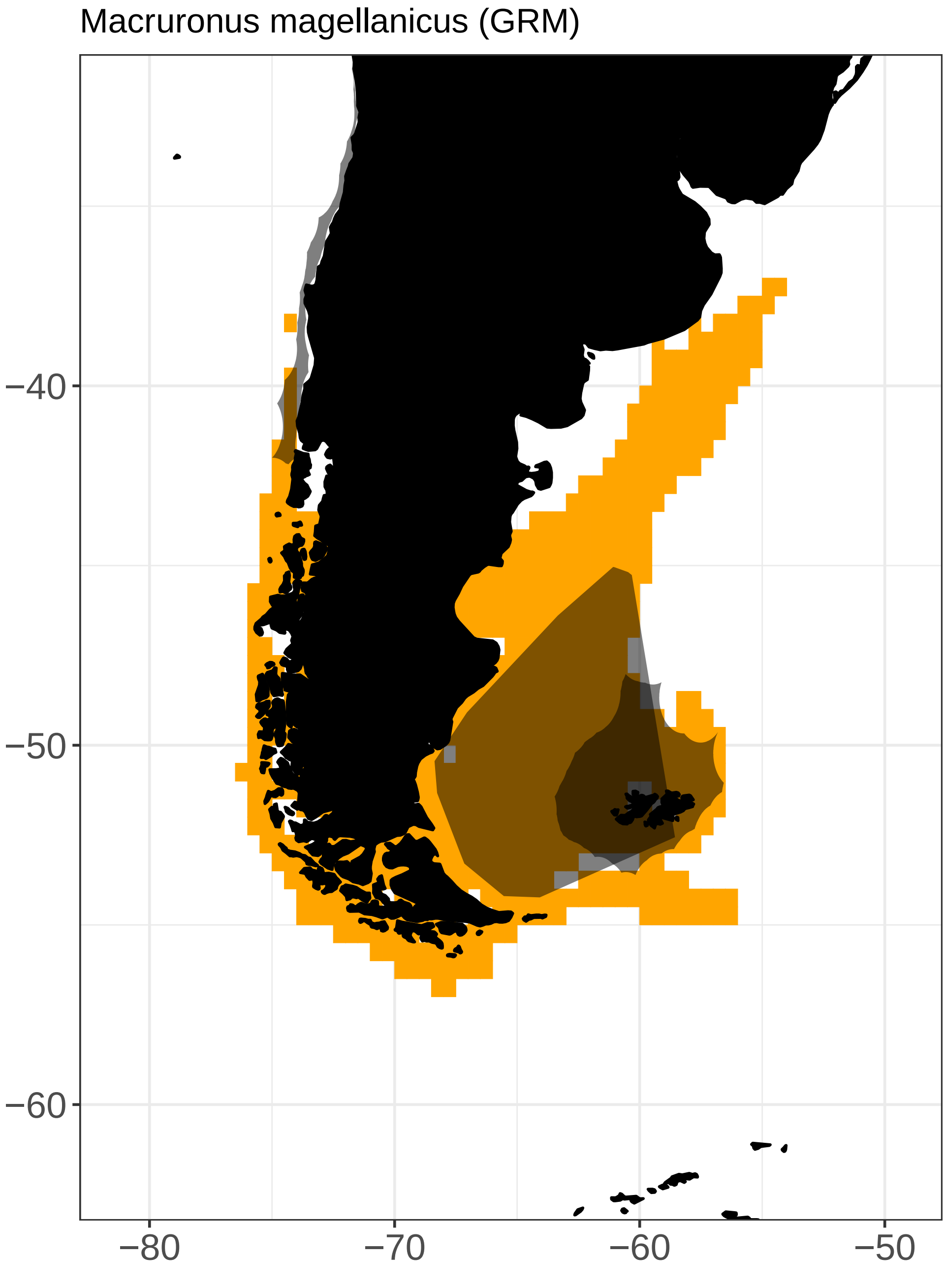

Macruronus magellanicus (GRM)
−40
−50
−60
−80
−70
−60
−50

Macruronus novaezelandiae (GRN)

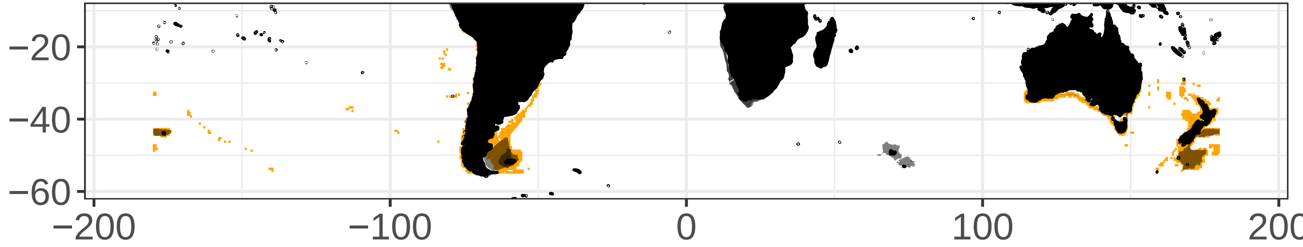

Melanogrammus aeglefinus (HAD)

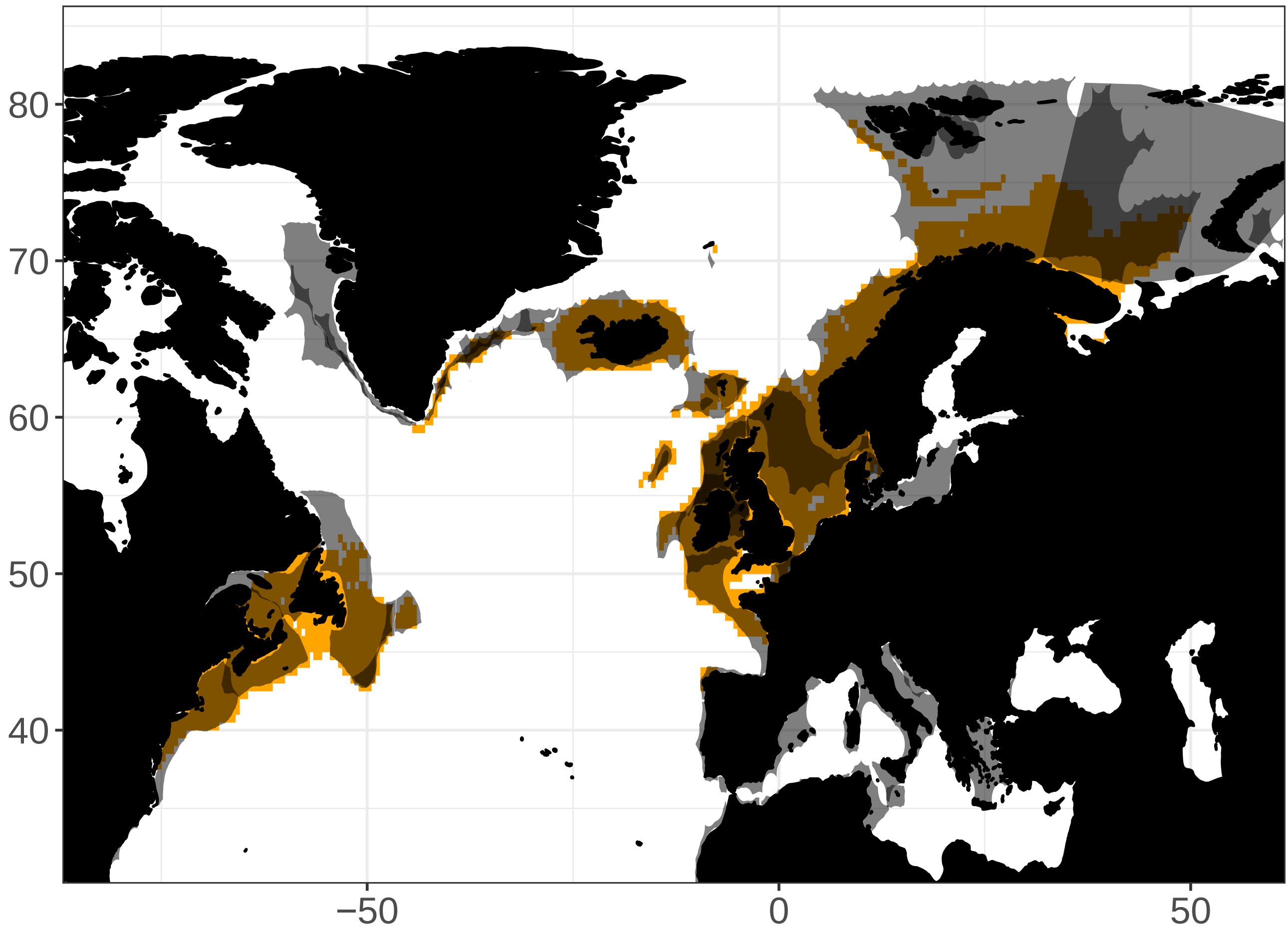

Mene maculata (MOO)

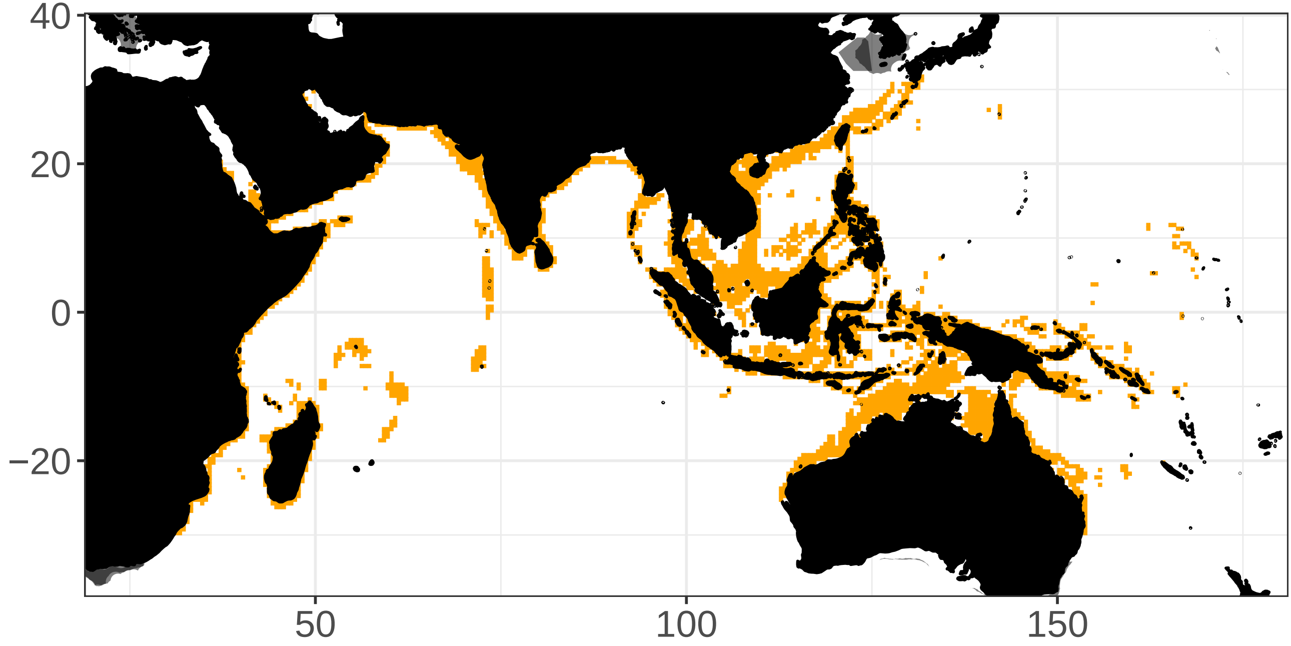

Merlangius merlangus (WHG)

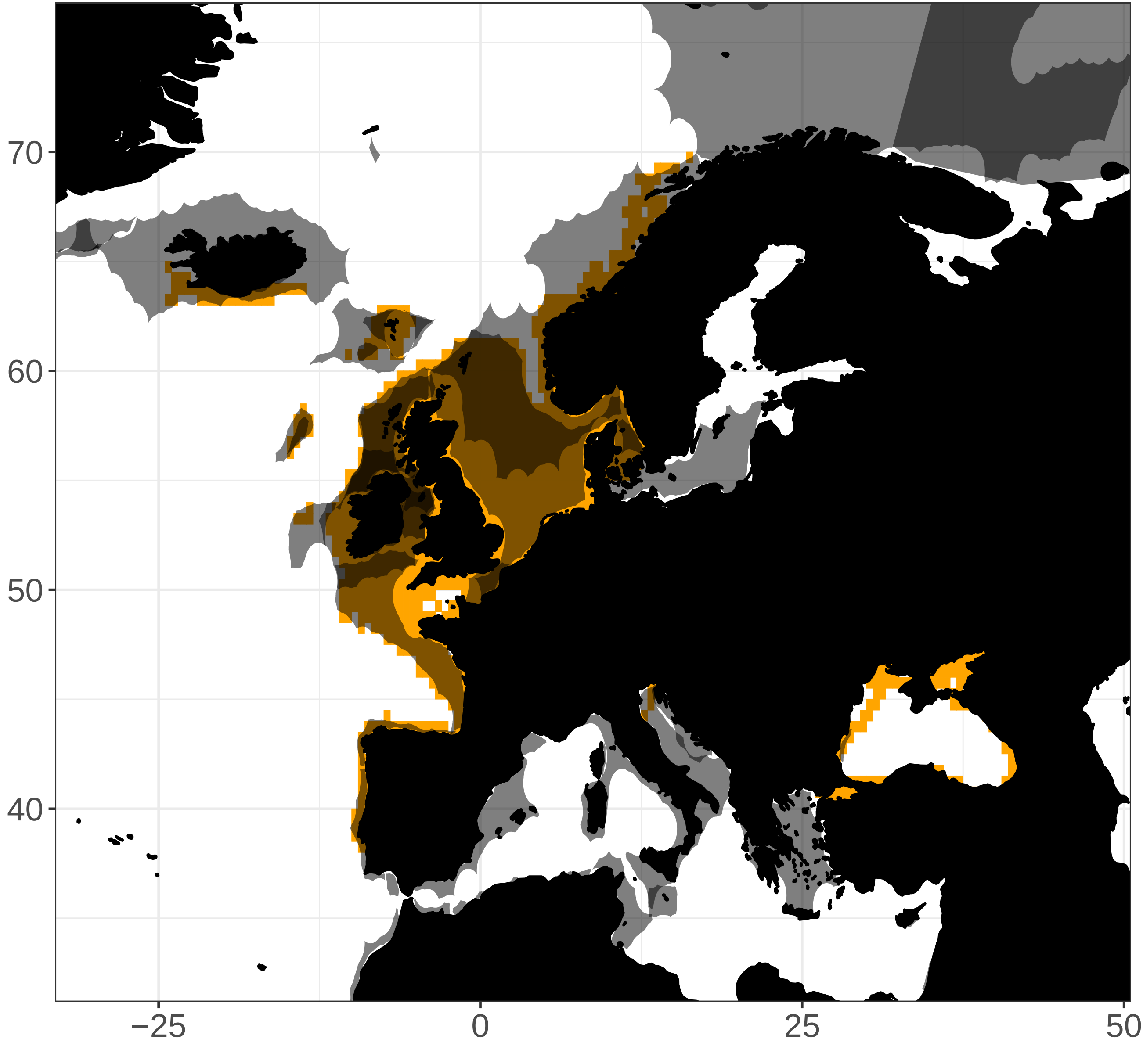

Merluccius capensis (HKK)

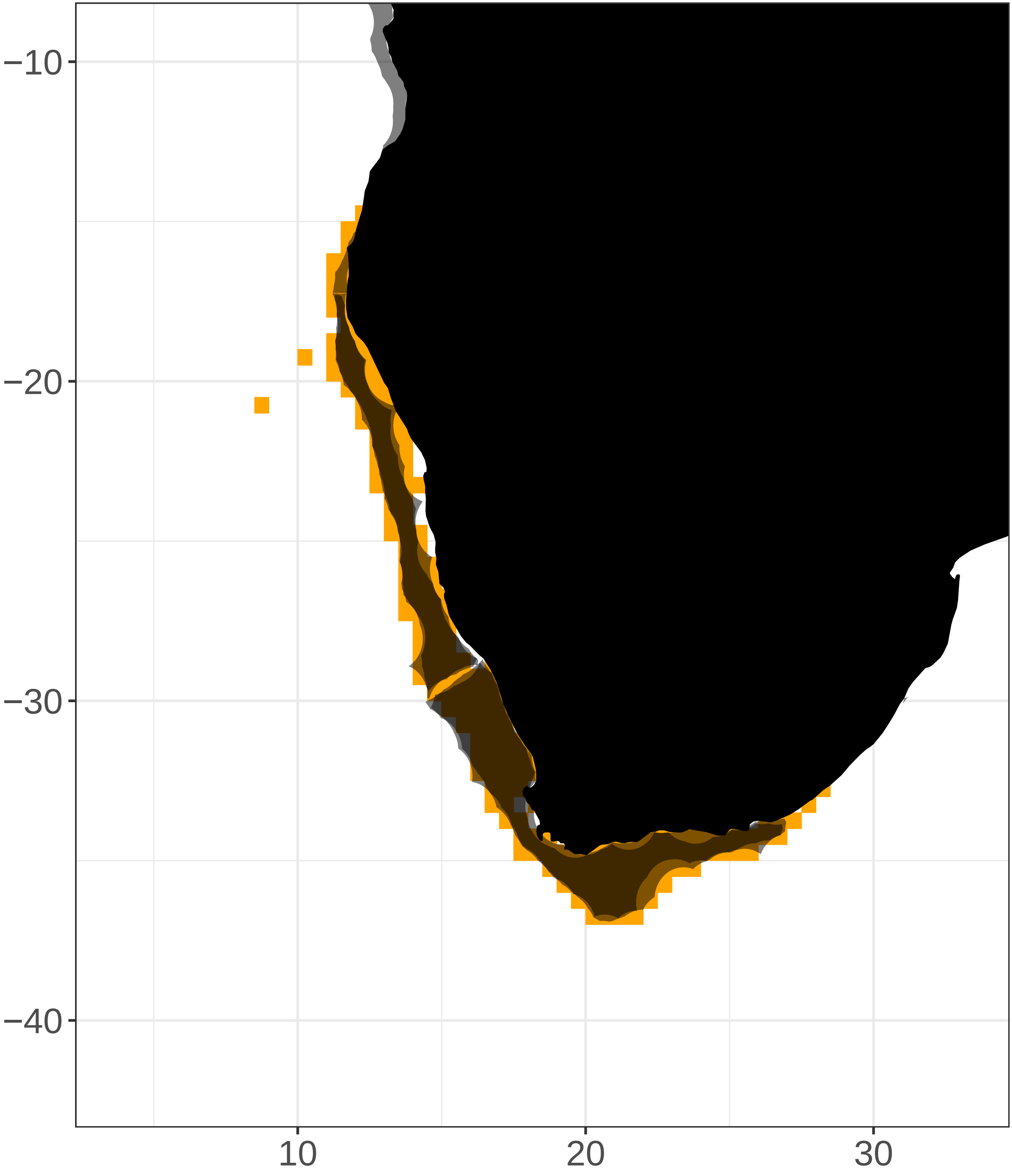

Merluccius gayi gayi (PHA)

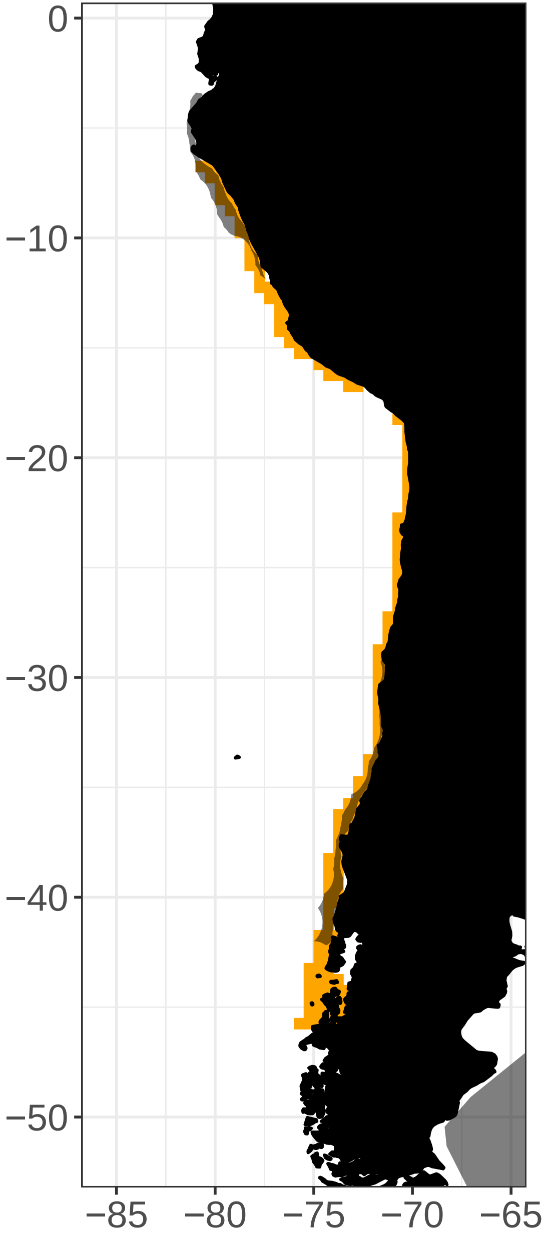

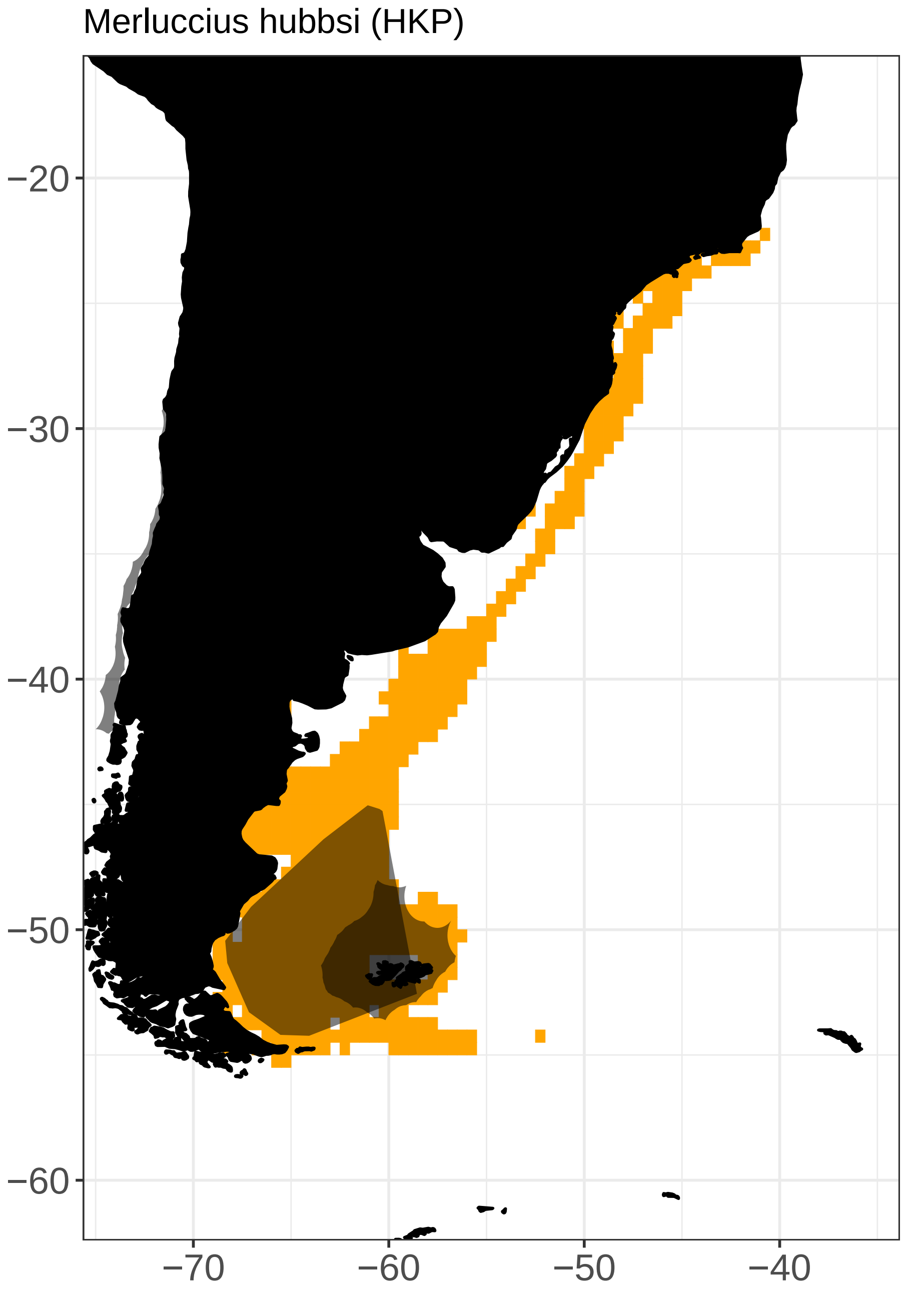

Merluccius hubbsi (HKP)
−20
−30
−40
−50
−60
−70
−60
−50
−40

Merluccius merluccius (HKE)

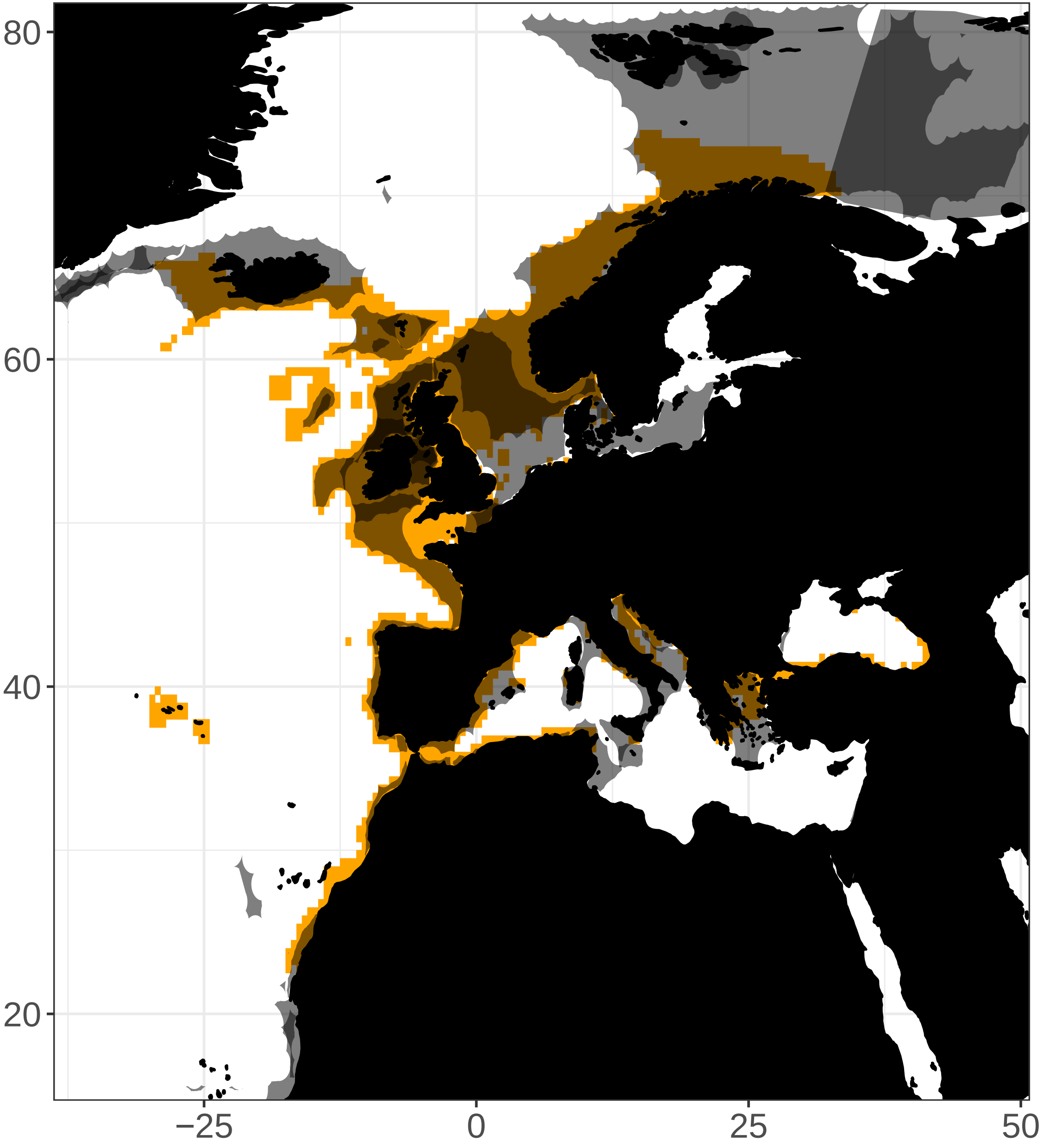

Merluccius productus (NHA)

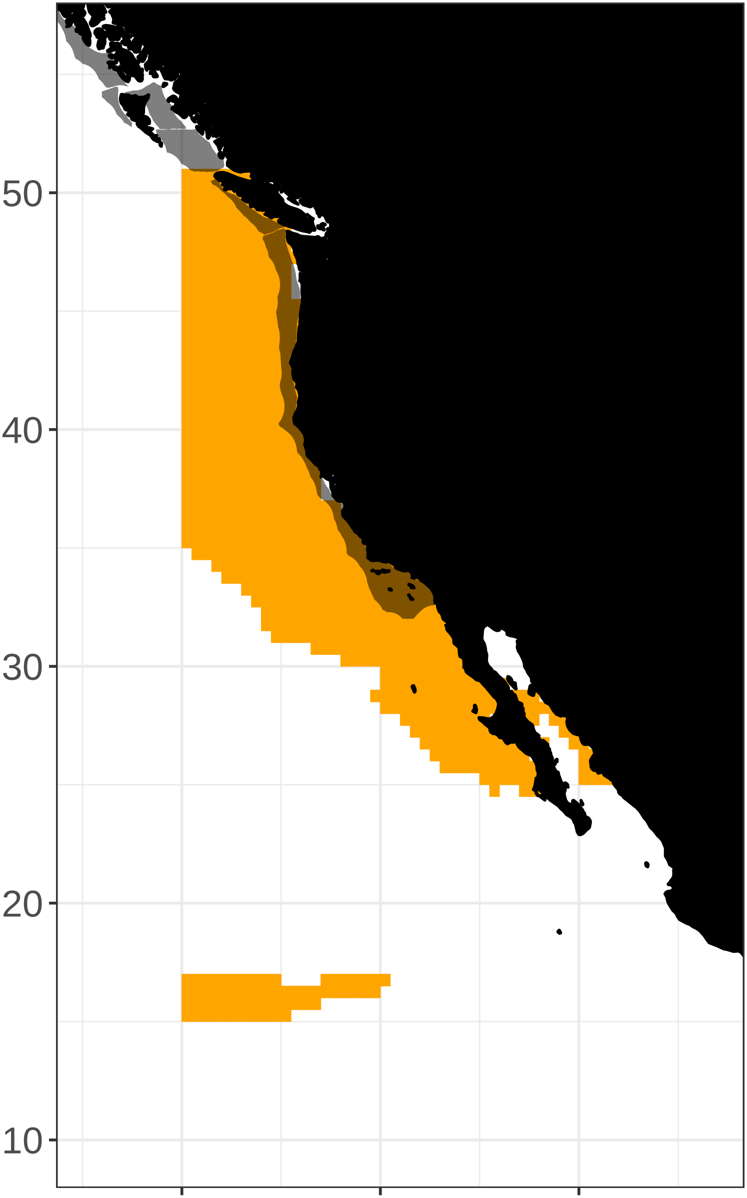

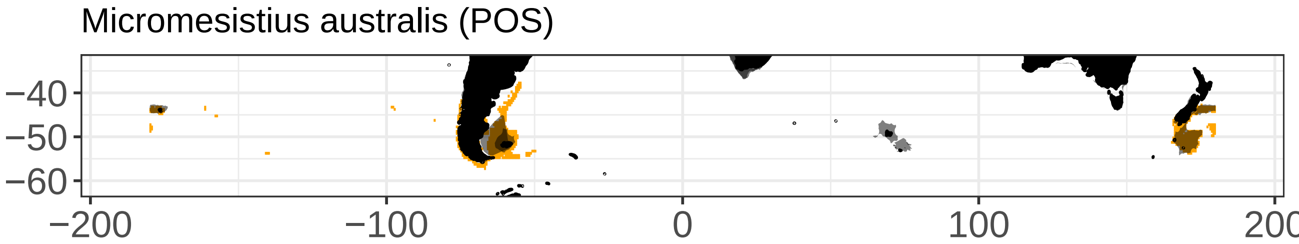

Micromesistius australis (POS)
−40
−50
−60
−200
−100
0
100
200

Micropogonias furnieri (CKM)

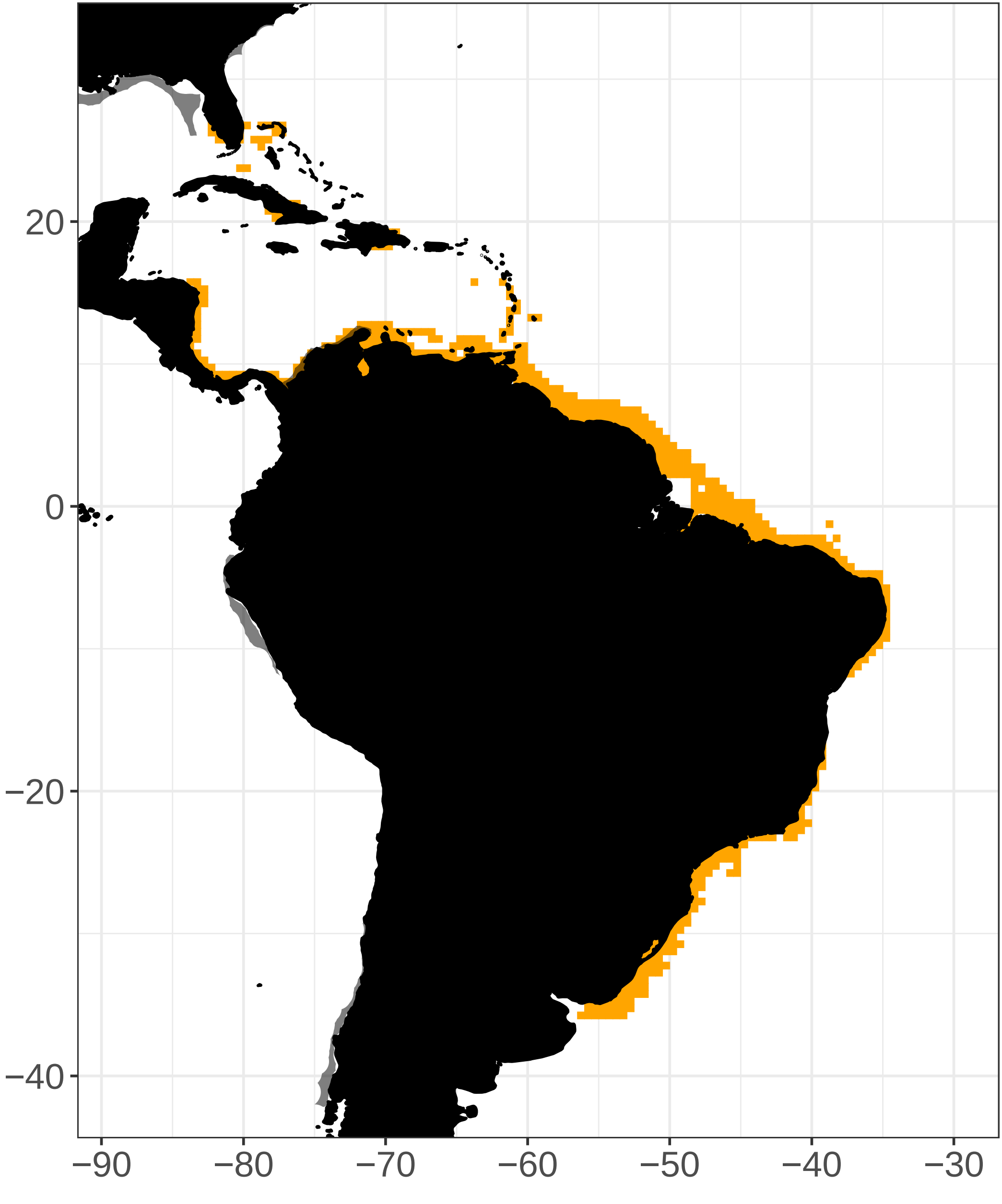

Mugil cephalus (MUF)

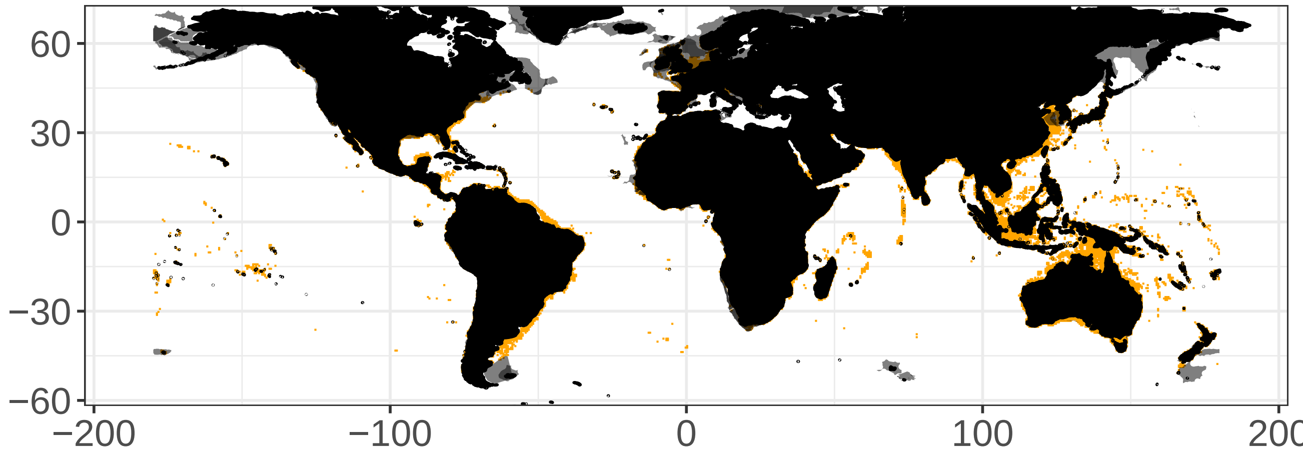

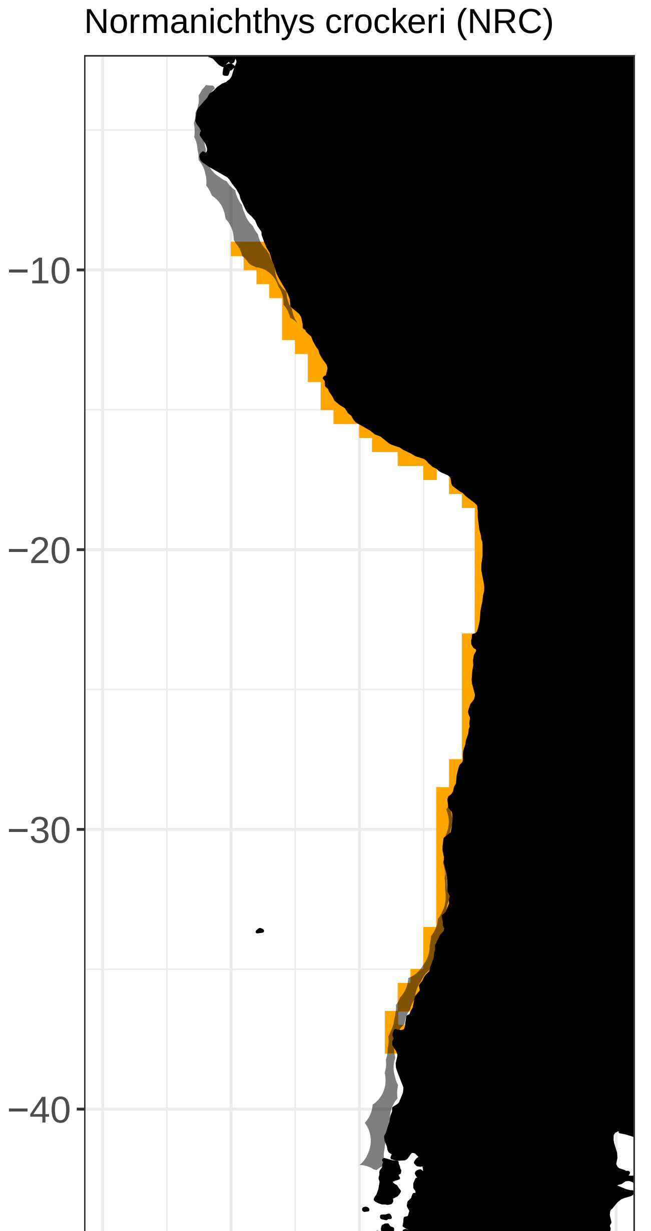

Normanichthys crockeri (NRC)
−10
−20
−30
−40
−85
−80
−75
−70
−65

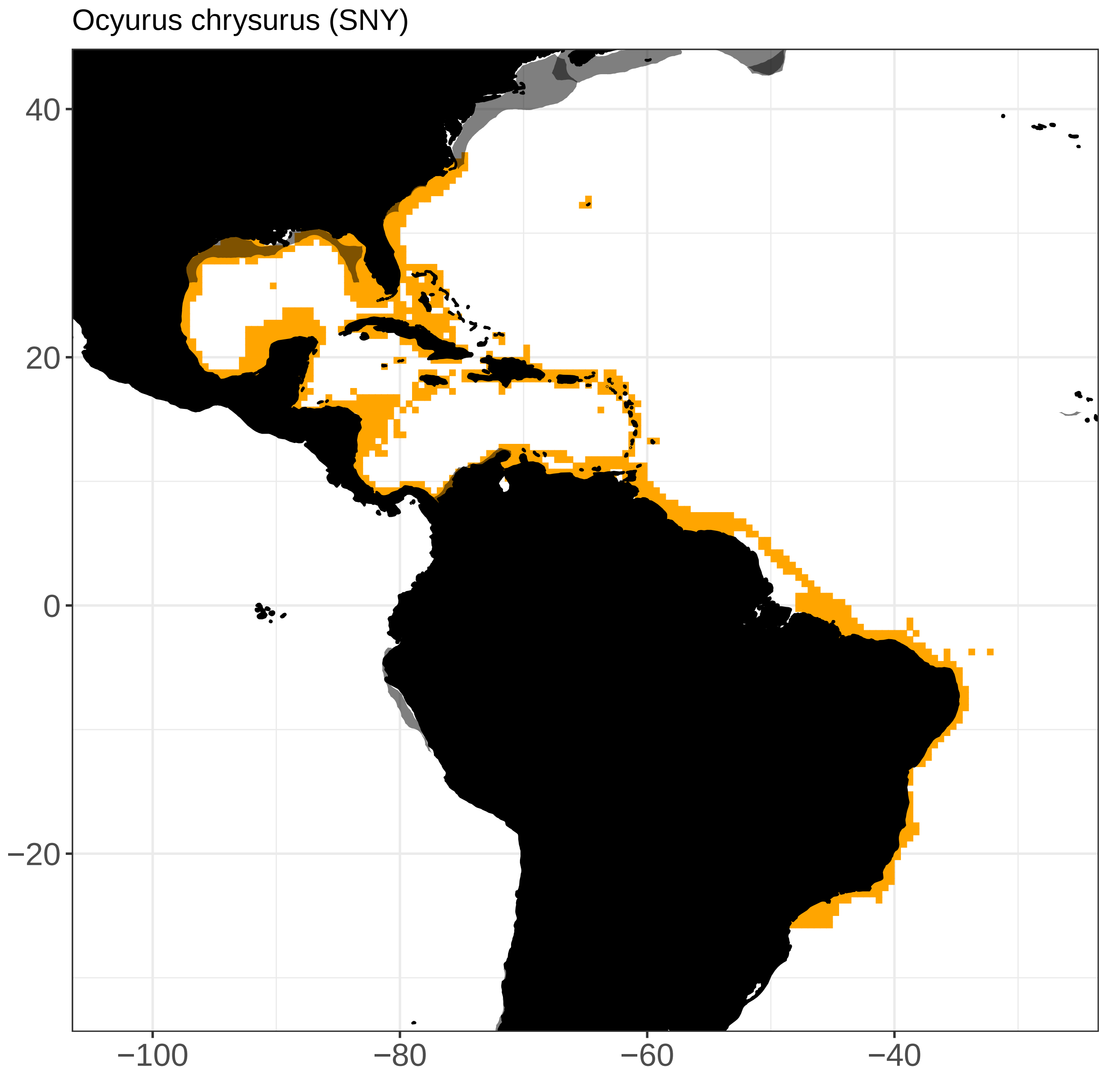
Ocyurus chrysurus (SNY)
40
20
0
−20
−100
−80
−60
−40

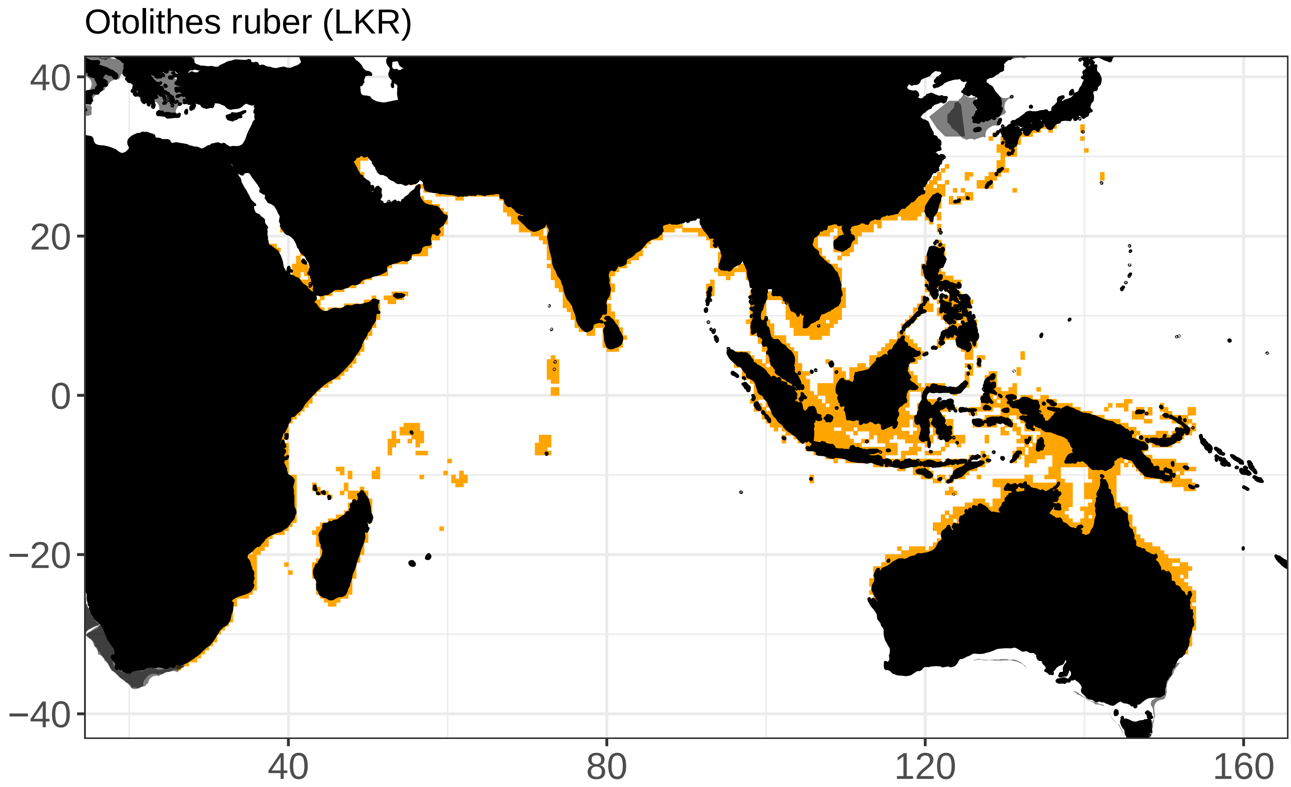
Otolithes ruber (LKR)
40
20
0
−20
−40
40
80
120
160

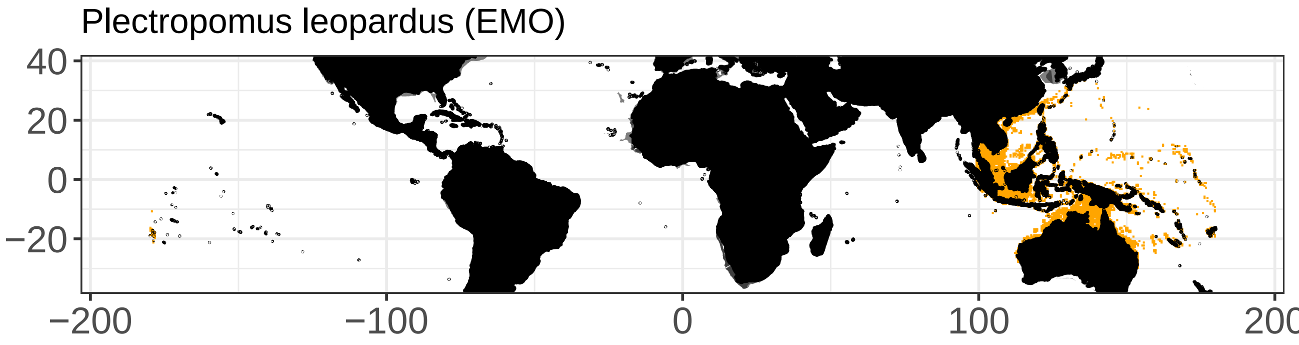

Plectropomus leopardus (EMO)
40
20
0
−20
−200
−100
0
100
200

Pollachius virens (POK)

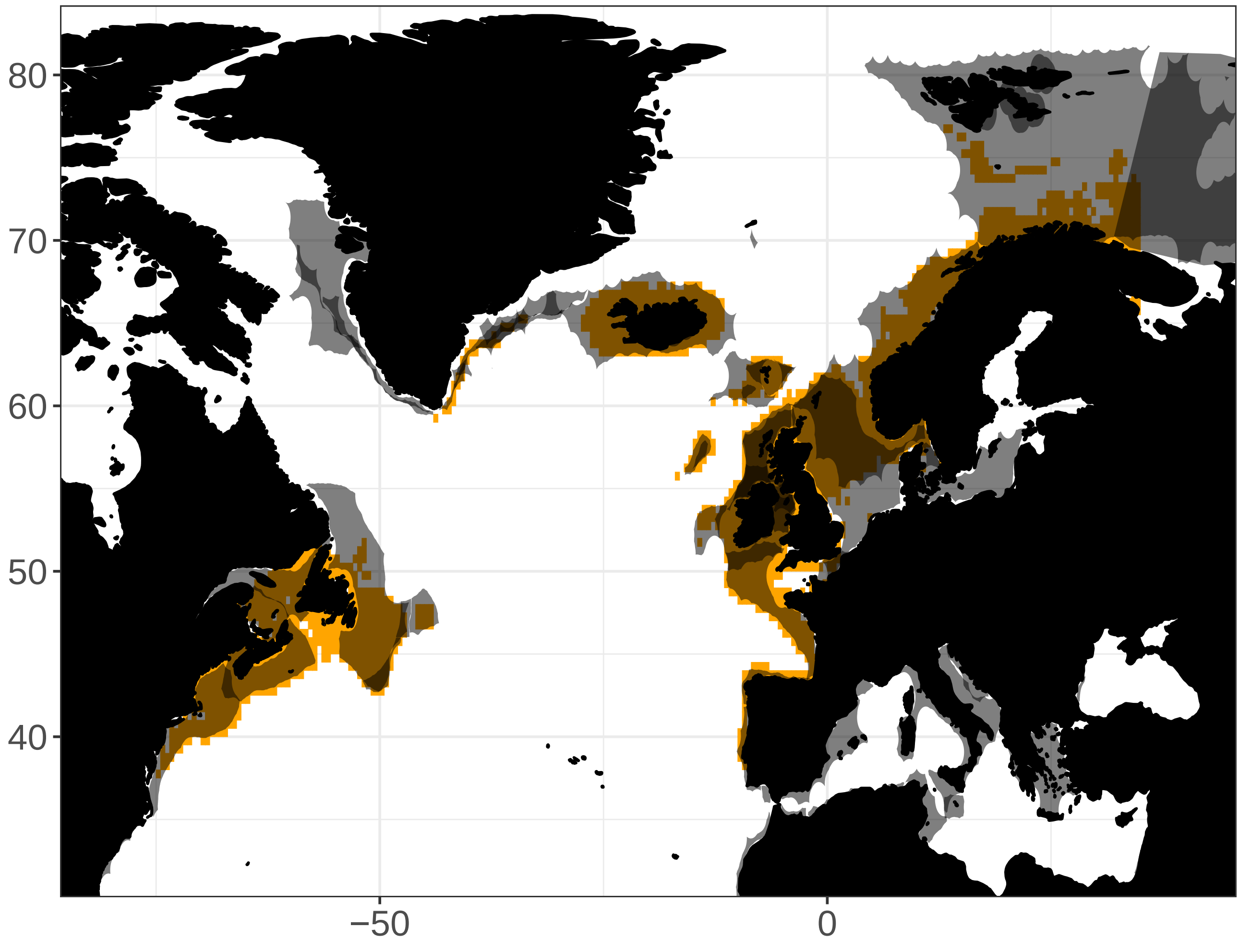

Polydactylus quadrifilis (TGA)

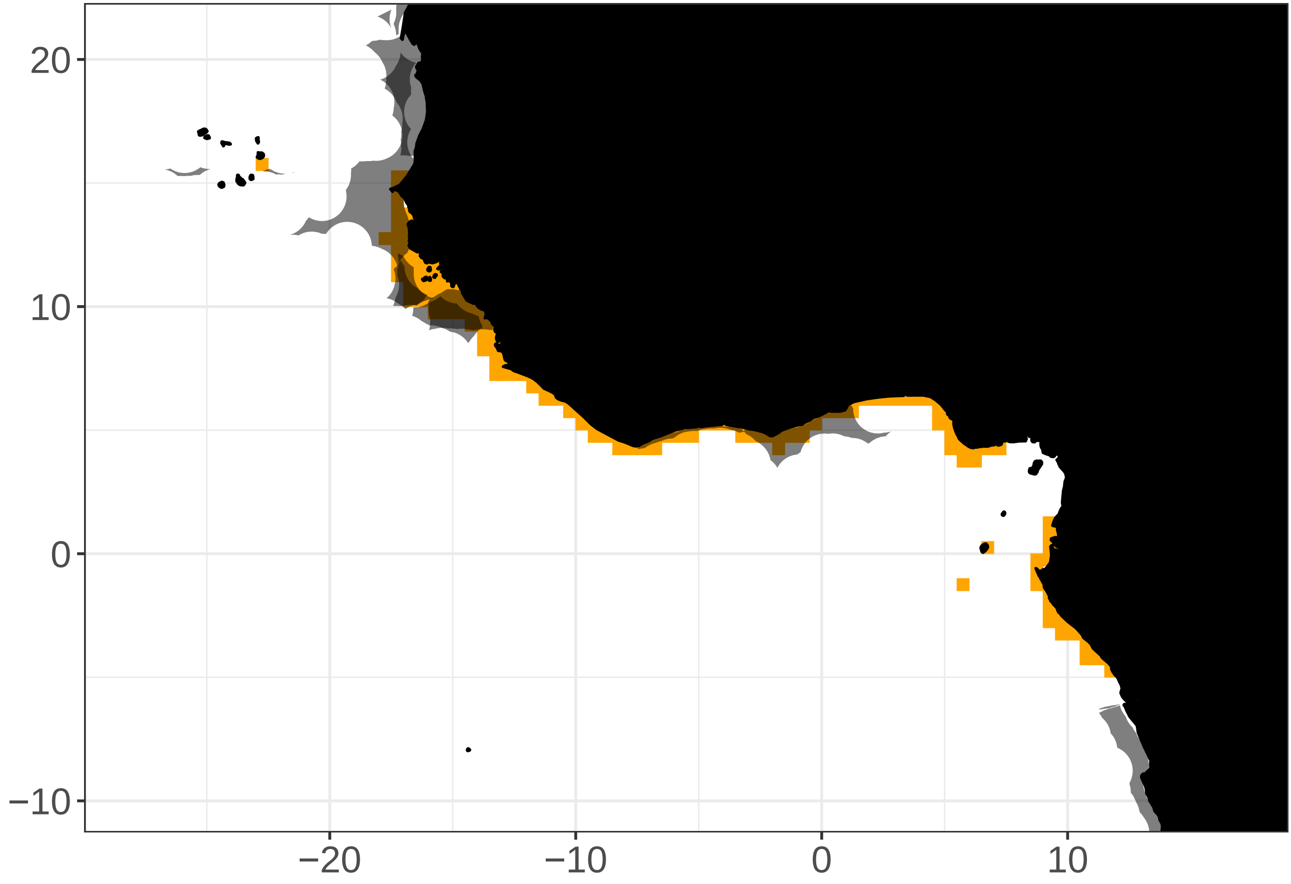

Reinhardtius hippoglossoides (GHL)

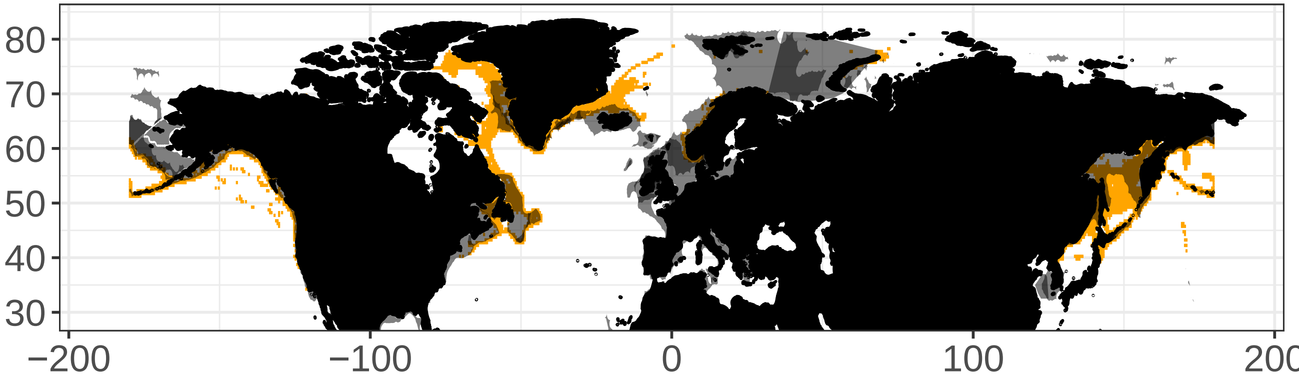

Thyrsites atun (SNK)

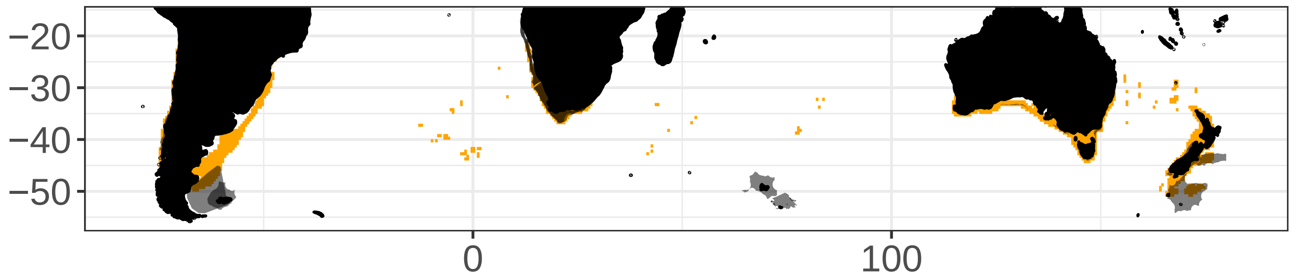

Trichiurus lepturus (LHT)

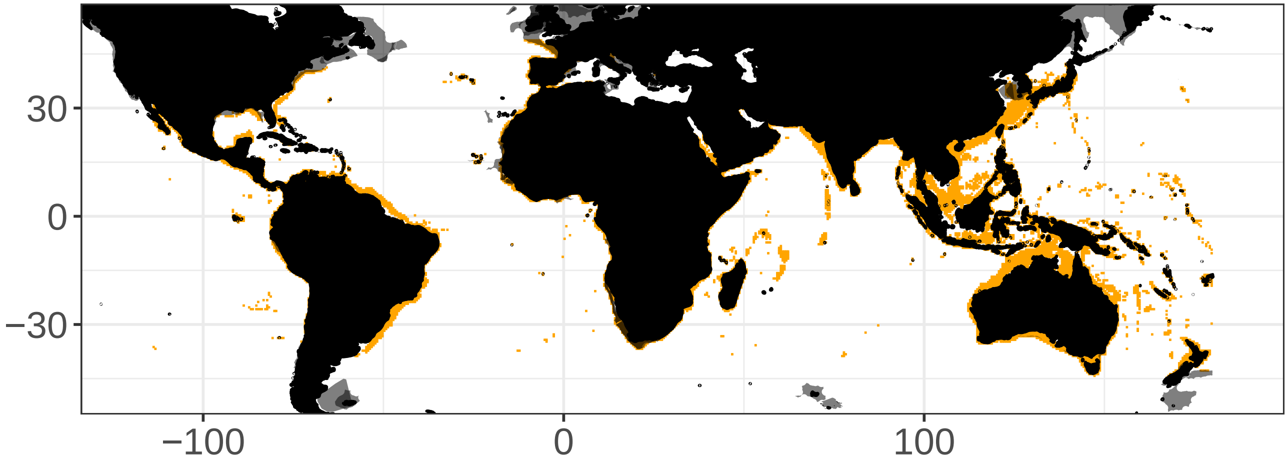
